# Glycosylation of *Plasmodium falciparum* TRAP supports sporozoite motility and invasion

**DOI:** 10.1101/2025.06.26.658380

**Authors:** Priya Gupta, Vladimir Vigdorovich, Nastaran Rezakhani, Lucia Pazzagli, Hardik Patel, Gigliola Zanghi, Mohd Kamil, Alexander Watson, Nelly Camargo, Erin Knutson, Robert L. Moritz, Stefan H. I. Kappe, D. Noah Sather, Ashley M. Vaughan, Kristian E. Swearingen

**Affiliations:** Center for Global Infectious Disease Research, Seattle Children’s Research Institute, Seattle, Washington, United States of America; Department of Pediatrics, University of Washington, Seattle, Washington, United States of America; Institute for Systems Biology, Seattle, Washington, United States of America

## Abstract

The human malaria parasite *Plasmodium falciparum* (*Pf*) expresses ten different thrombospondin type 1 repeat (TSR) domain-bearing proteins at different stages throughout its life cycle. TSRs can be modified by two types of glycosylation: O-fucosylation at conserved serine (S) or threonine (T) residues and C-mannosylation at conserved tryptophan (W) residues. *Pf*TRAP, which is expressed in mosquito-stage sporozoites, has one TSR domain that is O-fucosylated at Thr^256^ and C-mannosylated at Trp^250^. We employed site-directed mutagenesis by CRISPR/Cas9 gene editing to generate two *Pf*TRAP glyco-null mutant parasites, *Pf*TRAP_T256A and *Pf*TRAP_W250F, and assessed the fitness of these mutant parasites across the life cycle compared to the wild-type NF54 line as well as a *Pf*TRAP knockout line. The *Pf*TRAP glyco-null parasites exhibited major fitness defects comparable to knockout: sporozoites were unable to productively colonize the salivary glands and were highly impaired in gliding motility and the ability to invade cultured human hepatocytes. *Pf*TRAP abundance in these mutants was significantly decreased despite normal transcript levels. Biophysical assays with recombinant proteins confirmed that glycosylation of the *Pf*TRAP TSR stabilizes the domain and is likely required for its folding and secretion. These findings demonstrate that glycosylation of *Pf*TRAP’s TSR is critical for its proper expression and function, and underscore the importance of TSR glycosylation in the mosquito stage of the life cycle.

**IMPORTANCE:** Malaria is a mosquito-borne disease caused by *Plasmodium* parasites, of which *P. falciparum* is the deadliest. *Plasmodium* has ten proteins bearing thrombospondin type 1 repeats (TSRs), protein folds that aid cell-cell recognition and binding. Each of *Plasmodium*’s ten TSR-bearing proteins is important for invading tissues in the mosquito vector and human host. TSRs are decorated with sugar molecules, a modification termed glycosylation. To better understand the importance of TSR glycosylation in *Plasmodium*, we investigated the *P. falciparum* protein TRAP, which is only expressed in mosquito-stage parasite forms called sporozoites. When *Pf*TRAP was mutated to prevent glycosylation, abundance of the protein significantly decreased and parasites were unable to colonize the mosquito salivary glands. Furthermore, these mutant sporozoites were unable to move or to invade human liver cells. Our study reveals how TSR glycosylation can support the function of proteins that are required for parasite virulence.

## INTRODUCTION

Malaria is a major global health burden with an estimated 282 million cases and 610,000 deaths reported in 2024 [1]. It is caused by apicomplexan parasites of the genus *Plasmodium*, which complete their life cycles in vertebrate and arthropod hosts, with the most lethal malaria resulting from *P. falciparum* (*Pf*) infections [2]. The search for novel strategies to combat malaria includes identifying proteins that are essential to parasite virulence, especially those that are accessible to antibodies and can thus be the basis of subunit vaccines. A class of antigen that meets both of these criteria is the thrombospondin type 1 repeat (TSR), a small protein domain with a wide range of functions, including adhesion and cell-cell interactions [3]. Indeed, the two malaria vaccines recommended by the WHO, RTS,S/AS01 and R21/Matrix-M, include the TSR domain of the *Pf* protein CSP, and the vaccine candidate ME-TRAP is based on the *Pf* protein TRAP, which also contains a TSR domain [4]. The *Plasmodium* genome encodes a total of ten different TSR-bearing proteins that are expressed at various stages throughout the parasite life cycle (**Fig 1a**), and disrupting their expression or function can severely impair parasite development [5–14].

**Figure 1.**
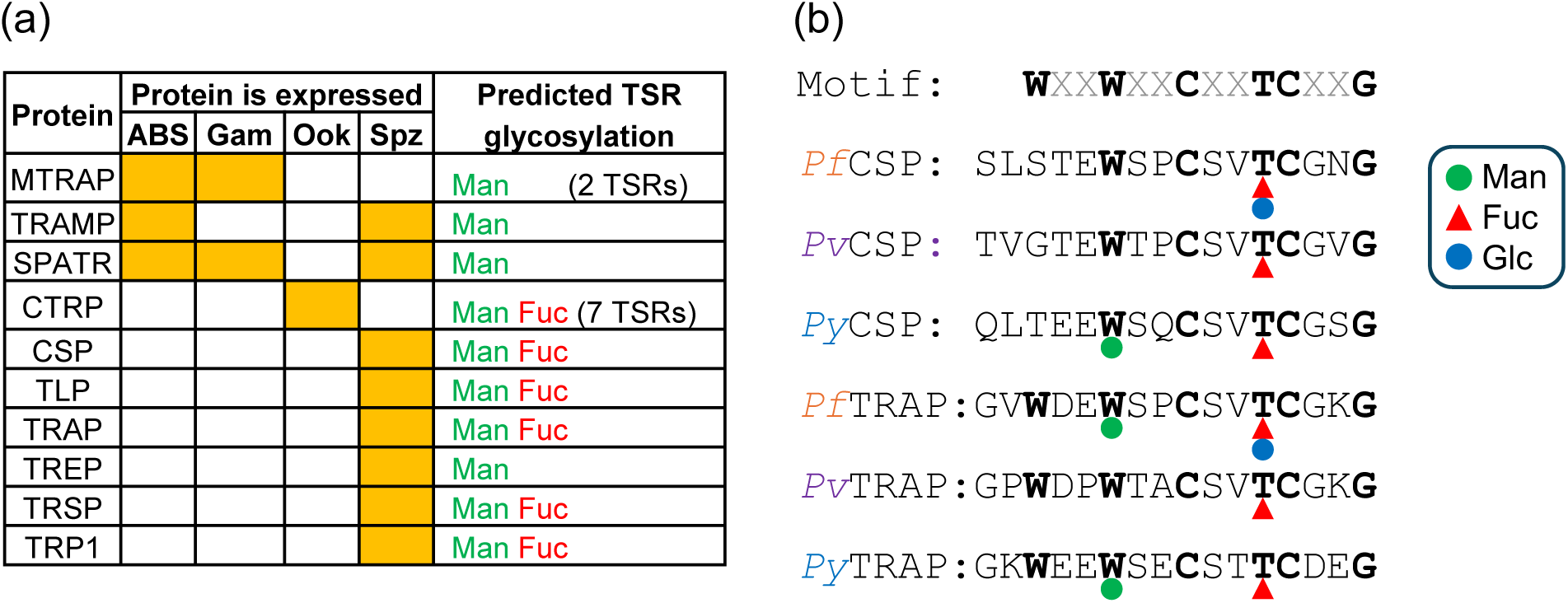
*Plasmodium* proteins with thrombospondin type 1 repeats (TSRs). (a) The ten TSR-bearing proteins encoded by the *Plasmodium* genome. Orange squares indicate that the protein is known to be expressed in the given stage based on published studies. ABS: asexual blood stages; Gam: gametocytes; Ook: ookinetes; Spz: sporozoites. The presence of motifs predicted or known to be glycosylated is indicated. Man: C-mannosylation of Trp; Fuc: O-fucosylation of Ser or Thr. (b) Glycosylation of CSP and TRAP observed in published studies of salivary gland sporozoites analyzed by mass spectrometry. *Pf*: *P. falciparum; Pv*: *P. vivax; Py*: *P. yoelii.* Green circle: C-mannose; red triangle: O-fucose; blue circle: β-1,3-glucose.

Although TSR domains contain few secondary structure elements, their characteristic fold is stabilized by a network of conserved disulfide bonds, and the domain is glycosylated in a highly specific manner [15–18]. The C-mannosyltransferase DPY19 recognizes a conserved primary sequence motif in the domain [19] and adds a C-linked mannose (C-Man) to conserved tryptophan (Trp or W) residues [18]. The O-fucosyltransferase POFUT2 only recognizes correctly folded TSRs [20] and modifies conserved serine (Ser or S) and threonine (Thr or T) residues with an O-linked fucose (O-Fuc) [16] that can be further extended with a β-1,3-linked glucose (Glc) by the glucosyltransferase B3GLCT, resulting in an O-Fuc-Glc disaccharide [21]. In addition to altering the physicochemical properties of TSR domains, these glycans restrict the domains’ conformational flexibility and provide a selection mechanism for correct conformations and disulfide linkage patterns [22, 23]. Preventing either O-fucosylation or C-mannosylation of TSRs by site-directed mutagenesis of glycosites [24] or by knockout of glycosyltransferases [25, 26] can interfere with protein folding and secretion, but the extent of these defects ranges from non-existent to severe for different proteins in an organism and even for different glycosites within the same protein.

*Plasmodium* TSR domains are glycosylated akin to their better-studied counterparts [27–29], and the *Plasmodium* POFUT2 [30] and DPY19 [31] have been characterized. A *Plasmodium* B3GLCT has yet to be identified, but extension of O-Fuc with a hexose to form a disaccharide has been observed in *Pf* [27], thus it is assumed that a *Pf*B3GLCT exists and that the disaccharide is O-Fuc-Glc. Although the glycosites in *Plasmodium* TSR domains are highly conserved, their modification is species- and protein-specific [27–29] (**Fig 1b**). For example, the essential sporozoite invasin TRAP contains a TSR domain that is modified with C-Man at a conserved Trp residue in the rodent parasite *P. yoelii* (*Py*) and in the human parasite *Pf,* but not in the human parasite *P. vivax* (*Pv*). Furthermore, although TRAP in all three species is modified with O-Fuc at a conserved Thr residue, this O-Fuc is extended to the O-Fuc-Glc disaccharide only in *Pf*.

The role of TSR glycosylation in *Plasmodium* biology is an area of active study, and there is seemingly contradictory data from work performed by different laboratories. Lopaticki *et al*. described a *Pf*POFUT2 knockout mutant that exhibited a significant decrease in the number of oocysts that developed in the midguts of infected mosquitoes [30], though a *Pf*POFUT2 knockout produced by Sanz *et al.* developed oocysts comparable to wild type [32]. In the *Pf*POFUT2 knockout of Lopaticki *et al*. as well as a *P.berghei* (*Pb*) POFUT2 knockout by Srivastava *et al.* [33], sporozoites isolated from mosquito salivary glands had reduced gliding motility and reduced ability to invade liver cells *in vitro* and *in vivo*, while a *Pb*POFUT2 knockout by Sanz *et al*. had no discernible defects in sporozoites’ ability to infect mice via mosquito bite [32]. In *Pf* and *Pb*POFUT2 knockout sporozoites, CSP expression was unaffected but TRAP levels were significantly reduced in the absence of O-fucosylation [30, 33]. The effect on the other three putatively O-fucosylated proteins expressed in sporozoites – TRSP, TLP, and TRP1 – was not investigated. It therefore remains to be determined which TSR-bearing protein or proteins are responsible for the observed phenotypes and how O-fucosylation supports their structure and function. Similarly, C-mannosylation is important for parasite fitness, but which proteins require the modification is largely unexplored. López-Gutiérrez *et al.* [34] and Lopaticki *et al.* [35] found that *Pf*DPY19 was dispensable for asexual blood stage growth, and Srivastava *et al*. reported the same for *Pb*DPY19 [36] despite the fact that three putatively C-mannosylated proteins – SPATR, MTRAP, and TRAMP – are expressed in these stages. SPATR and MTRAP levels were decreased in the absence of C-mannosylation but TRAMP was unaffected [35]. Lack of DPY19 led to a complete absence of sporozoites in both *Pb* and *Pf,* correlating with reduced gametocyte numbers, impaired fertilization [35], and total loss of gliding motility in ookinetes [36]. This latter phenotype points to CTRP requiring C-mannosylation, but additional work is needed to parse how each of these proteins contribute to the observed fitness defects. Furthermore, the fact that DPY19-null parasites do not produce sporozoites means that knockout lines such as these cannot be used to investigate the importance of C-mannosylation for the six TSR-bearing proteins expressed in sporozoites, all of which have putative or known C-mannosylation sites. It is also important to note that none of the above studies used mass spectrometry (MS) to directly confirm that their knockouts had actually prevented glycosylation of the proteins in question.

Taken together, these studies show that C-mannosylation and O-fucosylation of TSRs support parasite virulence, but the importance and function of each modification for each *Plasmodium* TSR-bearing protein remains largely unknown. Moreover, the evidence underscores that the role of glycosylation in the function of *Plasmodium’s* TSR-bearing proteins cannot be generalized but must be considered on a case-by-case basis for each glycosite of each protein in each species. Here we demonstrate the first comprehensive characterization of the role of O-fucosylation and C-mannosylation in the function of a *Plasmodium* TSR-bearing protein, the essential mosquito-stage invasin *Pf*TRAP. TRAP belongs to a superfamily of adhesins that have roles in motility and invasion [37, 38]. It is stored in secretory organelles known as micronemes [39] until released to the sporozoite surface upon activation [40, 41], whereupon it produces adhesion sites that are linked to the actin-myosin motility machinery [42]. TRAP is essential for sporozoite invasion of salivary glands (mediated in part by its TSR domain) and invasion of the liver [43, 44]. Previous work suggests that O-fucosylation of its TSR is critical for its expression [30, 33], but the role of C-mannosylation has not been explored. We have generated transgenic parasites that bear conservative single-residue substitutions in *Pf*TRAP to prevent either O-fucosylation or C-mannosylation and confirmed the complete absence of these modifications in the glyco-null mutants by MS-based proteomics. The parasites developed normally in blood stages and were able to infect mosquitoes, but sporozoites lacked gliding motility and were severely impaired in their ability to colonize salivary glands and invade hepatocytes, demonstrating that both O-fucosylation and C-mannosylation of *Pf*TRAP are critical for parasite fitness.

## RESULTS

### *Pf*TRAP glyco-null mutants exhibit normal blood stage development and transmission to mosquitoes

Previous work has shown that *Pf*TRAP is glycosylated with O-Fuc at Thr^256^ and C-Man at Trp^250^ [27]. To examine the role of TSR glycosylation in the function of *Pf*TRAP, we employed CRISPR/Cas9-mediated gene editing of the wild-type *Pf* NF54 line to generate two parasite lines with substitutions that prevent modification of these two residues, *Pf*TRAP_T256A and *Pf*TRAP_W250F. We selected amino acid substitutions with the least likelihood of disrupting the protein structure: O-fucosylation was prevented by replacing Thr^256^ with alanine (Ala or A) and C-mannosylation was prevented by replacing Trp^250^ with phenylalanine (Phe or F) [45]. We also generated a gene deletion parasite, *Pf*TRAP^—^ (**Fig S1, Fig S2, Table S1**). The knockout and glyco-null lines developed normally and exhibited no defects in the ability to form sexual stage gametocytes and gametes (**Fig S3, Table S2**). When mature gametocytes were fed to female *Anopheles stephensi* mosquitoes by membrane feeding [46], there were no significant differences in the number or prevalence of oocysts or in the number of sporozoites in the midguts (**Fig 2a-c, Table S2**).

**Figure 2.**
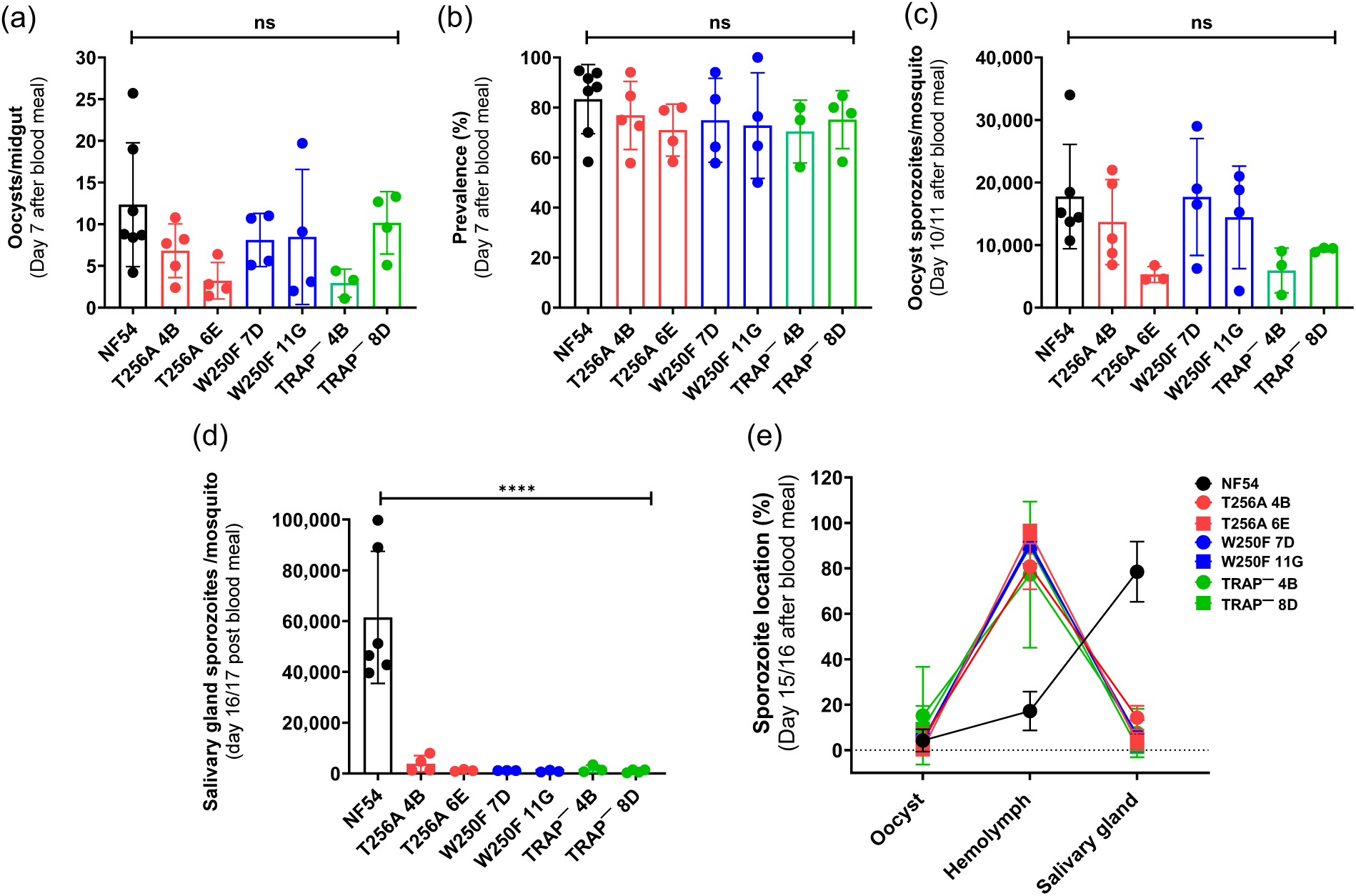
*Pf*TRAP glyco-null mutant sporozoites are deficient in colonization of mosquito salivary glands. The fitness of the *Pf*TRAP_T256A O-fucosylation-null mutant (clones 4B and 6E), *Pf*TRAP_W250F C-mannosylation null mutant (clones 11G and 7D), and *Pf*TRAP^—^ gene deletion mutant (clones 4B and 8D) parasite lines was compared to the wild-type NF54 line. There was no significant difference between any of the transgenic lines and the wild type for (a) the mean number of oocysts per mosquito, (b) oocyst prevalence (the percentage of mosquitoes with oocyst in midguts) 7 days post-feeding, or (c) the mean number of oocyst sporozoites per mosquito on day 10 or 11 post-feeding. (d) The mean number of salivary gland sporozoites per mosquito on day 16 or 17 post-feeding was significantly reduced for the glyco-null and knockout mutants compared to wild type (**** *p* < 0.0001). Each point represents an independent experiment. Bars are mean ±S.D. (e) Sporozoites were collected from oocysts, hemolymph, and salivary glands of the same mosquitoes on day 15 or 16 post-feeding. The relative proportion of sporozoites found in each compartment is shown as percentage. Data is mean ±S.D. of independent experiments. Source data are given in **Table S2**.

### *Pf*TRAP glyco-null mutant sporozoites egress from midguts but fail to colonize salivary glands

There was a significant (> 10-fold) reduction in the number of *Pf*TRAP^—^ sporozoites associated with salivary glands, consistent with the phenotype previously reported for a *Pb*TRAP^—^ line [7], and the glyco-null parasite numbers were comparable to those of the knockout parasite (**Fig 2d, Table S2**), demonstrating the importance of *Pf*TRAP expression and its glycosylation in salivary gland colonization by sporozoites. In order to determine whether the mutant sporozoites were unable to egress from the midguts or unable to invade the salivary glands, we dissected infected mosquitoes on day 15 or 16 post-feeding and isolated the oocyst, hemolymph, and salivary gland sporozoites from the same mosquitoes. In mosquitoes infected with wild-type parasites, the majority of sporozoites were associated with salivary glands, with significantly fewer found in the hemolymph, and very few still found in oocysts. By contrast, the glyco-null and *Pf*TRAP^—^sporozoites were predominantly found in the hemolymph (**Fig 2e, Fig S4, Table S2**). We therefore infer that *Pf*TRAP is dispensable for sporozoite egress from oocysts but essential for colonization of the salivary glands of the mosquito, and that preventing C-mannosylation or O-fucosylation of *Pf*TRAP functionally mimics deletion of the gene.

In order to determine whether the very few glyco-null and knockout sporozoites recovered from salivary glands had successfully invaded or were merely attached to the surface of the glands, we treated dissected salivary glands with trypsin to release any parasites adhered to the surface prior to grinding the tissue to release invaded sporozoites for counting. In mosquitoes infected with the wild-type parasite, 49% of sporozoites remained associated with salivary glands after trypsinization, indicating that they had invaded the tissue. By contrast, of the very few sporozoites associated with the salivary glands for the glyco-null and knockout parasites, only 12% of *Pf*TRAP_T256A, 14% of *Pf*TRAP_W250F, and 4% of *Pf*TRAP^—^ sporozoites still remained after trypsinization (**Fig S5**, **Table S2**) indicating that few, if any, sporozoites from glyco-null or knockout parasites had successfully invaded the salivary glands.

### Mass spectrometry reveals the glycosylation state of *Pf*TRAP in the glyco-null mutants

The predicted glycosites of *Pf*TRAP are contained on a single peptide produced by trypsin cleavage, enabling convenient detection by MS. To determine the stoichiometry of the glycoforms present, we analyzed oocyst sporozoites with a highly sensitive selected ion monitoring (SIM) MS approach targeting masses corresponding to possible permutations of the TSR glycopeptide (**Fig 3, File S1**). *Pf*TRAP from wild-type oocyst sporozoites exhibited the same glycosylation pattern as previously observed in salivary gland sporozoites: 100% of detected peptide was C-mannosylated at Trp^250^ and O-fucosylated at Thr^256^, with a portion of the O-Fuc extended to the O-Fuc-Glc disaccharide. As predicted, *Pf*TRAP_W250F sporozoites showed complete absence of C-mannosylation, yet 100% of the detected TSR peptide was still O-fucosylated. Conversely, no O-fucosylation was observed in the *Pf*TRAP_T256A sporozoites, while 100% of Trp^250^ was C-mannosylated. Unexpectedly, ∼9% of the detected TSR peptide was also C-mannosylated at Trp^247^. These results demonstrate that the *Pf*TRAP_T256A and *Pf*TRAP_W250F glyco-null mutant parasites prevent site-specific glycosylation of the *Pf*TRAP TSR, though the T256A mutation correlated with a small increase in off-target C-mannosylation.

**Figure 3.**
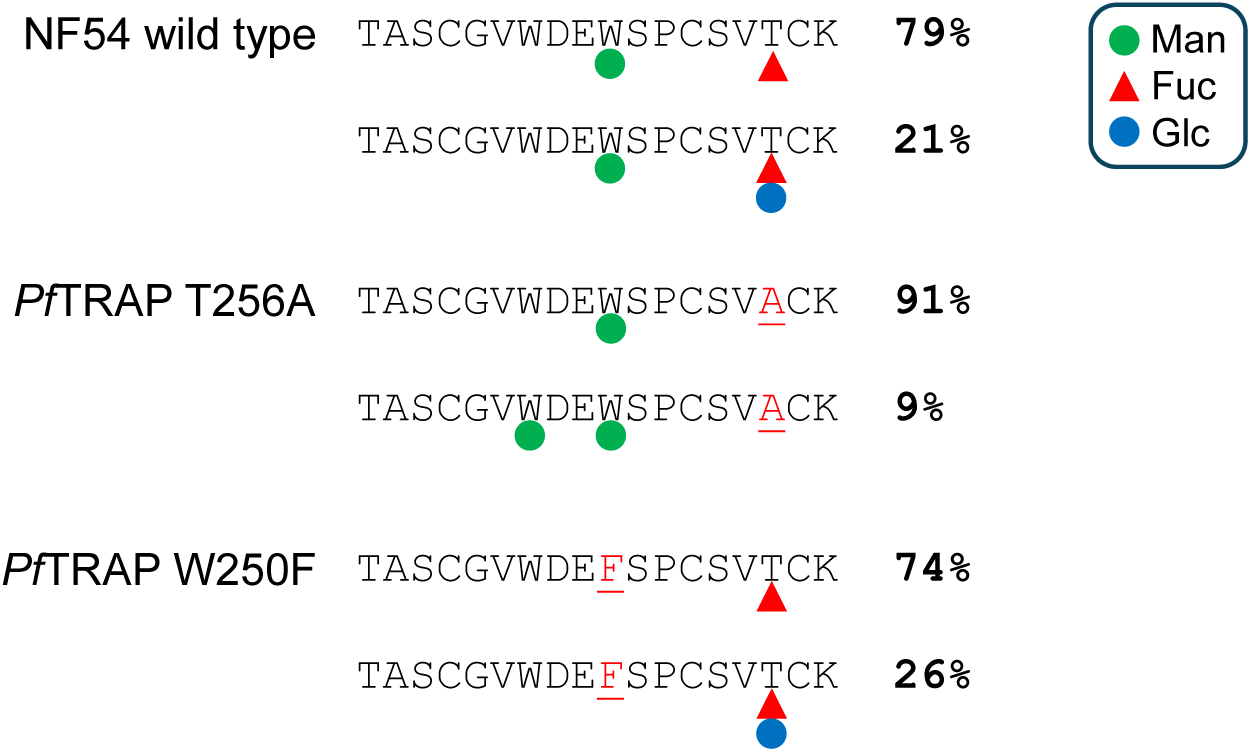
Glycosylation of *Pf*TRAP in wild-type and glyco-null sporozoites. The relative abundance of glycoforms of *Pf*TRAP in oocyst sporozoites was determined by mass spectrometry. Data are from one replicate each of wild-type NF54, *Pf*TRAP_T256A clone 4B, and *Pf*TRAP_W250F clone 11G. The sequence of the tryptic peptide bearing both glycosites is given, and the sites of substitution in the glyco-null mutants are indicated in red. The relative abundance of the detected glycoforms within each sample (determined by the chromatographic peak height) is given. Green circle: C-mannose; red triangle: O-fucose; blue circle: β-1,3-glucose. Chromatograms and annotated mass spectra are given in **File S1**.

### *Pf*TRAP glyco-null mutants exhibit decreased levels of *Pf*TRAP in sporozoites

We sought to understand whether the loss-of-function phenotype observed in the glyco-null mutants was caused by destabilization or mis-localization of *Pf*TRAP. To this end, we performed immunofluorescence microscopy using polyclonal antibodies raised against recombinant *Pf*TRAP [47]. As expected, *Pf*TRAP in wild-type sporozoites co-localized with *Pf*AMA1, which is also found in micronemes [48], and no *Pf*TRAP was detected in *Pf*TRAP^—^ parasites (**Fig 4a, Fig S6**). *Pf*TRAP localization was not altered in the glyco-null mutants, but protein abundance was significantly reduced as quantified by fluorescence intensity (**Fig 4b**). *Pf*TRAP mRNA abundance was not reduced in the glyco-null mutants compared to wild type (**Fig 4c**), indicating that the reduction in *Pf*TRAP levels was due to adverse effects on the *in vivo* stability of the protein caused by disrupting glycosylation of its TSR.

**Figure 4.**
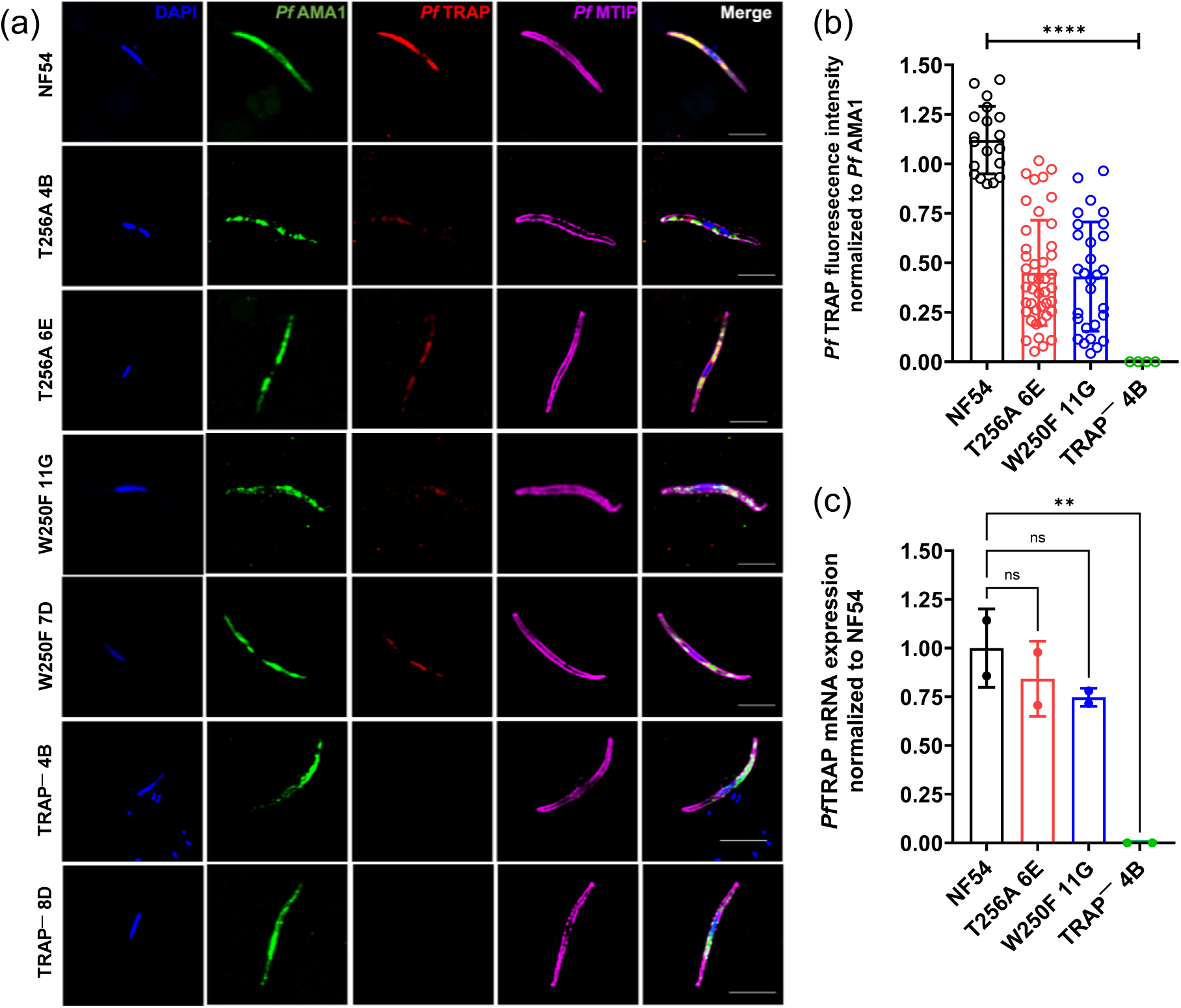
*Pf*TRAP glyco-null mutants exhibit reduced levels of *Pf*TRAP. (a) Immunofluorescence assays show localization of *Pf*TRAP (red) in hemolymph sporozoites. *Pf*AMA1 (green), a micronemal protein and *Pf*MTIP (magenta), an inner membrane complex protein, are shown to confirm the localization of *Pf*TRAP. Scale bar = 4μm. (b) Relative fluorescence intensity of *Pf*TRAP normalized against *Pf*AMA1. Each point represents an imaged sporozoite. Bars show the mean ±S.D. Both the *Pf*TRAP_T256A O-fucosylation-null mutant and the *Pf*TRAP_W250F C-mannosylation-null mutant showed a statistically significant (**** *p* < 0.0001) reduction in *Pf*TRAP compared to NF54. There was no *Pf*TRAP staining in *Pf*TRAP^—^ sporozoites. (c) Abundance of *Pf*TRAP mRNA transcripts in hemolymph sporozoites measured by qRT-PCR. Abundance is shown relative to the mean abundance in NF54. Bars show the mean ±S.D. of two technical replicates. The transcript levels in the glyco-null mutants did not differ significantly from wild type, and no transcript was detected in the knockout parasite (** *p* <0.01).

### *Pf*TRAP glyco-null mutants have reduced motility and are deficient in hepatocyte invasion

To further probe the functional relevance of glycosylation in the function of *Pf*TRAP, we assessed sporozoite gliding by measuring trails of shed *Pf*CSP on a solid substrate [49, 50] (**Fig 5a, Table S3**). We quantified the motility phenotype by categorizing sporozoite movement into four categories: sporozoites with no associated trails (non-motile), sporozoites with discharge of *Pf*CSP protein only and no motility, sporozoites with 1-4 concentric trails, and sporozoites with more than four associated trails (highly motile) (**Fig 5b**). Approximately 60% of wild-type sporozoites were highly motile, while *Pf*TRAP^—^ sporozoites were unable to glide at all. Fewer than 9% of *Pf*TRAP_T256A O-fucosylation-null sporozoites were highly motile, and fewer than 20% had any *Pf*CSP trails. Even more strikingly, none of the *Pf*TRAP_W250F C-mannosylation-null sporozoites were highly motile, and fewer than 1% displayed any motility (**Fig 5c**). These results demonstrate that *Pf*TRAP is essential for gliding motility and suggest that glycosylation of *Pf*TRAP’s TSR supports this function.

**Figure 5.**
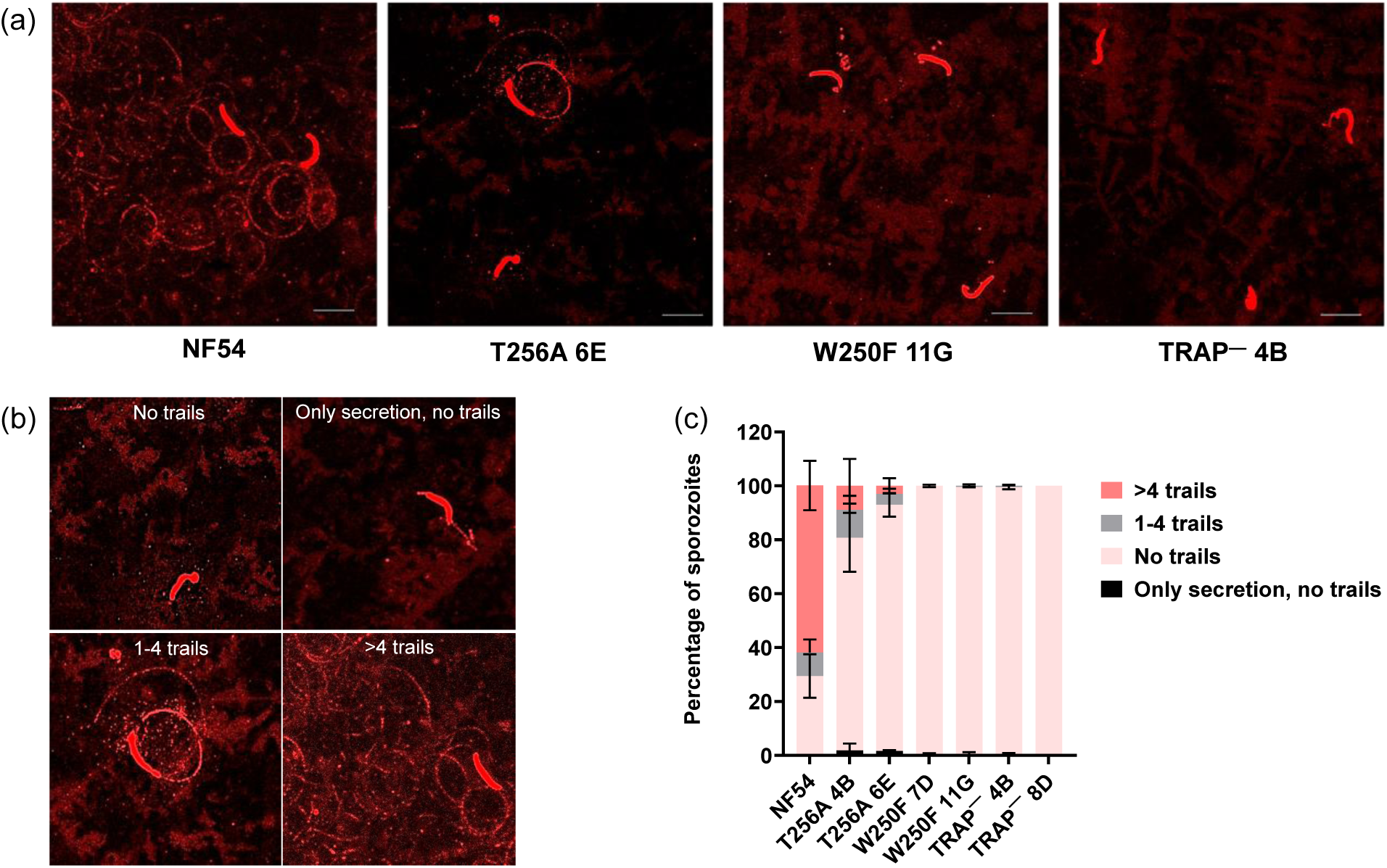
*Pf*TRAP glyco-null mutant sporozoites are unable to glide on a solid substrate. (a) Immunofluorescence assay showing the gliding trails of shed *Pf*CSP in glyco-null mutants and the *Pf*TRAP knockout parasite compared to wild-type NF54. Scale bar = 10 μm. (b) Categorization of the motility phenotype into 4 categories: sporozoites with no associated trails, sporozoites with a discharge of CSP protein and no trails, sporozoites with 1-4 concentric trails, and sporozoites with more than 4 associated trails. (c) Quantitative assessment of the ability of glyco-null mutant sporozoites to glide. Data is the mean ±S.D. of two independent experiments (**Table S3**).

To further assess sporozoite fitness, we performed an *in vitro* assay that employs flow cytometry to quantify the ability of sporozoites to invade cultured HC-04 hepatocytes [51]. We observed that the number of intracellular sporozoites was significantly reduced for the glyco-null and the knockout mutants compared to wild type, though invasion was not entirely abrogated (**Fig 6, Table S4, Fig S7**). Notably, the invasion defect was less severe for *Pf*TRAP_T256A sporozoites than for *Pf*TRAP*_*W250F sporozoites, which were comparable to *Pf*TRAP^—^. These results show that *Pf*TRAP is critical (but not essential) for invasion, and that this function is supported by glycosylation of *Pf*TRAP’s TSR.

**Figure 6.**
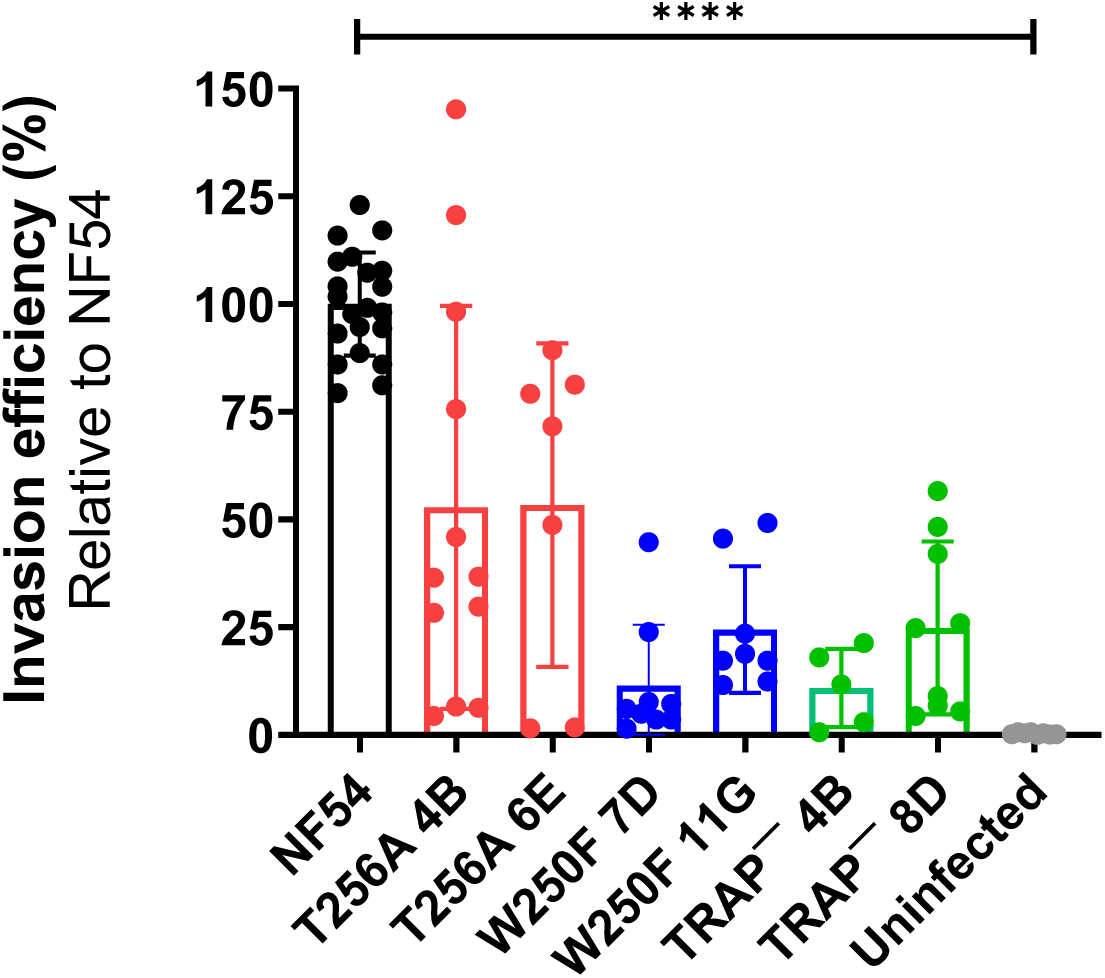
*Pf*TRAP glyco-null mutants have reduced hepatocyte invasion efficiency. Each point represents a well of HC-04 cultured hepatocytes infected with salivary gland sporozoites. Sporozoite internalization was quantified as the percent of cells positive for *Pf*CSP. Invasion efficiency was normalized to the mean internalization for NF54 in the same plate. Bars are the mean ±S.D. of replicate wells and replicate infections. The glyco-null and knockout lines exhibited significantly lower internalization than NF54 (**** *p* < 0.0001). The T256A clones exhibited higher invasion efficiency than the W250F and *Pf*TRAP^—^clones. The invasion efficiency of the W250F and *Pf*TRAP^—^ clones was equally low, but invasion was not completely blocked (**Table S4**).

### Glycosylation supports the stability and secretion of recombinant *Pf*TRAP TSR

To further explore how glycosylation supports expression of *Pf*TRAP and to investigate the effect of the point mutations on domain stability, we engineered an expression system that enabled us to recombinantly express the *Pf*TRAP TSR with or without C-Man and O-Fuc. We obtained a knockout CHO line that lacks endogenous DPY19 [26] (kind gift of Dr. Hans Bakker) and is therefore incapable of mannosylating TSRs. We adapted the adherent line to suspension growth [52] and used the Daedalus lentiviral transduction system [53] to create stable lines expressing *Pf*TRAP TSR constructs [47]. To obtain mannosylated protein, we transduced the CHO line with a codon-optimized *Pf*DPY19-coding sequence [31] (kind gift of Dr. Françoise Routier). It was critical to use the parasite DPY19 in place of the endogenous DPY19 because *Pf*TRAP expressed in mammalian cells is mannosylated at both Trp^247^ and Trp^250^ [29], whereas *Pf*TRAP co-expressed with *Pf*DPY19 is only mannosylated at Trp^250^, matching the *in vivo* glycosylation state [31]. The activities of endogenous POFUT2 and B3GLCT in mammalian cell lines result in O-Fuc and O-Fuc-Glc at Thr^256^ of *Pf*TRAP, matching the parasite *in vivo* modifications [54]. To produce *Pf*TRAP TSR without O-Fuc (and, therefore, without O-Fuc-Glc), we disrupted the endogenous POFUT2 in the *Pf*DPY19-expressing line. We also used these lines to produce *Pf*TRAP TSR with the same W250F and T256A substitutions implemented in our transgenic parasite lines. Recombinant *Pf*TRAP TSR was recovered from supernatant, enriched on Ni-NTA resin via a His-tag, and further purified by size-exclusion chromatography (SEC). Recovery was quantified by the area under the curve of the SEC peak and used as a proxy for the effect of glycosylation on protein secretion (**File S2**). The purity and glycosylation stoichiometry of the recovered *Pf*TRAP TSR was determined by MS (**File S3**).

Characterization of the recombinant *Pf*TRAP TSRs is summarized in **Table 1**. The glycopeptides detected by MS analysis of recombinantly expressed wild-type *Pf*TRAP TSR exhibited chemical signatures matching those seen in the sporozoites samples and in previous studies of recombinant *Pf*TRAP TSR [29]. Trace amounts of the protein exhibited non-canonical modification with labile masses matching hexose and deoxyhexose, likely arising from off target modification by other endogenous glycosyltransferases [55, 56], but the dominant species were consistent with the expected glycosylation. Wild-type *Pf*TRAP TSR expressed in the CHO lines was constitutively fucosylated, consistent with what is seen in the parasite, though >99% of Fuc was extended to a Fuc-Glc disaccharide, in contrast to the parasite, where the majority of *Pf*TRAP bears only the monosaccharide. Similar to previous work [31, 35], only ∼20% of detected *Pf*TRAP TSR was modified with C-Man, indicating that this modification is less critical than O-Fuc for folding and secretion of this domain. All C-mannosylation associated with the TSR was localized to Trp^250^ as seen *in vivo*. The *Pf*TRAP TSR could still be expressed in the absence of DPY19 and when C-mannosylation was blocked with a W250F substitution, but recovery of these constructs was lower than for the mannosylated wild-type protein. By contrast, expressing the *Pf*TRAP TSR in the CHO line with disrupted POFUT2 led to drastically reduced yields, resulting in quantities and purity that were not sufficient for biophysical assays. MS analysis of the recovered protein confirmed that POFUT2 activity was mostly but not completely abrogated, with <2% of the protein modified with O-Fuc or O-Fuc-Glc, while C-mannosylation was comparable to protein expressed in the parental line. Protein recovery was similarly diminished for *Pf*TRAP TSR with a T256A substitution. MS confirmed that O-Fuc and O-Fuc-Glc were completely absent from the protein, confirming Thr^256^ as the site of modification. In this protein as well as in wild-type *Pf*TRAP TSR expressed in the POFUT2^—^ line, a small amount of protein could be detected with an additional C-Man at Trp^247^, similar to what we saw in the *Pf*TRAP_T256A transgenic parasite.

**Table 1.** Effect of glycosylation on yield and thermal stability of recombinantly expressed *Pf*TRAP TSR domain. The TSR domain of *Pf*TRAP was recombinantly expressed in CHO cells lacking endogenous DPY19. a) The *Pf*TRAP TSR sequence was evaluated in its wild-type (WT) sequence or with the same W/F and T/A substitutions evaluated in the transgenic parasites. b) “+” indicates the cell line also expressed *Pf*DPY19. c) The cell lines endogenously express POFUT2 and B3GLCT. “-” indicates the endogenous POFUT2 was disrupted. d) Protein yield ± S.D. from two independent production batches was estimated from the peak area during SEC purification. The two fucosylation-null constructs (bottom two rows) were highly contaminated with background proteins (**File S2**). e) The proportion of the protein modified with a single C-Man at Trp^250^, C-Man at both Trp^247^ and Trp^250^, and O-Fuc or O-Fuc-Glc (presumably at T^256^) as determined by MS. Values are given as ± S.D. of two independent samples (**File S3**). f) The melting point ± S.D. as determined by nanoDSF from duplicate measurements of two independent samples (**File S2**). “n/a” indicates protein could not be isolated in sufficient quantity or purity to perform the assay.

| *Pf*TRAP TSR Construct<sup>a</sup> | *Pf*DPY19<sup>b</sup> | POFUT2<sup>c</sup> | Yield (μg/L of culture)<sup>d</sup> | C-Man<sup>e</sup> | C-Man×2<sup>e</sup> | O-Fuc<sup>e</sup> | O-Fuc-Glc<sup>e</sup> | T<sub>melt</sub><sup>f</sup> |
| --- | --- | --- | --- | --- | --- | --- | --- | --- |
| WT | + | + | 156 ± 85 | 22 ± 2% | 0% | 0.26 ± 0.05% | 99.7 ± 0.1% | 63 ± 1° |
| WT | - | + | 46 ± 12 | 0% | 0% | 0.29 ± 0.09% | 99.7 ± 0.1% | 55 ± 1° |
| W250F | + | + | 20 ± 17 | 0% | 0% | 0.8 ± 0.2% | 98% ± 1% | 41 ± 1° |
| WT | + | - | ~5 | 25% | 0.34% | 0.02% | 1.4% | n/a |
| T256A | + | + | ~3 | 14% | 0.35% | 0% | 0% | n/a |

In order to elucidate the importance of C-mannosylation and O-fucosylation for folding and stability of the *Pf*TRAP TSR, we used nanoscale differential scanning fluorimetry (nanoDSF) to assess the thermal stability of the recombinantly produced glycoforms described above (**Table 1**, **File S2**). Even though only ∼20% of the protein co-expressed with *Pf*DPY19 was mannosylated, its melting temperature (T_melt_) was significantly higher than that of the unmannosylated form, confirming that mannosylation of Trp^250^ stabilizes the TSR. The observed T_melt_ of the W250F construct was even lower than that of the unmannosylated wild-type construct, suggesting that this substitution was deleterious to the stability of the domain. It was not possible to obtain sufficient material to perform nanoDSF for wild-type protein expressed in the absence of POFUT2 or with the T256A substitution that prevents O-fucosylation, suggesting that O-fucosylation is critical for *Pf*TRAP stability. Taken together, our results suggest that *Pf*TRAP requires O-fucosylation for correct folding and secretion from the ER, and that blocking this modification by the T256A substitution prevents correct folding, leading to protein instability. For its part, C-mannosylation stabilizes the TSR and also supports protein folding, but to a lesser extent than O-fucosylation. Substituting the mannosylated Trp with Phe further destabilizes the domain, leaving open the possibility that this substitution caused the levels of *Pf*TRAP seen in the *Pf*TRAP_W250F transgenic parasites to be lower than what would have been seen if it were possible to produce sporozoites lacking DPY19.

## DISCUSSION

Work in recent years has provided insights into the specialized structure and function of TSR domains and has shown that glycosylation of these domains with C-Man, O-Fuc, and O-Fuc-Glc supports their folding, secretion, and function. The *Plasmodium* genome encodes ten TSR-bearing proteins and the glycosyltransferases to modify them, so it can be inferred that O-fucosylation and C-mannosylation in *Plasmodium* are important for maintaining the correctly folded and functional state of these proteins. However, evidence to date shows that the role and essentiality of TSR glycosylation varies from protein-to-protein, so the importance of each TSR-bearing protein and glycosite must be considered on an individual basis. We have done that here by introducing conservative, single-residue substitutions into the TSR of an essential *P. falciparum* invasin, *Pf*TRAP, to prevent modification of the two known glycosites on the protein, and have demonstrated that these glycosylations are critical for protein expression and function.

In order to have a baseline for comparison of any phenotypes produced by our glyco-null mutants, we produced a *Pf*TRAP^—^ mutant. A rodent malaria *Pb*TRAP^—^ mutant has been characterized [7], but ours is the first study to comprehensively explore the essentiality of *Pf*TRAP. Similar to the *Pb* studies, *Pf*TRAP was dispensable for blood stage development and transmission to mosquitoes, consistent with the lack of *Pf*TRAP expression in these stages. *Pf*TRAP is highly expressed in oocyst sporozoites that develop in the mosquito midgut [57], but was dispensable for sporozoite egress from oocysts and entry into the mosquito hemolymph. Our experiments demonstrate that the first bottleneck for *Pf*TRAP^—^ parasites is colonization of the mosquito salivary glands. The role of *Pf*TRAP in salivary gland colonization is poorly understood. It has been reported that *Pf*TRAP interacts via its vWA domain with proteins found on the surface of mosquito salivary glands [58], but to date no mosquito ligands for the TRAP TSR have been identified.

From the limited number of *Pf*TRAP^—^ sporozoites that could be recovered from dissected salivary glands, we performed *in vitro* assays to assess parasite fitness relating to transmission to the human host. The gliding motility assays showed that, like *Pb*TRAP [7], *Pf*TRAP is essential for this process; *Pf*TRAP^—^ sporozoites were completely incapable of producing the trails of shed *Pf*CSP characteristic of the highly motile wild-type sporozoites. In the *in vitro* assay with cultured HC-04 hepatocytes, the number of hepatocytes with internalized *Pf*TRAP^—^ sporozoites was significantly reduced compared to wild-type parasites, though the signal from the assay was still above the baseline recorded for uninfected cells. These results are consistent with the previous observation that *Pb*TRAP^—^ parasites were unable to infect rats by mosquito bite, but intravenous injection of large numbers of sporozoites could lead to a blood stage infection, albeit with a delay-to-patency corresponding to a 10,000-fold reduction in infectivity [7]. The mechanism by which *Pf*TRAP facilitates liver invasion is the subject of active research. In addition to its role in gliding motility, *Pf*TRAP interacts with hepatocyte proteins [59–61] and recognizes sulfated proteoglycans on the hepatocyte surface [62], mediated at least in part by its TSR [63].

In order to elucidate how TSR glycosylation supports the essential functions of *Pf*TRAP, we generated transgenic *Pf* lines with conservative substitutions at Thr^256^ or Trp^250^, the sites of O-fucosylation and C-mannosylation, respectively. MS analysis of these parasites provided direct evidence that the single-residue substitutions at the predicted glycosites had the desired effect, *i.e.*, replacing Thr^256^ with Ala blocked O-fucosylation and replacing Trp^250^ with Phe blocked C-mannosylation. The observation that a portion of *Pf*TRAP was C-mannosylated at both TSR domain Trp residues in the *Pf*TRAP_T256A mutant was unexpected, as the original report of *Pf*TRAP glycosylation in salivary gland sporozoites [27] and the analysis of oocyst sporozoites presented here found only Trp^250^ to be C-mannosylated. We hypothesize that the primary structure or folding kinetics of the domain renders Trp^247^ of *Pf*TRAP TSR mostly refractory to C-mannosylation by *Pf*DPY19, possibly similar to how *Pv*TRAP and *Pf*CSP are entirely refractory to C-mannosylation. We further hypothesize that destabilization of the TSR domain by the lack of O-Fuc shifted the equilibrium of folding to favor additional stabilization by the second C-Man at Trp^247^, producing a small but detectable proportion of this glycoform in the O-fucosylation-null mutant as well as in the recombinant *Pf*TRAP TSR. Future work will investigate this phenomenon in detail.

Our results show that single-residue substitution of the *Pf*TRAP glycosites results in a significant reduction of protein abundance despite normal levels of transcript, evidence that O-fucosylation and C-mannosylation are required for the correct folding of *Pf*TRAP. However, immunofluorescence and MS confirmed that *Pf*TRAP was still present in the parasites. Furthermore, IFA showed glyco-null *Pf*TRAP co-localizing with the membrane antigen *Pf*AMA1, which, like *Pf*TRAP, accumulates in micronemes until secreted to the parasite surface [48], suggesting that some amount of glyco-null *Pf*TRAP was secreted from the ER and correctly localized to the secretory vesicles. In many cases, when levels of an essential protein are reduced but not completely eliminated (e.g., as can happen in conditional knock-down approaches), the associated fitness defect is reduced or absent compared to a complete gene knockout [64]. In the experiments we describe here, however, glyco-null mutants showed defects as severe as the knockout, suggesting that correctly glycosylated *Pf*TRAP is essential for these processes. It has been shown in other systems that glycosylation of TSRs can mediate their function by stabilizing binding domains and even by directly interacting with binding partners [65]. Interestingly, the O-fucosylation-null mutant *Pf*TRAP_T256A retained some gliding motility in the *in vitro* assay, albeit at highly reduced levels, while the C-mannosylation-null *Pf*TRAP_W250F mutant mirrored the *Pf*TRAP^—^ parasite in being almost completely incapable of gliding on a solid substrate. Similarly, the invasion defect for *Pf*TRAP_T256A sporozoites was less severe than for *Pf*TRAP_W250F parasites, whose deficiency was comparable to the knockout. These results suggest that Trp^250^ and its C-mannosylation are essential for the role of *Pf*TRAP in gliding motility and invasion, while Thr^256^ and its O-fucosylation are less critical (though still important), observations which may lend insight into how *Pf*TRAP’s TSR supports these aspects of parasite fitness.

Since the intention of the site-directed mutagenesis experiments described here was to determine the importance of each glycan for protein structure and function, we replaced these glycosites with amino acids that would prevent glycosylation while minimizing disruption to the protein structure caused by the substitution itself. Ala is the most common amino acid used in site-directed mutagenesis studies (*e.g.*, alanine scanning [66, 67]) because it can usually replace most residues without affecting the secondary structure of the protein. Substitution of Ser and Thr with Ala is common for preventing phosphorylation [68], and thus was the logical choice for preventing O-fucosylation of *Pf*TRAP Thr^256^. However, Ala substitution is more problematic when the sidechain is integral to protein structure, as is often the case with Trp, especially in TSRs [24]. One of the characteristic features of TSRs is a fold comprising three anti-parallel strands, held together by disulfide bonds, wherein Trp residues on one strand intercalate between Arg residues on the opposite strand [69]. It is the Trp residues of this so-called Trp-Arg ladder that are modified with C-Man. As has been done by others who study TSR glycosylation [24], we selected Phe as the best substitute for Trp. Both residues have planar, non-polar, aromatic sidechains, and as such Phe is capable of interacting with Arg in the same type of non-covalent stacking that produces the Trp-Arg ladder critical to TSR structure [70]. Furthermore, studies with recombinant TSRs have shown that Trp/Phe substitution in the DPY19 recognition motif (*e.g.*, WXXW→WXXF as we did here) can still allow C-mannosylation of the first Trp while Phe itself cannot be C-mannosylated [24, 71]. In order to better parse the effect of these substitutions and modifications on protein stability, we recombinantly expressed the *Pf*TRAP TSR domain with the same modifications introduced in our transgenic parasite lines. We confirmed that O-fucosylation is critical for folding and secretion of the *Pf*TRAP TSR and that this effect was recapitulated by blocking the modification via substitution with Ala at Thr^256^, providing evidence that fitness defects in our transgenic *Pf*TRAP_T256A parasite are a direct result of the importance of O-fucosylation for *Pf*TRAP expression and not an artifact of the substitution. C-mannosylation of recombinant *Pf*TRAP TSR was less critical for protein folding and secretion, but preventing the modification correlated with lower protein recovery, and our melting point assay confirmed that the thermostability of the domain was significantly lower in the absence of mannosylation. These results confirm that mannosylation of Trp^250^ stabilizes the *Pf*TRAP TSR and aids in folding of the domain, which is likely why the site is constitutively modified in the parasite *in vivo*. Substituting the C-mannosylation site at Trp^250^ with Phe further destabilized the protein as evidenced by a significant decrease in T_melt_ compared to the unmannosylated protein with wild-type sequence. Phe was therefore a less-than-ideal substitute for Trp at this site, and it cannot be stated with certainty how much of the observed protein abundance defect observed in the *Pf*TRAP_W250F transgenic parasites was the result of this substitution as opposed to the lack of mannosylation. This fact only serves to highlight how critical conserved Trp residues are to the structure and function of TSRs. It is remarkable that the sequence motifs required for TSR glycosylation are so highly conserved across phyla, and that a single-residue substation in a protein domain could render a protein functionally dead.

Previous studies that deleted POFUT2 and DPY19 in *Plasmodium* demonstrated that TSR glycosylation is essential for parasite virulence. However, since multiple TSR-bearing proteins are expressed at any given stage, and the role of TSR glycosylation varies with each protein, that work did not answer the question of which proteins were affected or why glycosylation was required for protein function. The work we have described here, in which we prevented single modifications on a single protein, demonstrates how both O-fucosylation and C-mannosylation of a single essential invasin, TRAP, can support parasite fitness. More broadly, because TSRs and the enzymes that glycosylate them are highly conserved across diverse phyla, the results of this work will impact the field of glycobiology by providing new insights into the function of these rare but critical protein modifications.

## MATERIALS AND METHODS

### Generation of *Plasmodium falciparum* transgenic glyco-null mutants

The glyco-null and knockout mutant parasite lines were generated using CRISPR-Cas9-mediated gene editing, utilizing the pFC-L3 cloning vector [72]. Three guide sequences were individually cloned into the guide cloning site within the BsmBI restriction enzyme site. Recombination arms containing the specific mutations were synthesized by GenScript and inserted into the vector containing the guide sequences via NotI and SalI cloning sites. The sequences of the guide RNA and primers used in the process are provided in **Table S1**. To introduce the mutations, 5% sorbitol-synchronized *Pf* NF54 ring-stage parasites were co-transfected with 100 µg of plasmid DNA using electroporation at 0.31 kV and 950 µF with a Bio-Rad Gene Pulser, as previously described [73]. Following transfection, positive selection was applied by treating the parasites with 8 nM WR99210 (WR; Jacobus Pharmaceuticals) 24 hours post-transfection for a period of 5 days. After this selection period, media was changed daily to remove the drug pressure, and parasites were allowed to grow until they were detectable on thick smears, typically 17-21 days post-transfection. Recombinant parasites were first screened by PCR genotyping and subsequently cloned by limiting dilution. To confirm the presence of the desired mutations, restriction fragment length polymorphism analysis was performed by amplifying the TSR domain region with primers listed in **Table S1**. The PCR products were then digested with AluI and EcoRI restriction enzymes to detect the *Pf*TRAP_T256A and *Pf*TRAP_W250F mutations, respectively. Positive clones were further confirmed by sequencing to verify the correct genomic sequence, ensuring the accuracy of the induced mutations.

### *Plasmodium falciparum* culturing and transmission to mosquitoes

Asexual cultures of *Pf* NF54 strain were maintained using standard laboratory procedures [74]. Briefly, the parasites were cultured *in vitro* in type O+ erythrocytes and RPMI 1640 medium supplemented with 50 µM hypoxanthine. The cultures were aerated with a gas mixture of 5% CO₂, 5% O₂, and 90% N₂. Parasitemia was monitored daily to assess parasite growth.

To initiate gametocyte development, healthy asexual cultures containing predominantly rings were set up at 1% parasitemia and 5% hematocrit in a 6-well plate. The gametocyte cultures were maintained for 14 days. Media changes were done daily to provide fresh nutrients and maintain optimal conditions for gametocyte maturation. On day 15, stage V gametocytemia was assessed, and female *Anopheles stephensi* mosquitoes (4–7 days old) were fed with the gametocyte cultures at 0.3% gametocytemia and 50% hematocrit using a membrane feeding apparatus [46].

The fed mosquitoes were maintained at 27°C and 75% humidity for 17 days, with access to a 10% w/v dextrose solution and 0.05% w/v p-aminobenzoic acid in water. Oocyst development in the mosquito midguts was monitored on day 7 after feeding. On day 7, a supplemental feed of a 50:50 mixture of blood and serum was provided to the mosquitoes. Mosquitoes were dissected on day 10 or 11 to collect midguts and on day 16 or 17 to collect salivary glands for sporozoite purification.

### Isolation of hemolymph sporozoites

To isolate hemolymph sporozoites, mosquitoes were dissected on day 15 or 16 after feeding [75]. The last segment of the abdomen was excised using a 1 mL syringe. Dissected mosquitoes were then flushed by inserting a 1 mL insulin syringe into the lateral side of the thorax and injecting Schneider’s media. This procedure allowed for the drainage of hemolymph from the abdomen. The collected hemolymph was carefully harvested on the dissection stage and transferred to a microcentrifuge tube.

### Trypsinization of the salivary glands of the mosquitoes

To determine the proportion of internalized sporozoites among salivary gland–associated sporozoites, salivary glands were dissected and incubated with trypsin (50 μg/mL) for 15 minutes at 37°C [7]. Trypsin is a protease enzyme that detaches sporozoites that are only loosely attached to the surface of the glands. After incubation, the samples were centrifuged for 5 minutes to pellet the glands and the supernatant, which contained the detached sporozoites, was removed. The remaining salivary gland tissue was then ground to release the internalized sporozoites into solution.

### Proteomic analysis of sporozoites and recombinant proteins

Extended sample preparation and mass spectrometry (MS) methods are available along with the raw data in ProteomeXchange. Sporozoites analyzed by MS were first purified by Accudenz discontinuous gradient [76], then digested with trypsin using an SP3 method [77]. Recombinant proteins were digested with trypsin in-solution. Liquid chromatography (LC) was performed with an EASY-nLC 1000 (Thermo Fisher Scientific) or a Vanquish Neo (Thermo Fisher Scientific) with a trap-elute setup. The trap column was a PepMap 100 C18 (Thermo Fisher Scientific #164946) with 75 µm i.d. and a 2 cm bed of 3µm 100 Å C18 or a PepMap Neo (Thermo Fisher Scientific; Cat# 174500) with 300 µm i.d. and a 5 mm bed of 5µm 100 Å C18. The analytical column was an EASY-Spray column (Thermo Fisher Scientific) with 75 µm i.d. and a 50 cm bed (Cat# ES903) or 15 cm bed (Cat# ES904) of 2µm 100 Å C18 operated at 55°C or 45°C, respectively. The LC mobile phases consisted of buffer A (0.1 % v/v formic acid (FA) in water) and buffer B (0.1 % v/v FA in ACN). Sporozoite digests were separated via a gradient of 4% B to 28% B over 60 min and recombinant *Pf*TRAP TSR peptides were separated over a 20 or 30 min gradient. The MS method employed high-mass-accuracy Orbitrap scans. The method for sporozoite samples comprised a precursor scan from 400-1600 *m/z*, seven precursor selected ion monitoring (SIM) MS1 scans, and seven MS2 SIM scans employing HCD fragmentation. The entire duty cycle of the MS was devoted to obtaining high-sensitivity precursor and fragment scans of the potential glycoforms of the *Pf*TRAP peptide containing Thr^256^ and Trp^250^. The seven targeted *m/z* values correspond to all possible doubly charged permutations of the TSR glycopeptide, i.e., potential addition of C-Man to Trp^247^ and Trp^250^ and O-Fuc or O-Fuc-Glc at Thr^256^. A volume of tryptic digest corresponding to 2.5×10^5^ to 6.0×10^5^ sporozoites was analyzed for each SIM experiment. The method for recombinant protein comprised a precursor MS1 scan followed by seven data-dependent HCD scans. Dynamic exclusion was not enabled. An inclusion list of *m/z* matching the seven potential glycoform masses directed the mass analyzer to fragment these ions whenever detected, and otherwise to select the most abundant precursor ions detected in the MS1 scan. A volume of digest corresponding to nominally 3 to 5 pmol of protein was analyzed for each experiment.

### Proteomics data analysis

All mass spectrometry data, including raw data, search parameters, databases, and results have been deposited to the ProteomeXchange Consortium [78] via the MassIVE partner repository with the dataset identifier PXD064887. Peptide spectrum matches (PSM) have been assigned Universal Spectrum Identifiers (USI) [79]. Raw mass spectrometry data were converted to mzML with centroided peaks using MSConvert version 3.0.19106 (Proteowizard [80]), then searched and analyzed with the Trans-Proteomic Pipeline (TPP) v 7.3.0 [81] running on a Slurm 23.11.4 cluster running under Ubuntu 20.04. Sporozoite spectra were searched against a combined FASTA database assembled from the reference *P. falciparum* 3D7 [82, 83] and *Anopheles stephensi* [84] databases (PlasmoDB [85] and VectorBase [86], VEuPathDB [87] version 68) and the common Repository of Adventitious Proteins (www.thegpm.org/cRAP). Recombinant protein spectra were searched against a combined FASTA database assembled from the *Cricetulus griseus* (Chinese hamster) reference proteome from UniProt [88], cRAP, and the recombinant *Pf*TRAP TSR construct sequences. De Bruijn decoy protein entries were generated using a tool in the TPP. Two decoys were created for each real entry, denoted DECOY0 and DECOY1. Spectra were searched using Comet version 2025.01 rev. 1 [89]. Fully tryptic peptides with no missed cleavages were allowed, with variable modification of 15.9949 Da (oxidation) at Met and Trp and 162.0528 (Man) at Trp with potential neutral loss of 120.0423 Da for C-mannosylation. Variable modification of 162.0528 at Phe was also allowed when searching the data from the W250F transgenic parasite. Potential neutral losses of 146.0579 (Fuc) and 308.1107 (Fuc-Glc) were accounted for with the mass offset parameter [29]. For the recombinant proteins, neutral loss of 162.0528 and 470.1635 (Fuc + Fuc-Glc) were also allowed to account for trace occurrence of isobaric species with labile hexose and deoxyhexose. The precursor mass tolerance was set to +/−10 ppm. Isotope error was enabled (i.e., allowing that the identified precursor mass may be at the second isotope peak rather than the monoisotopic peak), which was critical for the SIM data since the MS2 isolation windows were offset by 1.0 Da, i.e., centered on the second isotope peak of the doubly charged peptide ions. Only high-quality PSM (Comet Expect scores <1×10^−5^ corresponding to a decoy-estimated FDR of 0%) were taken for further analysis. The identities of *Pf*TRAP TSR peptides were verified manually by inspecting annotated PSMs in the Lorikeet (https://github.com/UWPR/Lorikeet/) viewer in the TPP and in Quetzal [90]. Relative abundances of peptide glycoforms were calculated from the peak heights of extracted ion chromatograms produced using Xcalibur Qual Browser v 4.2.28.14. (**File S1 & S3**).

### qPCR analysis of the sporozoites

Hemolymph sporozoites were isolated at day 15 or 16 after infectious blood meal. RNA was isolated from 1×10^6^ sporozoites using an miRNeasy Mini RNA extraction kit (Qiagen; Cat# 217004). RNA was reverse transcribed into cDNA using the oligo dT-primer and Quantitect reverse transcription kit (Qiagen; Cat# 205313). Synthesized cDNAs were used in qPCR analysis to quantify the expression of TRAP. 18S rRNA housekeeping gene was used as a reference. Reactions were performed in QuantStudio 3 real-time PCR system (Applied Biosystems by Thermo Fisher Scientific) using 2× SYBR Green qPCR Master Mix (Selleckchem; Cat# 21203) according to the manufacturer’s instructions. PCR cycling conditions were 95^°^C (10 min), 35 PCR cycles of 95^°^C (15 Sec), 60^°^C (1 min), and final hold at 10^°^C (10 Sec). Primer sequences are listed in **Table S1**.

### Recombinant production of *Pf*TRAP and mouse immunizations for polyclonal serum production

Production and purification of recombinant *Pf*TRAP was performed similarly to the previously described protocol [47]. Briefly, an amino acid sequence corresponding to the *Pf*TRAP ectodomain (codons 24–511 from PF3D7_1335900) was used to generate a version codon-optimized for production in the human (HEK293) expression system, with an upstream tPA leader and a C-terminal His_8_ followed by an AviTag. The construct sequence was cloned into pcDNA3.4 plasmid vector (Thermo Fisher Scientific) and used for transfection of suspension FreeStyle 293F cells (Thermo Fisher Scientific) grown in the FreeStyle 293 Expression Medium (Thermo Fisher Scientific). At 5 days following transfection, the cells were removed by centrifugation to harvest the culture supernatant. Following addition of NaCl (350 mM final concentration) and NaN_3_ (0.02% final concentration), the supernatant was passed through HisPur Ni-NTA resin (Thermo Fisher), and the resulting purified protein was further polished by size exclusion chromatography (SEC) over a standardized column containing Superdex 200 resin (Cytiva), producing the final stock of purified *Pf*TRAP. Intramuscular immunizations were carried out using 8–12-week-old BALBc/J mice (Jackson Labs) at 3 time points on weeks 0/2/6 using 20 µg protein in 50% AddaS03 adjuvant (InvivoGen) per dose, formulated according to manufacturer’s instructions. Seven days following the final immunization, the animals were sacrificed to harvest blood, which was centrifuged to remove red blood cells to prepare polyclonal sera for use in in vitro assays.

### Immunofluorescence analysis of sporozoites

The sporozoites were isolated and centrifuged at 6,000 RPM for 6 minutes, then resuspended in 80 µL of 4% paraformaldehyde (PFA) for fixation at room temperature for 20 minutes. After fixation, the sporozoites were spun again at 6,000 RPM for 6 minutes and washed once with phosphate-buffered saline (PBS). Approximately 2.0×10^4^ sporozoites were loaded per well on a microscope slide and allowed to air dry overnight in a fume hood. The sporozoites were then permeabilized and blocked by incubating the slide at room temperature for 1 hour in blocking solution, which consisted of PBS containing 2% BSA and 0.2% Triton X-100. Following blocking, the sporozoites were incubated for 2 hours at room temperature with primary antibodies (anti-*Pf*TRAP, anti-*Pf*AMA1, anti-*Pf*MTIP) in blocking solution. After primary antibody incubation, the slides were washed three times with PBS. The sporozoites were then incubated for 1 hour at room temperature with secondary antibodies (anti-mouse 594, anti-rat 488, anti-rabbit 647) in blocking solution. The slides were washed five times with PBS and mounted with Prolong Antifade reagent to preserve fluorescence.

### Sporozoite motility assay

Gliding assays were conducted as previously described [50]. Chamber slides with eight wells (Thermo Fisher Scientific; Cat# 154534) were coated with anti-*Pf*CSP 2A10 antibody (10 µg/mL dilution in PBS) overnight at room temperature. Each well was seeded with 2.0×10^4^ salivary gland sporozoites and incubated for 3 h at 37°C in 5% CO₂, using complete DMEM containing 10% FBS. After incubation, the samples were fixed with 4% PFA at 37°C for 20 minutes. To visualize the sporozoites, the wells were incubated with primary anti-*Pf*CSP 2A10 (1:500 dilution) antibody for 2 h at room temperature, followed by secondary goat anti-mouse Alexa 488 or 594 (1:500 dilution in 3% BSA). The gliding sporozoites and their associated trails were observed using a BZ-X 810 Keyence microscope. For each condition, multiple fields (≥100 sporozoites) were counted at 20× magnification across two independent experiments.

### Cell invasion assay

The ability of sporozoites to invade hepatocytes was assessed by a fluorescence-activated cell sorting (FACS)-based invasion assay as previously described [51]. Briefly, HC-04 hepatoma cells were cultured in Dulbecco’s Modified Eagle Medium supplemented with 10% heat-inactivated fetal bovine serum (FBS) at 37°C in 5% CO₂ atmosphere. Cells were split every 2–3 days once they reached approximately 90% confluency. HC-04 cells were seeded at 7.5×10^4^ per well in a 96-well plate that was pre-coated with rat tail collagen. After 24 hours of incubation, 3.5×10^4^ sporozoites (MOI=0.47) were added to each well and incubated for 3 hours. The cells were then trypsinized and further processed to obtain a single-cell suspension. The cells were fixed and permeabilized using BD Cytofix/Cytoperm solution (BD Biosciences; Cat# 51-2090KZ) for 15 min on ice and subjected to intracellular staining with anti-*Pf*CSP monoclonal antibody 2A10 conjugated with Alexa Fluor 647 (1:50 dilution) at room temperature for 1 h on a shaker. The cells were then washed with FACS buffer (1% FBS in PBS) and resuspended into FACS buffer containing 2% PFA. The samples were sorted on a BD LSR II and the data was analyzed using FlowJo v10.10.0. Two or three replicate wells from two or three independent experiments were quantified. Invasion was quantified as the percent of cells positive for CSP. Relative invasion efficiency was quantified by subtracting the mean invasion of uninfected HC-04 and normalizing to the mean invasion of NF54-infected HC-04 (**Table S4**)

### Generation of cell lines for recombinant protein production

A CHO-derived cell line lacking DPY19, gift of Dr. Hans Bakker [26], was adapted to suspension growth as described previously [52]. The lentivirus-based Daedalus transduction system [53] was used to produce cell lines stably expressing the *Pf*TRAP TSR and *Pf*DPY19. *Pf*TRAP TSR constructs [47] (**File S2**) were cloned into the Daedalus vector and lentiviral vector stocks were produced by co-transfection with packaging and pseudotyping constructs (psPAX2 and pMD2.G, respectively), gifts of Didier Trono (Addgene plasmids #12260 and #12259) [53]. To enable identification of cells co-transduced with both *Pf*DPY19 and *Pf*TRAP constructs, we generated an mCherry-expressing version of the Daedalus vector by replacing EGFP-coding sequence between NcoI and XbaI restriction sites, producing Daedalus-mCherry. DNA sequence encoding *Pf*DPY19 (UniprotKB:C0H4S1_PLAF7) and codon-optimized for expression in mammalian cells, gift of Dr. Françoise Routier [31], was cloned into the Daedalus-mCherry plasmid. Knockouts of the *Cricetulus griseus pofut2* gene (NCBI gene ID: 100765129) in the CHO + *Pf*DPY19 line were generated by introducing genomic lesions using CRISPR/Cas9. Guide sequences (**File S2**) were cloned into the lentiCRISPR v2 vector, gift of Feng Zhang (Addgene plasmid # 52961; http://n2t.net/addgene:52961; RRID:Addgene_52961), incorporated into lentiviral vectors and individually used to transduce the CHO line. Following expansion, cells were plated at clonal density (3 cells per well) and cultured subcloned by limiting dilution. Subclones were expanded and their genomic DNA was extracted using QuickExtract DNA Extraction Solution (Lucigen) from cell pellets for subsequent screening by PCR. In order to generate diagnostic PCR products covering multiple CRISPR/Cas9 lesion sites, 2 PCR amplifications using reactions containing primers Cg-pofut2-oligo1+Cg-pofut2-oligo2 and Cg-pofut2-oligo3+Cg-pofut2-oligo4 (**File S2**), containing a complimentary overlap region, were purified, combined and used for further amplification with the Cg-pofut2-oligo1+Cg-pofut2-oligo4 terminal primers to generate the final diagnostic PCR product. The diagnostic PCR product was used for Sanger sequencing (Azenta), and the chromatogram data obtained from subclone amplicon was compared to that from the parental lines using decodr.org service [91] to identify lesions with the highest likelihood for gene disruption (e.g., indels that disrupt the reading frame or mutations predicted to affect intron splicing) for further work.

### Recombinant protein purification

Cells transduced with the target TSR constructs were expanded in CD OptiCHO (Thermo Fisher Scientific) serum-free medium supplemented 8 mM GlutaMAX (Thermo Fisher Scientific) and grown for 5–7 days until viability fell below 80%. Culture supernatants were harvested by centrifugation and supplemented with NaN_3_ (to 0.2% final concentration) and NaCl (to 350 mM final concentration). His-tagged protein was purified from treated culture supernatants by immobilized-metal affinity chromatography (IMAC) using HisPur Ni-NTA resin (Thermo Fisher Scientific) in buffer EQ (25 mM Tris, pH 8, 300 mM NaCl and 0.2% NaN3), supplemented with 40 mM imidazole at the wash step and with 200 mM imidazole at the elution step. Following concentration of the protein-containing fractions, the samples were further purified by SEC on a calibrated Superdex 200 Increase 10/300 GL column (Cytiva) in HBS-E(10 mM HEPES, pH 7, 150 mM NaCl, 2 mM EDTA). The area under the curve of the SEC peaks was used to quantify protein yield from each production run (**File S2).**

### Measurement of protein thermal stability

Nanoscale differential scanning fluorimetry (nanoDSF) was performed using UNCLE (Unchained Labs) with protein samples equilibrated in the HBS-E buffer. Samples were ramped at 25–95° C at 1° C/min and intrinsic tryptophan fluorescence and static light scattering (SLS) were continuously monitored following excitation at 266 nm. Thermal melt profiles were derived from 350/330 nm fluorescence intensity ratios, with T_melt_ estimates generated using the first derivative calculated with the gratia R package (**File S2**).

### Statistics

Unless stated otherwise, statistical significance was assessed in GraphPad Prism by one-way ANOVA with *p*-values adjusted by Dunnett’s multiple comparisons test.

## RESOURCE AVAILABILITY

### Materials availability

Parasite lines generated in this study are available upon request.

### Data availability

All data supporting the findings of this study are available within the article and its Supplementary Information files, or are available from the corresponding authors upon request. Proteomics data were deposited to the ProteomeXchange Consortium (http://www.proteomexchange.org) with the identifier PXD064887 via MassIVE (https://massive.ucsd.edu/).

## ACKNOWLEDGMENTS

We thank the insectary staff at Seattle Children’s Research Institute for their support in rearing mosquitoes for parasite transmission studies. We thank Dr. Roland Strong at the Fred Hutchison Cancer Research Center for access to nanoDSF instrumentation. We thank Dr. Hans Bakker and Dr. Françoise Routier of Hannover Medical School for the kind gift of the CHO DPY19^—^ line and the *Pf*DPY19 plasmid and for fruitful discussion.

## AUTHOR CONTRIBUTIONS

### CRediT Designations

Conceptualization: PG, AMV, KES

Data curation: PG, VV, KES

Formal analysis: PG, VV, KES

Funding acquisition: RLM, DNS, AMV, KES

Investigation: PG, VV, NR, LP, HP, GZ, MK, AW, NC, EK, KES

Methodology: PG, VV, AMV, KES

Project administration: DNS, AMV, KES

Resources: DNS, AMV, KES

Supervision: RLM, SHIK, DNS, AMV, KES

Visualization: PG, VV, KES

Writing – original draft: PG, VV, KES

Writing – review & editing: PG, VV, NR, LP, HP, GZ, MK, DNS AMV, KES

## SUPPLEMENTARY FIGURES

**Figure S1. Schematic of the construct used to generate glyco-null mutants.**

**Figure S2. Schematic of the construct used to generate the *Pf*TRAP^—^ parasite.**

**Figure S3. *Pf* TRAP^—^ and glyco-null mutants exhibit normal sexual stage development.**

**Figure S4. Distribution of sporozoites in the oocyst, hemolymph, and salivary glands of the mosquitoes 15 or 16 days after infectious blood meal.**

**Figure S5. Trypsinization of salivary glands to remove uninvaded sporozoites.**

**Figure S6. *Pf*TRAP glyco-null mutants exhibit reduced levels of *Pf*TRAP in salivary gland sporozoites.**

**Figure S7. Representative flow cytometry data showing defects in invasion of HC-04 cells by *Pf*TRAP glyco-null mutants.**

## SUPPLEMENTARY FILES

**File S1.** Extracted ion chromatograms and mass spectra of the *Pf*TRAP glycopeptide detected in sporozoites

**File S2.** Recombinant *Pf*TRAP TSR data

**File S3.** Extracted ion chromatograms and mass spectra of the *Pf*TRAP glycopeptide detected in recombinant *Pf*TRAP TSR

## SUPPLEMENTARY TABLES

**Table S1**. List of guides and primers used in the study

**Table S2.** Parasite line phenotype data shown in Fig 2, Fig S3, Fig S4, and Fig S5

**Table S3.** Motility phenotype data shown in Fig 5

**Table S4.** Sporozoite internalization data shown in Fig 6

