## Supplementary Figures for "Glycosylation of *Plasmodium falciparum* TRAP supports sporozoite motility and invasion"

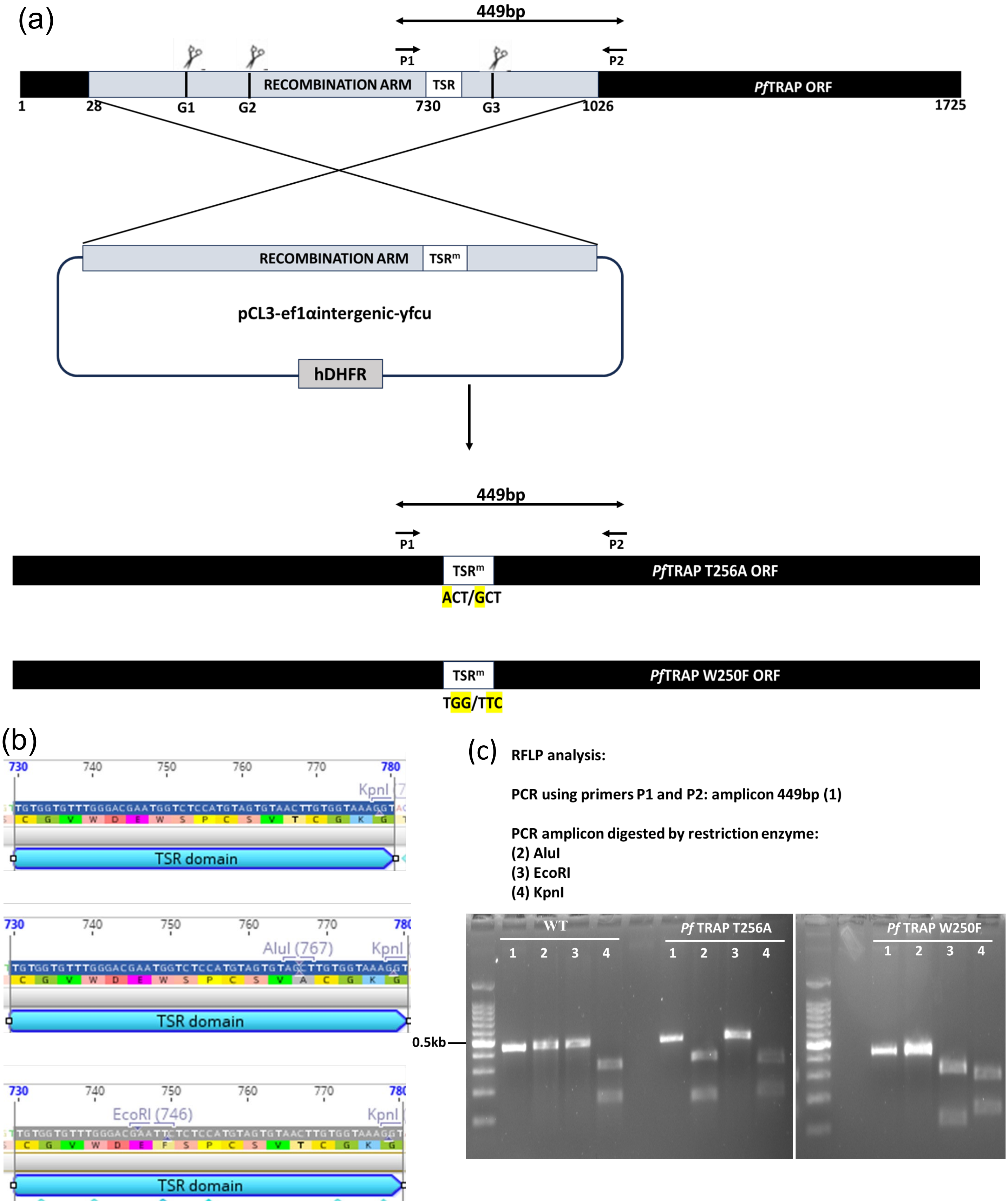


**Figure S1. Schematic of the construct used to generate glyco-null mutants.**
(a) Strategy used to generate *Pf*TRAP glyco-null mutants using the pcl3-pfc-EF1α-yFCU cloning vector.
(b) Restriction sites generated upon introduction of mutation in the TSR domain. (c) RFLP analysis to validate generation of glyco-null mutant parasites. The sequences of the guides and primers used for this cloning are listed in Table S1.

(a)

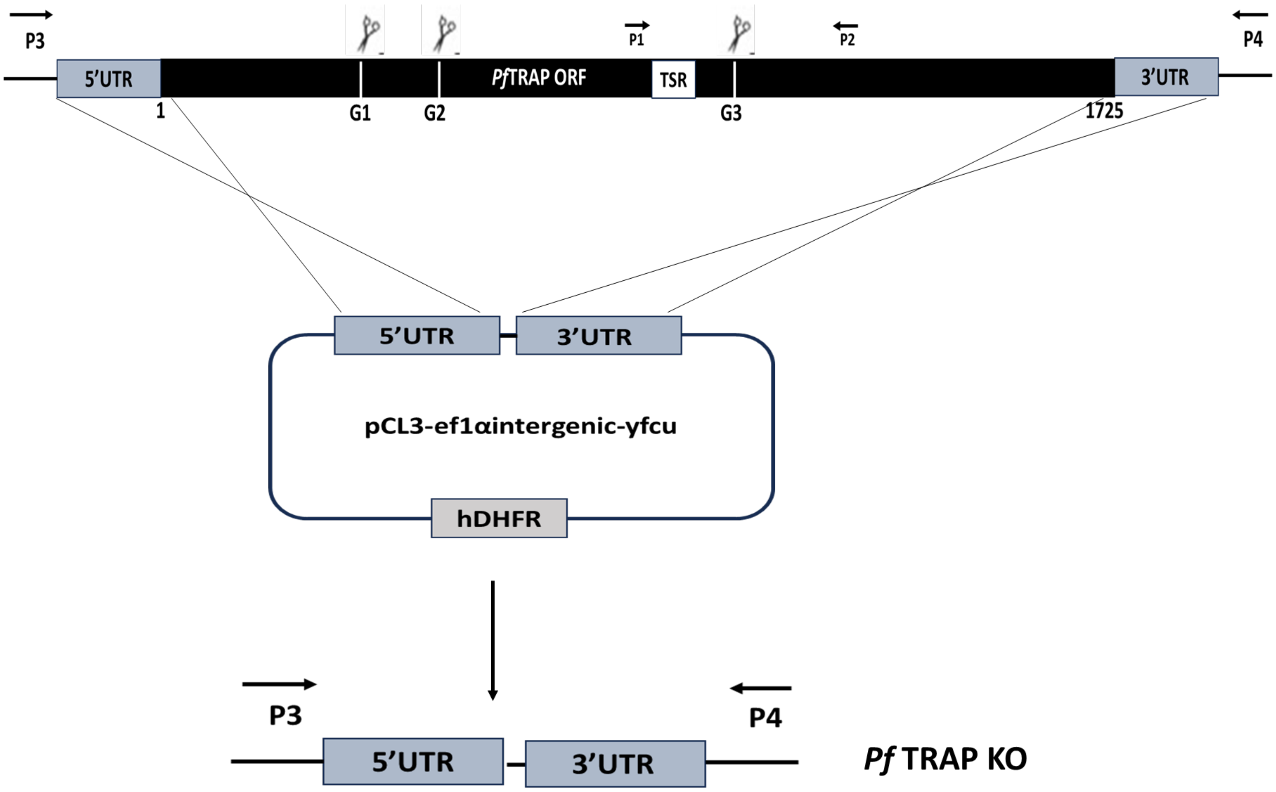


(b)

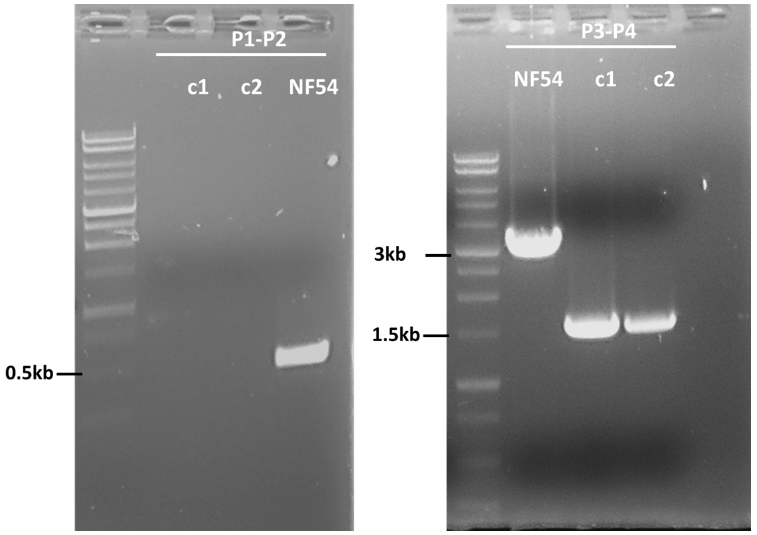


| Primer combination | WT | KO |
|---|---|---|
| P1/P2 | 449bp | No band |
| P3/P4 | 3kb | 1.5kb |

**Figure S2. Schematic of the construct used to generate the *Pf*TRAP— parasite.** (a) Strategy used to generate *Pf*TRAP— parasite using the pcl3-pfc-EF1α-yFCU cloning vector. (b) Genotyping to confirm the deletion of the *Pf*TRAP gene. The table lists the expected PCR fragment size upon deletion of the *Pf*TRAP gene. The sequences of the guides and primers used for this cloning are listed in Table S1.

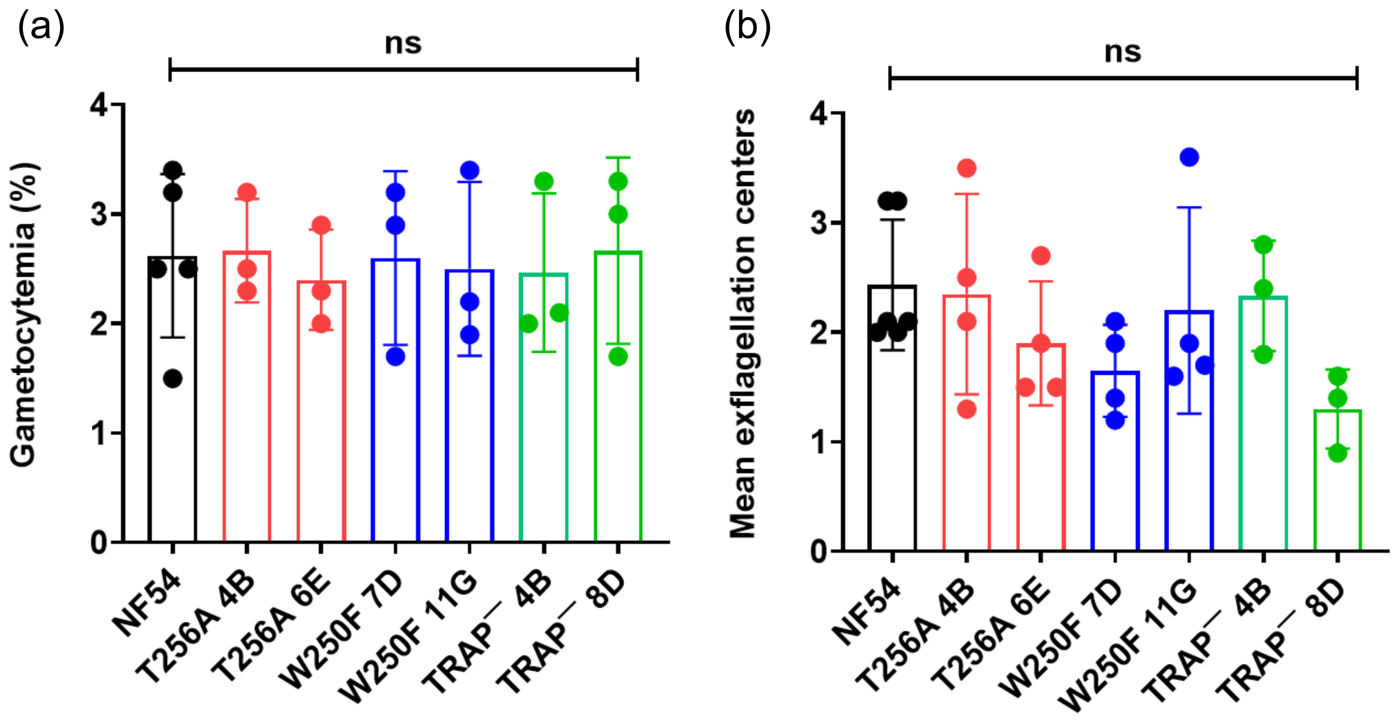


**Figure S3. *Pf* TRAP⁻ and glyco-null mutants exhibit normal sexual stage development.**
(a) Gametocytogenesis was quantified as the prevalence (parasitemia) of transmission-ready stage V gametocytes in a thin smear of infected blood 15 days after initiating gametocytogenesis. (b) Gametogenesis was quantified as the mean number of exflagellation events observed in 10 microscope fields. Neither gametocytogenesis nor gametogenesis were significantly different for transgenic lines compared to wild type NF54. Each point represents an independent experiment. Bars are mean ±S.D. The values represented here are given in **Table S2**.

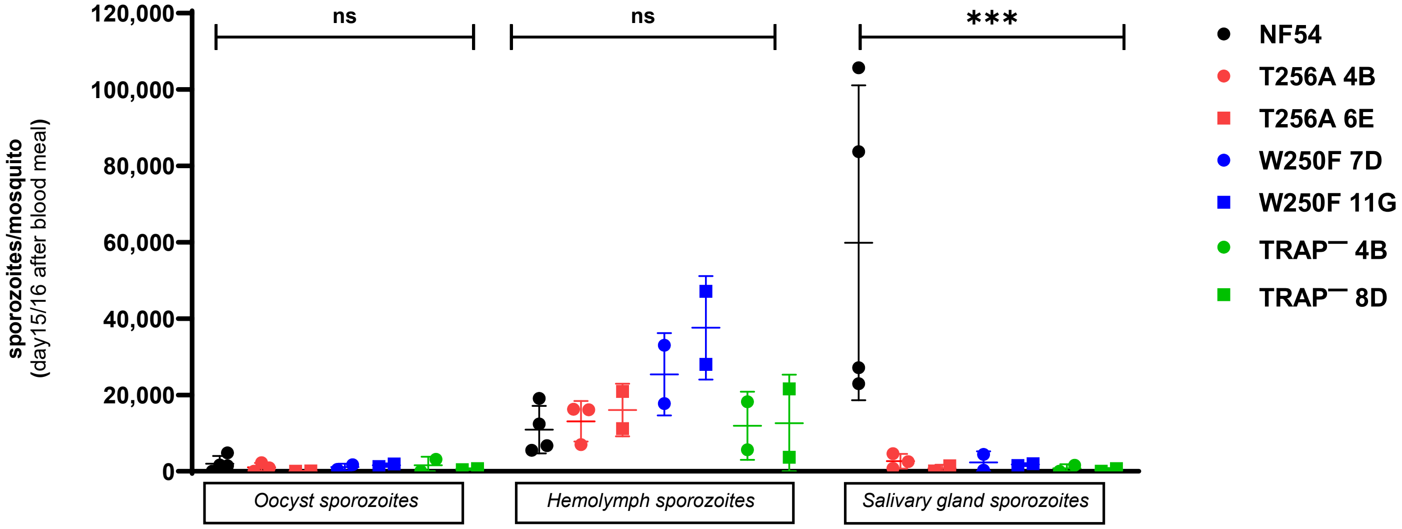


**Figure S4. Distribution of sporozoites in the oocyst, hemolymph, and salivary glands of mosquitoes 15 or 16 days after infectious blood meal.** Sporozoites were collected from oocysts, hemolymph, and salivary glands of the same mosquitoes on day 15 or 16 after infectious blood meal. Data is the mean ±S.D. from independent experiments. Statistical significance was calculated by two-way ANOVA adjusted by Dunnett's multiple comparisons test. The values represented here are given in Supplementary Table S2.

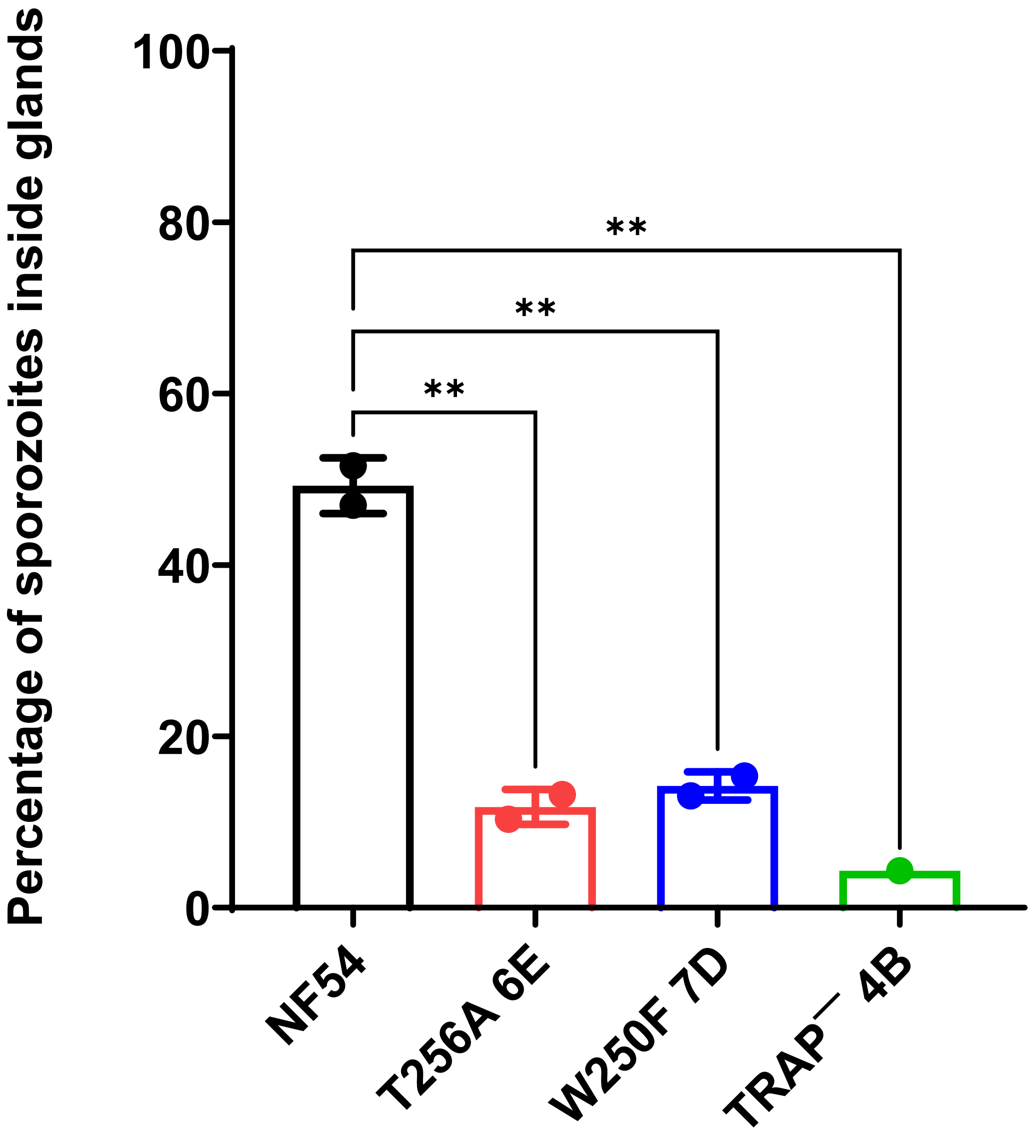


**Figure S5. Trypsinization of salivary glands to remove uninvaded sporozoites.** Salivary glands were dissected from infected mosquitoes and ground to release sporozoites. Shown is the quantity of sporozoites recovered from salivary glands that were first treated with trypsin to remove uninvaded sporozoites as a percentage relative to the number of sporozoites recovered from untreated salivary glands. Bars show the mean ±S.D. from independent experiments. Statistical significance was calculated by one-way ANOVA adjusted by Dunnett's multiple comparison test (** $p < 0.01$). The values represented here are given in Table S2.

Salivary gland sporozoites

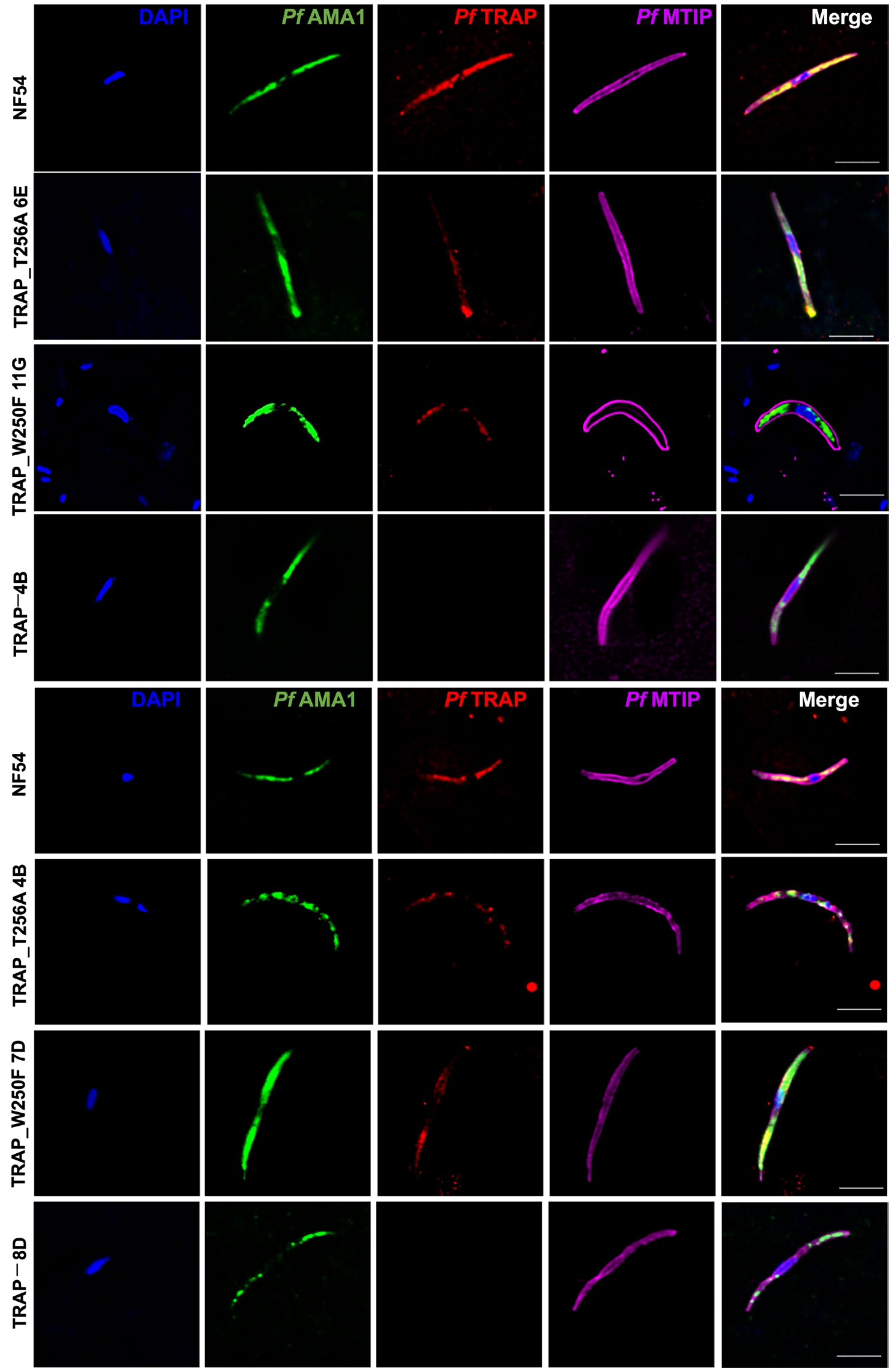


**Figure S6. *Pf*TRAP glyco-null mutants display reduced levels of *Pf*TRAP in salivary gland sporozoites.** Immunofluorescence images of salivary gland sporozoites. *Pf*TRAP (red), AMA-1 (green), MTIP (magenta). Scale bar = 4 µm.

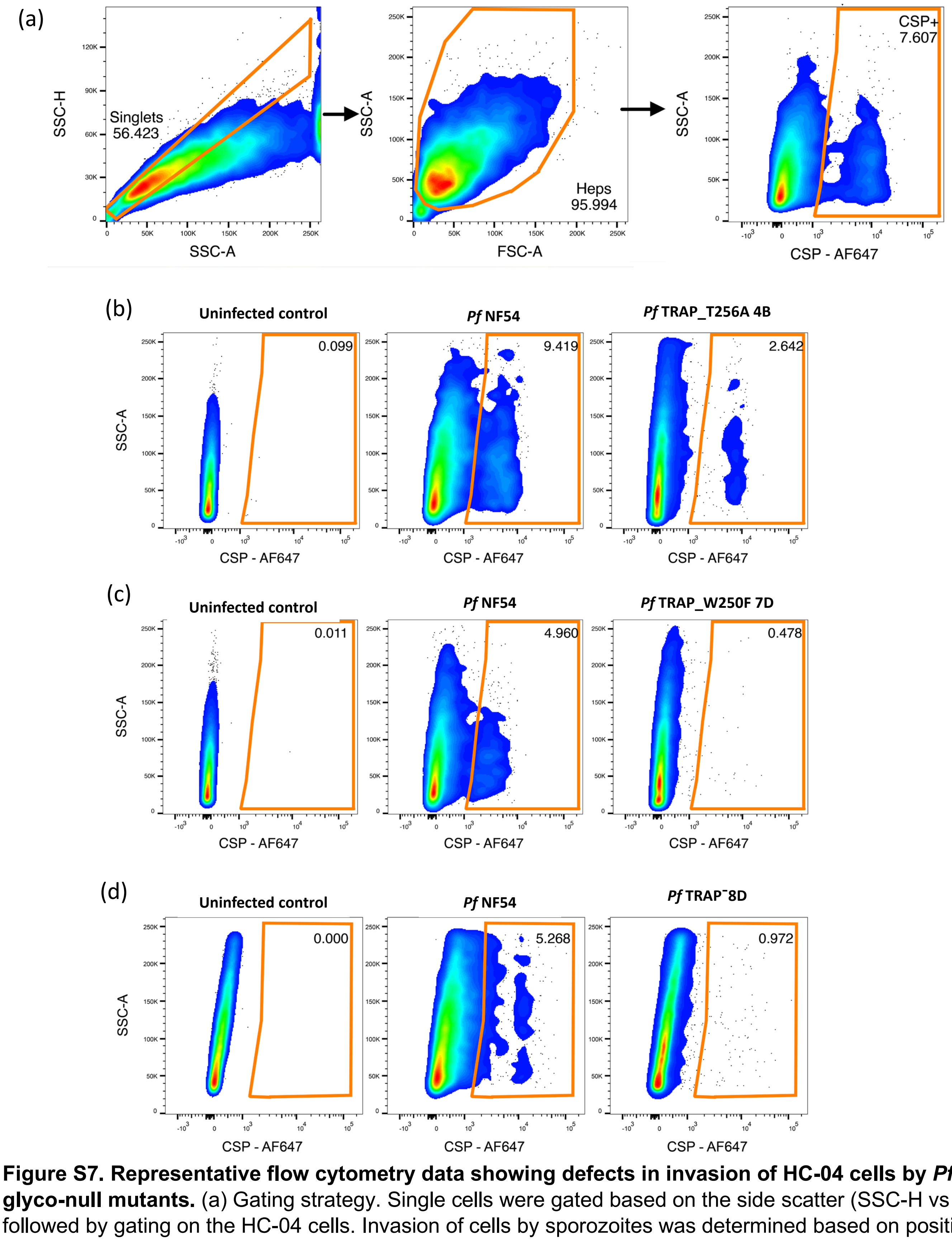


**Figure S7. Representative flow cytometry data showing defects in invasion of HC-04 cells by *Pf*TRAP glyco-null mutants.** (a) Gating strategy. Single cells were gated based on the side scatter (SSC-H vs SSC-A) followed by gating on the HC-04 cells. Invasion of cells by sporozoites was determined based on positive staining for *Pf*CSP. The gating strategies were defined using uninfected control. (b-d) Representative comparisons of sporozoite invasion between wild type NF54 and (b) *Pf*TRAP_T256A (clone 4B), (c) *Pf*TRAP_W250F (clone 7D), (d) *Pf*TRAP— parasite (clone 8D).
