## Supplementary material for "Glycosylation of *Plasmodium falciparum* TRAP supports sporozoite motility and invasion": File S1

### FILE S1. Extracted ion chromatograms and mass spectra of the *Pf*TRAP glycopeptide detected in sporozoites.

All source data for the figures shown here, including raw mass spectrometry data, extracted ion chromatogram values, and complete peak lists of identified peptides, have been deposited to the ProteomeXchange Consortium via the MassIVE partner repository with the dataset identifier PXD064887.
Files can be downloaded from https://massive.ucsd.edu/ProteoSAFe/static/massive.jsp

All MS2 spectra used to identify the *Pf*TRAP glycopeptide, including those shown here, have been assigned Universal Spectrum Identifiers (USI) to enable investigation of the peak annotations. USI viewers are available at

- https://proteomecentral.proteomexchange.org/usi/
- https://proteomecentral.proteomexchange.org/quetzal/
- https://massive.ucsd.edu/ProteoSAFe/usi.jsp

Note that only Quetzal annotates peaks with neutral loss of 120.04 Da arising from cross-ring cleavage of Trp-C-Man.

Mass spectrometry evidence for glycosylation of *Pf*TRAP is provided here for three different samples:

1. **NF54 wild type oocyst sporozoites**
2. ***Pf*TRAP W250F clone 11G oocyst sporozoites**
3. ***Pf*TRAP T256A clone 4B oocyst sporozoites**

The following types of evidence are provided for each sample:

1. **Extracted ion chromatogram.** Xcalibur Qual Browser was used to extract the MS1 signal with respect to retention time from a narrow *m/z* range (0.1–0.3 *m/z*) containing the monoisotopic mass or base peak of the predicted glycoforms in each experiment. The base peak (the second isotope peak) was used for the W250F clone 11G sample to minimize interference from isobaric contaminants. Peaks were only plotted if an MS2 at or near the apex produced a high-quality peptide spectrum match confirming the identity of the glycoform.
2. **Table of detected glycoforms and relative abundance.** For each experiment, the selected ion monitoring (SIM) experiment targeted seven masses corresponding to the 12 possible glycoforms of *Pf*TRAP TSR peptide that contains the three predicted glycosites. These glycoforms encompass all possible permutations of C-Man at $Trp^{247}$ and $Trp^{250}$ and O-Fuc or O-Fuc-Glc at $Thr^{256}$. The table lists all glycoforms detected in the sample along with the retention time and MS1 peak height observed in the extracted ion chromatogram. These abundances are used to calculate the relative abundance of each glycoform reported in Fig. 3 of the manuscript.
3. **Representative mass spectra.** A representative MS2 spectrum was selected for each peak in the extracted ion chromatogram and salient information is provided. These include:
    a. Spectrum information, including the xcorr and Expect scores from the Comet search as well as the *m/z* of the SIM scan window, the targeted precursor *m/z*, and the *m/z* of the detected peptide. Note that, since O-linked glycans are labile in the gas phase, evidence of O-Fuc and O-Fuc-Glc present as neutral loss of 146 and 308 Da, respectively. All MS2 SIM scans windows were offset by 0.5 *m/z* (1.0 Da) in order to center the isolation window on the second isotope peak. Since the center mass of a SIM scan is reported as the precursor mass in the resulting spectrum, all peptides are identified with a mass error of -1.0 Da.
    b. The Universal Spectrum Identifier (USI) for the representative spectrum. This USI can be used to investigate the annotated MS2 spectrum in the USI viewers listed above. These viewers allow the user to investigate alternative peptide sequences, modifications, and modification localizations. Note that the peptide sequence may include {dHex} or {dHex(1)Hex(1)} to account for the unlocalized O-glycan.
    c. The MS2 spectrum annotated by the Lorikeet viewer as implemented in the Trans-Proteomic Pipeline. Neutral loss of $C_4H_8O_4$ from cross-ring cleavage of Trp-C-Man is annotated when present. For data-dependent acquisition data, this viewer automatically displays the parent MS1 spectrum associated with the displayed MS2 spectrum. For the data presented here, there is no parent MS1 spectrum since the data were collected in a data-independent, targeted fashion. For clarity, shown is an MS1 SIM scan that targets the precursor mass and was collected at the same retention time.
    d. A view of the MS1 SIM scan taken from Xcalibur Qual Browser. Data were collected in profile mode with 30,000 resolution. Each MS1 SIM scan was 4 *m/z* wide and centered at +1.5 *m/z* offset from the monoisotopic precursor mass.

#### 1. *Pf*TRAP NF54 oocyst sporozoites
#### Glycopeptide sequence: TASCGVWDEWSPCSVTCGK

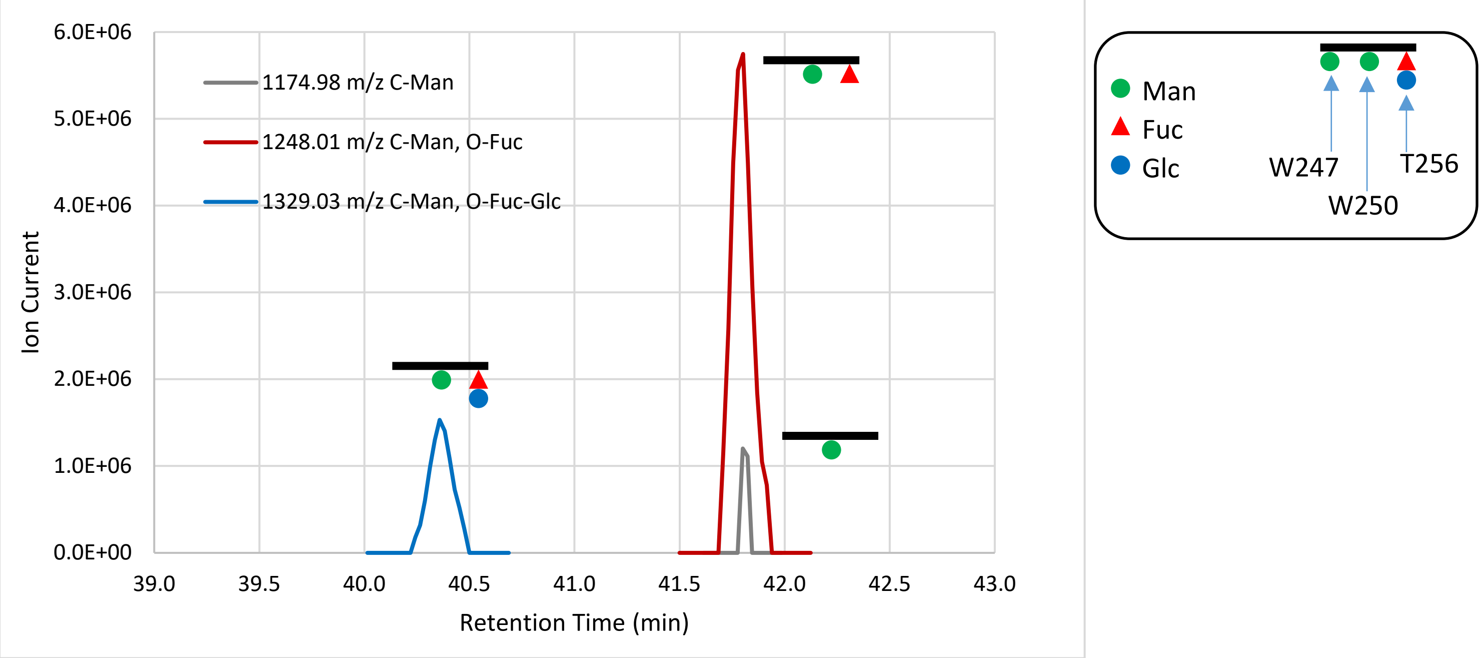


**Fig 1. Extracted ion chromatograms of the detected glycopeptides.** Signal was extracted from the monoisotopic peak for each species. Signal is only presented for peaks that produced MS2 spectra positively identifying the peptide. The peptide species that is C-mannosylated but lacks O-Fuc (1174.98 *m/z*) co-elutes with peptide that is modified with both C-Man and O-Fuc-Glc, indicating that it arises from neutral loss of the O-glycan due to in-source fragmentation while the C-Man moiety remains.

**Table 1. Glycoforms positively identified by MS2 fragment spectra.**

| Glycoform | m/z (z=2) | RT (min) | Peak height | Percent |
|---|---|---|---|---|
| TASCGVW[ ]DEW[ ]SPCSVT[ ]CK | 1093.95 | | | 0.0% |
| TASCGVW[ ]DEW[ ]SPCSVT[Fuc ]CK | 1166.98 | | | 0.0% |
| TASCGVW[Man]DEW[ ]SPCSVT[ ]CK | 1174.98 | | | 0.0% |
| TASCGVW[ ]DEW[Man]SPCSVT[ ]CK | 1174.98 | | | 0.0% |
| TASCGVW[ ]DEW[ ]SPCSVT[FucGlc]CK | 1248.01 | | | 0.0% |
| TASCGVW[Man]DEW[ ]SPCSVT[Fuc ]CK | 1248.01 | | | 0.0% |
| TASCGVW[ ]DEW[Man]SPCSVT[Fuc ]CK | 1248.01 | 41.80 | 5.75E+06 | 79.0% |
| TASCGVW[Man]DEW[Man]SPCSVT[ ]CK | 1256.00 | | | 0.0% |
| TASCGVW[Man]DEW[ ]SPCSVT[FucGlc]CK | 1329.03 | | | 0.0% |
| TASCGVW[ ]DEW[Man]SPCSVT[FucGlc]CK | 1329.03 | 40.36 | 1.53E+06 | 21.0% |
| TASCGVW[Man]DEW[Man]SPCSVT[Fuc ]CK | 1329.03 | | | 0.0% |
| TASCGVW[Man]DEW[Man]SPCSVT[FucGlc]CK | 1410.06 | | | 0.0% |

***Pf*TRAP NF54 wild type oocyst sporozoites** **TASCGVWDEW[Man]SPCSVT[FucGlc]CGK**

| xcorr score | Expect Score | Fragment ions | Retention time (min) | Retention time (sec) | MS2 window center m/z | MS2 window width | Target m/z | Observed Precursor m/z | Peptide m/z | Neutral Loss |
|---|---|---|---|---|---|---|---|---|---|---|
| 4.146 | 2.98E-15 | 23/36 | 40.37 | 2422.20 | 1329.53 | 1.6 | 1329.03 | 1329.03 | 1174.98 | 308.10 |

**USI:** mzspec:PXD064887:Eclipse_2025-02-07_KES_ES903_NF54ooSpz_60min_PRM_250k:scan:25334:{dHex(1)Hex(1)}TASC[Carbamidomethyl]GVWDEW[Hex]SPC[Carbamidomethyl]SVTC[Carbamidomethyl]GK/2

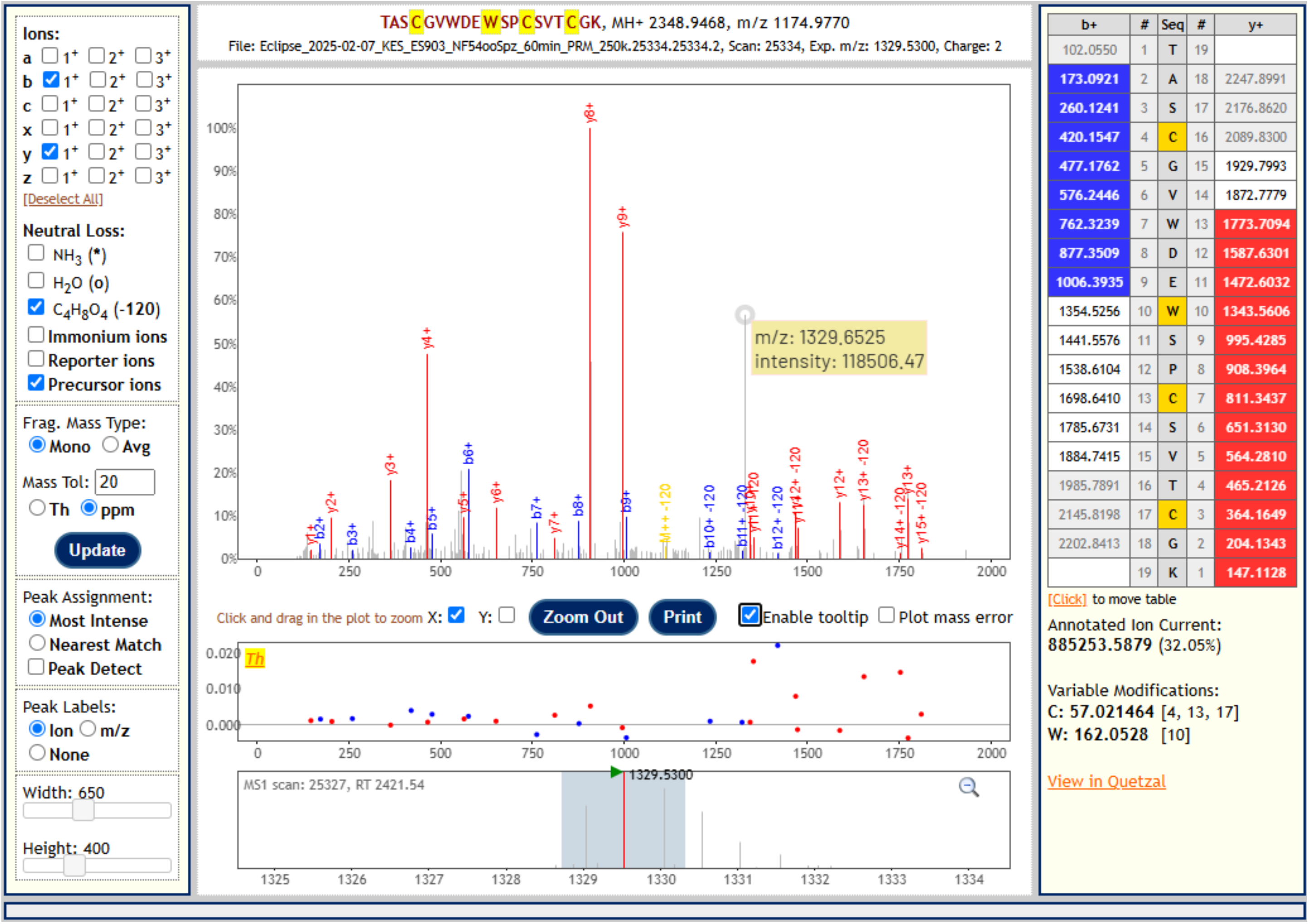

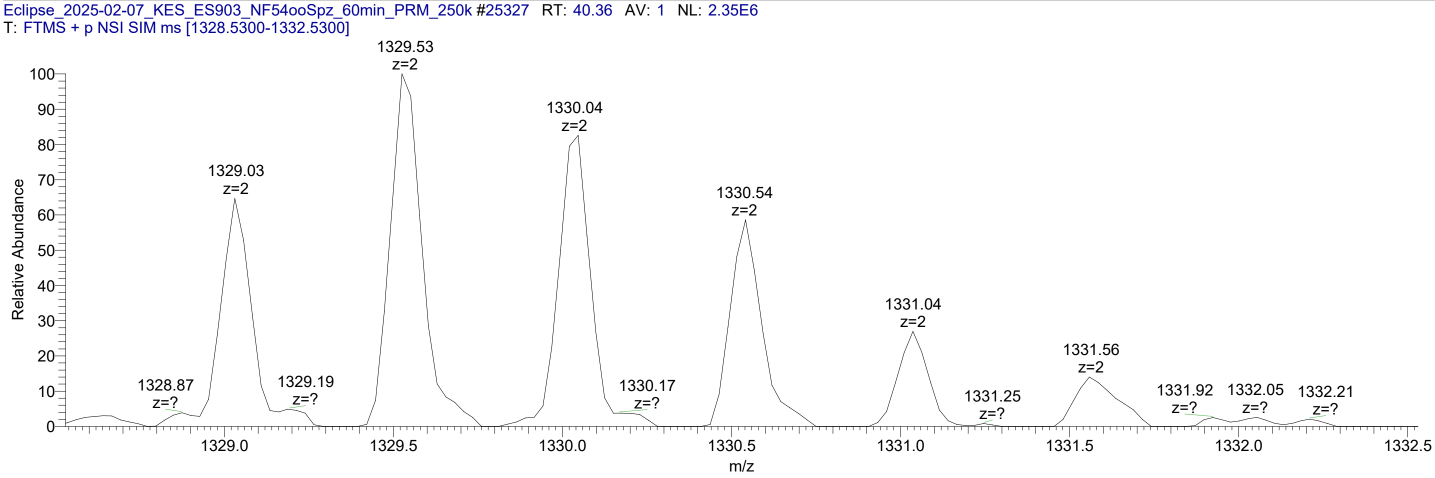

The unannotated peak at 1329.6525 *m/z* in the MS2 spectrum is a singly charged species of unknown origin, likely arising from the interfering species visible in the MS1 spectrum.

***Pf*TRAP NF54 wild type oocyst sporozoites    TASCGVWDEW[Man]SPCSVT[Fuc]CGK**

| xcorr score | Expect Score | Fragment ions | Retention time (min) | Retention time (sec) | MS2 window center m/z | MS2 window width | Target m/z | Observed Precursor m/z | Peptide m/z | Neutral Loss |
|---|---|---|---|---|---|---|---|---|---|---|
| 4.263 | 2.35E-14 | 25/36 | 41.81 | 2508.70 | 1248.51 | 1.6 | 1248.01 | 1248.01 | 1174.98 | 146.06 |

**USI:** mzspec:PXD064887:Eclipse_2025-02-07_KES_ES903_NF54ooSpz_60min_PRM_250k:scan:26262:{dHex}TASC[Carbamidomethyl]GVWDEW[Hex]SPC[Carbamidomethyl]SVTC[Carbamidomethyl]GK/2

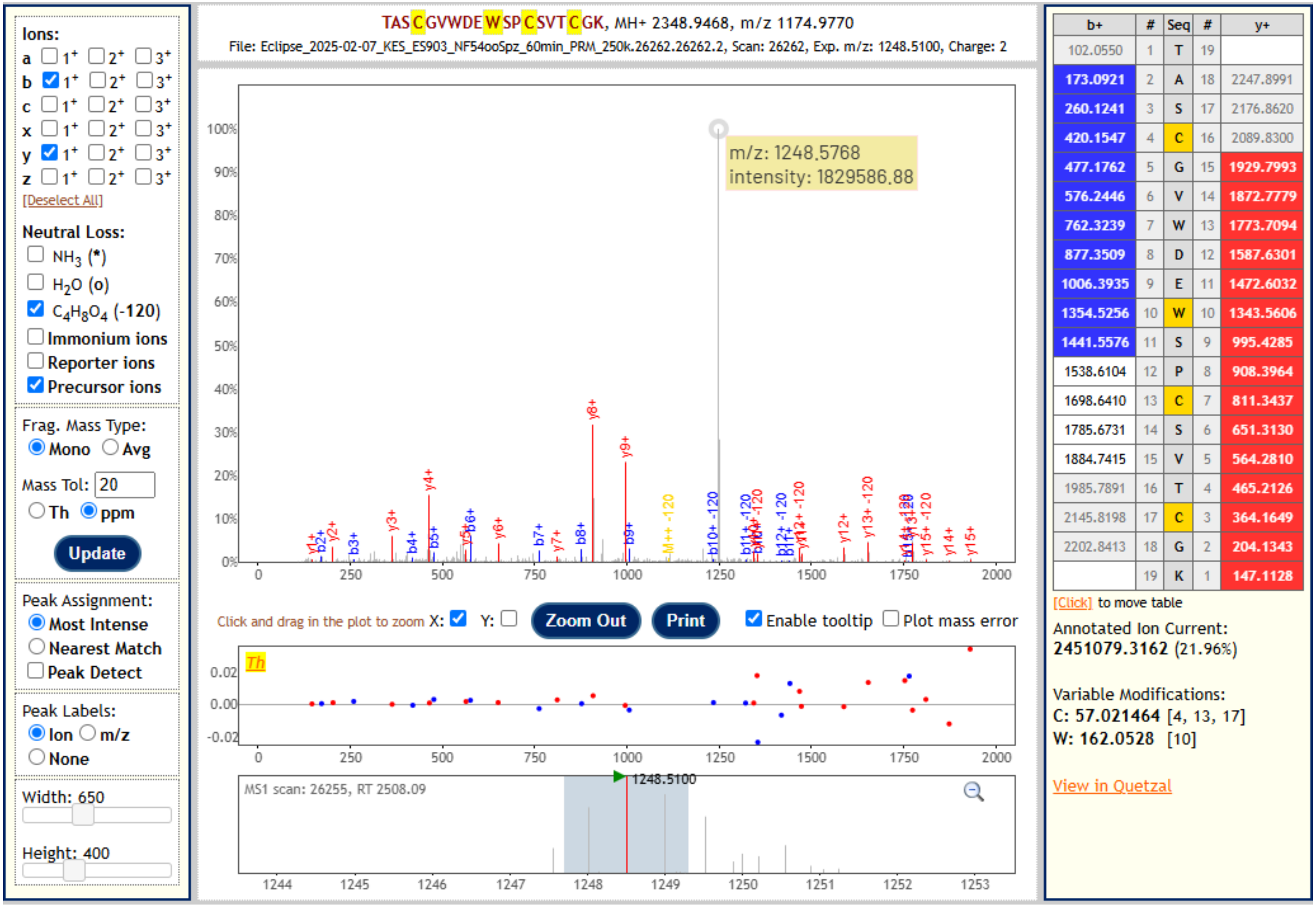


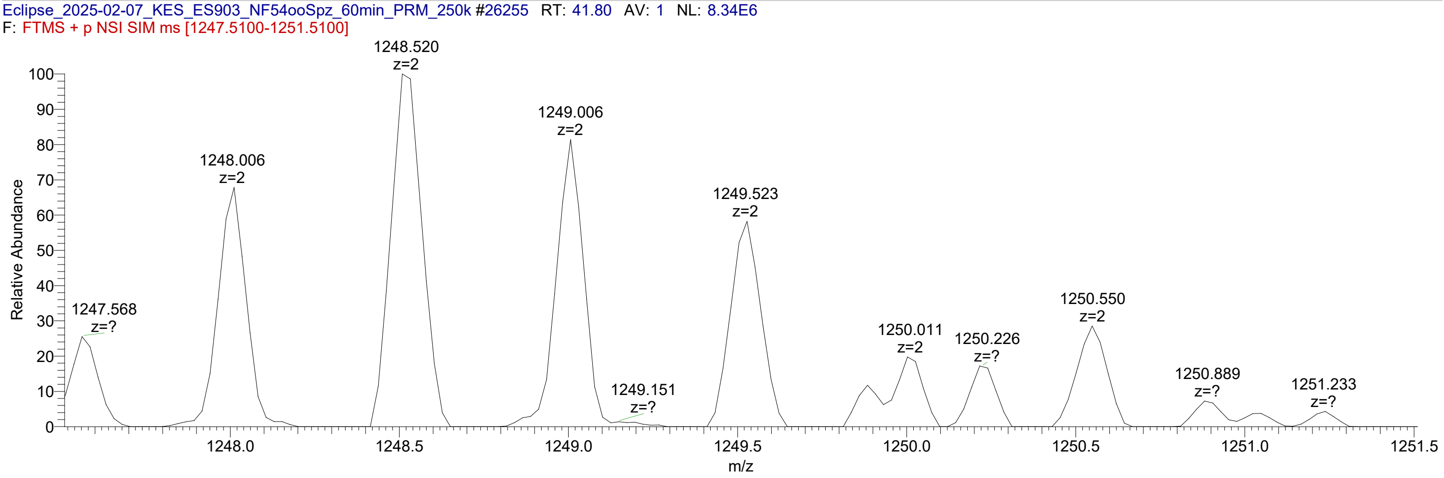


The unannotated peak at 1248.5768 in the MS2 spectrum is a singly charged species of unknown origin, likely arising from the interfering species visible in the MS1 spectrum.

**_Pf_TRAP NF54 wild type oocyst sporozoites          TASCGVWDEW[Man]SPCSVTCGK**

This unmodified peptide species co-elutes with the O-Fuc species, indicating it arises from neutral loss due to in-source fragmentation.

| xcorr score | Expect Score | Fragment ions | Retention time (min) | Retention time (sec) | MS2 window center m/z | MS2 window width | Target m/z | Observed Precursor m/z | Peptide m/z | Neutral Loss |
|---|---|---|---|---|---|---|---|---|---|---|
| 3.053 | 2.05E-11 | 19/36 | 41.79 | 2507.20 | 1175.48 | 1.6 | 1174.98 | 1174.98 | 1174.98 | 0.00 |

**USI:** mzspec:PXD064887:Eclipse_2025-02-07_KES_ES903_NF54ooSpz_60min_PRM_250k:scan:26246:TASC[Carbamidomethyl]GVWDEW[Hex]SPC[Carbamidomethyl]SVTC[Carbamidomethyl]GK/2

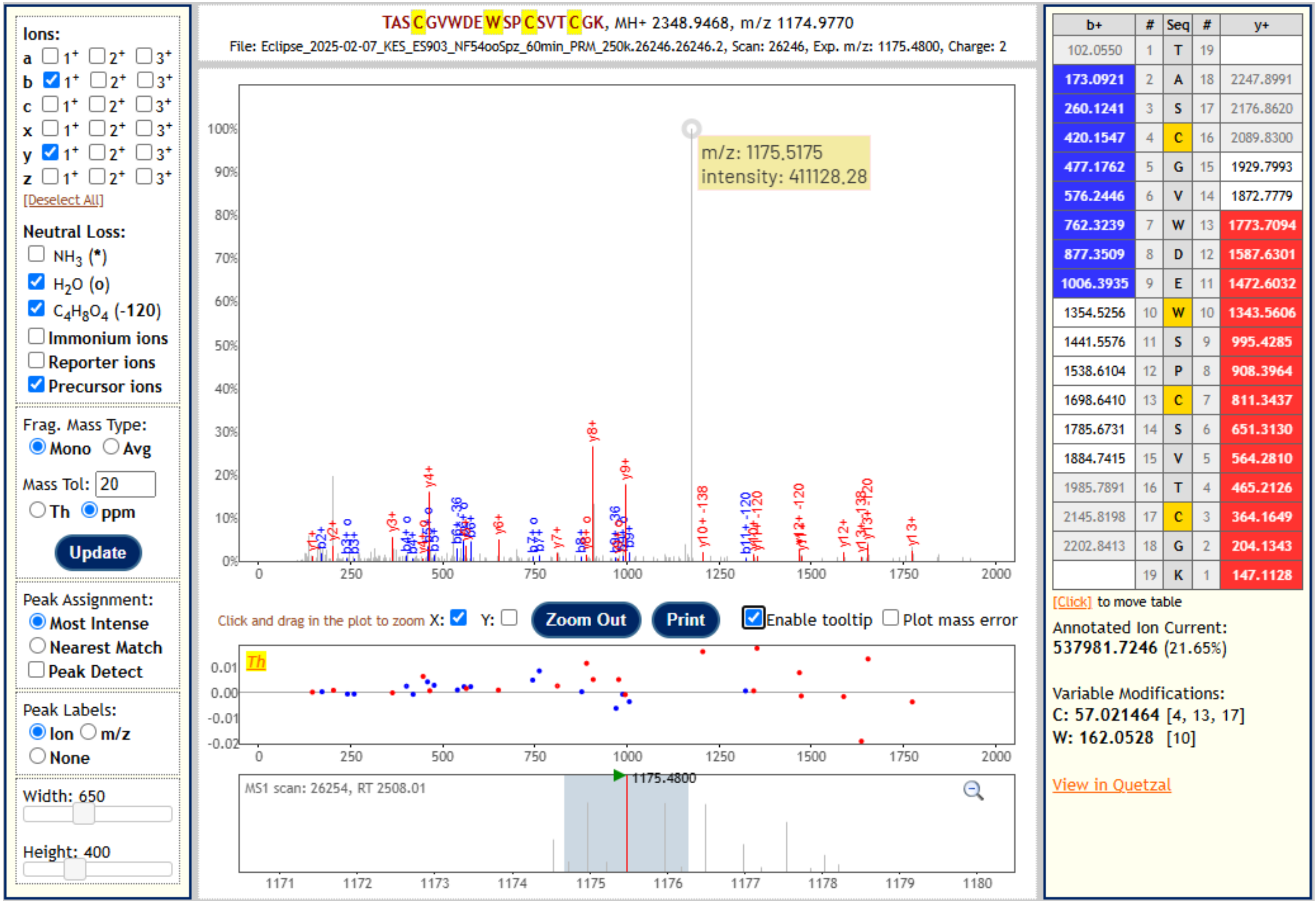


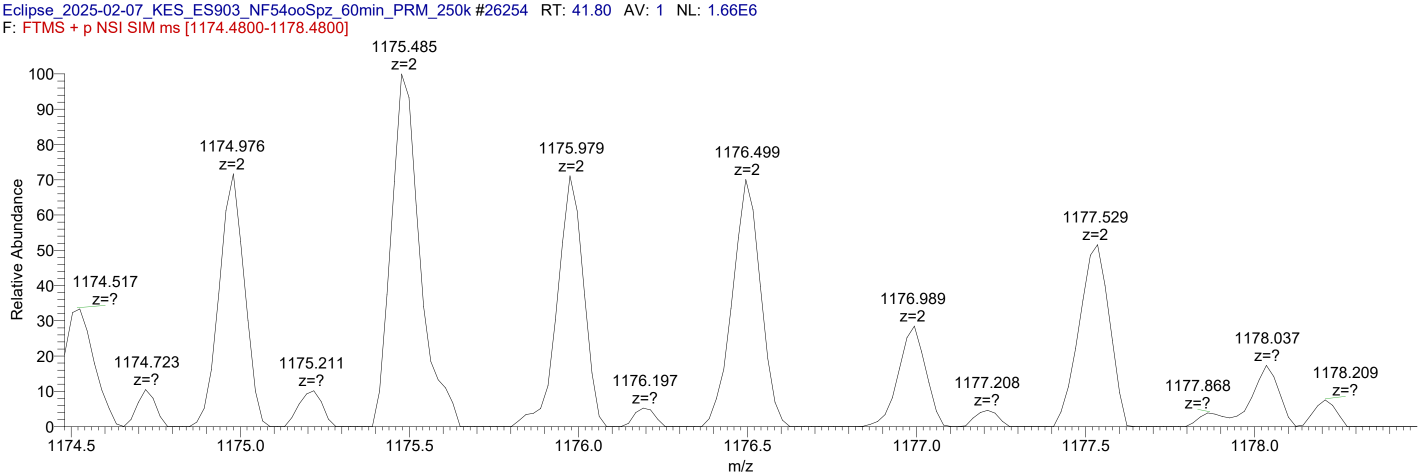


The unannotated peak at 1175.5175 *m/z* in the MS2 spectrum is a singly charged species of unknown origin, likely arising from the interfering species visible in the MS1 spectrum.

#### 2. *Pf*TRAP W250F clone 11G oocyst sporozoites
#### Glycopeptide sequence: TASCGVWDE<u>F</u>SPCSVTCGK

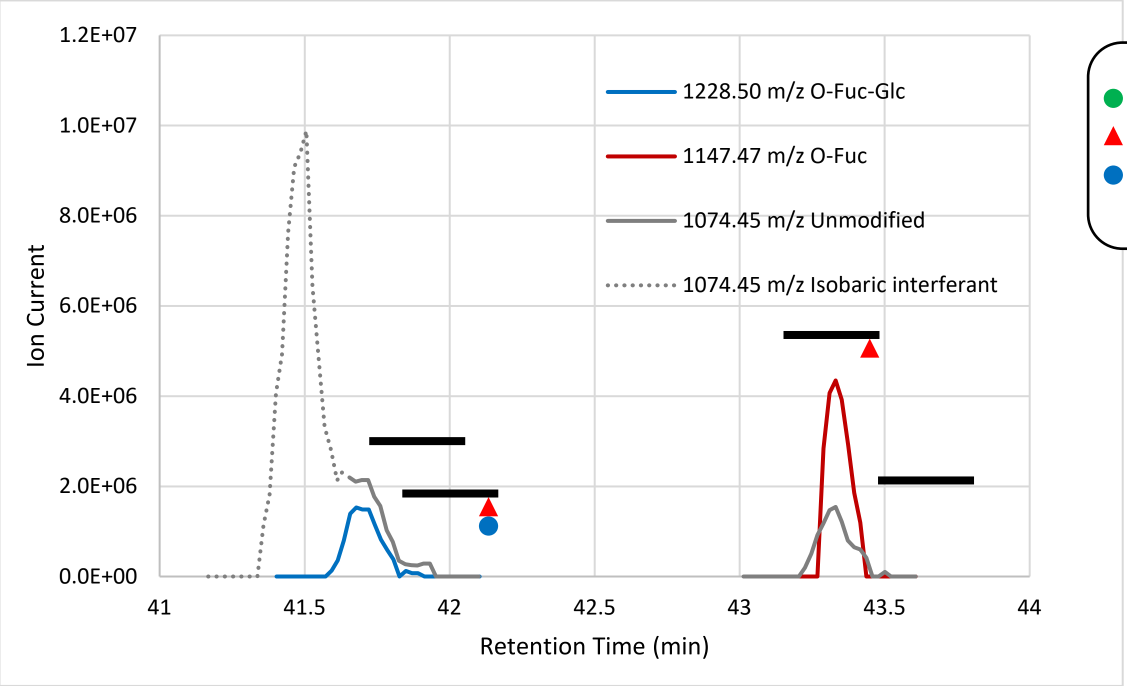


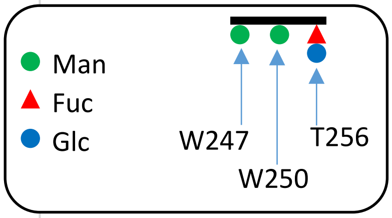


**Fig 2. Extracted ion chromatograms of the detected glycopeptides.** Signal was extracted from the base peak (second isotope peak) for each species. Signal is only presented for peaks that produced MS2 spectra positively identifying the peptide. The unmodified peptide species (1074.45 *m/z*) co-elutes with the O-Fuc and O-Fuc-Glc species, indicating it arises from neutral loss of the O-glycan due to in-source fragmentation. The peak from this species overlaps with an isobaric species eluting at 41.5 min (dashed line), but MS2 spectra distinguish between the two.

**Table 2. Glycoforms positively identified by MS2 fragment spectra.**

| Glycoform | m/z (z=2) | RT (min) | Peak height | Percent |
|---|---|---|---|---|
| TASCGVW[ ]DEF[ ]SPCSVT[ ]CK | 1074.45 | | | 0.0% |
| TASCGVW[ ]DEF[ ]SPCSVT[Fuc ]CK | 1147.47 | 43.33 | 4.35E+06 | 74.0% |
| TASCGVW[Man]DEF[ ]SPCSVT[ ]CK | 1155.47 | | | 0.0% |
| TASCGVW[ ]DEF[Man]SPCSVT[ ]CK | 1155.47 | | | 0.0% |
| TASCGVW[ ]DEF[ ]SPCSVT[FucGlc]CK | 1228.50 | 41.68 | 1.53E+06 | 26.0% |
| TASCGVW[Man]DEF[ ]SPCSVT[Fuc ]CK | 1228.50 | | | 0.0% |
| TASCGVW[ ]DEF[Man]SPCSVT[Fuc ]CK | 1228.50 | | | 0.0% |
| TASCGVW[Man]DEF[Man]SPCSVT[ ]CK | 1236.50 | | | 0.0% |
| TASCGVW[Man]DEF[ ]SPCSVT[FucGlc]CK | 1309.53 | | | 0.0% |
| TASCGVW[ ]DEF[Man]SPCSVT[FucGlc]CK | 1309.53 | | | 0.0% |
| TASCGVW[Man]DEF[Man]SPCSVT[Fuc ]CK | 1309.53 | | | 0.0% |
| TASCGVW[Man]DEF[Man]SPCSVT[FucGlc]CK | 1390.55 | | | 0.0% |

***Pf*TRAP W250F clone 11G oocyst sporozoites** **TASCGVWDEFSPCSVT[Fuc]CGK**

| xcorr score | Expect Score | Fragment ions | Retention Time (min) | Retention Time (sec) | MS2 window center m/z | MS2 window width | Target m/z | Observed Precursor m/z | Peptide m/z | Neutral Loss |
|---|---|---|---|---|---|---|---|---|---|---|
| 3.493 | 4.65E-11 | 21/36 | 43.30 | 2597.90 | 1147.97 | 1.6 | 1147.47 | 1147.48 | 1074.45 | 146.04 |

**USI:** mzspec:PXD064887:isbECLIPSE_2024-10-10_KES_ES903_60minPRM_ooSpz_TRAP-W250F-11G_500k:scan:28240:{dHex}TASC[Carbamidomethyl]GVWDEFSPC[Carbamidomethyl]SVTC[Carbamidomethyl]GK/2

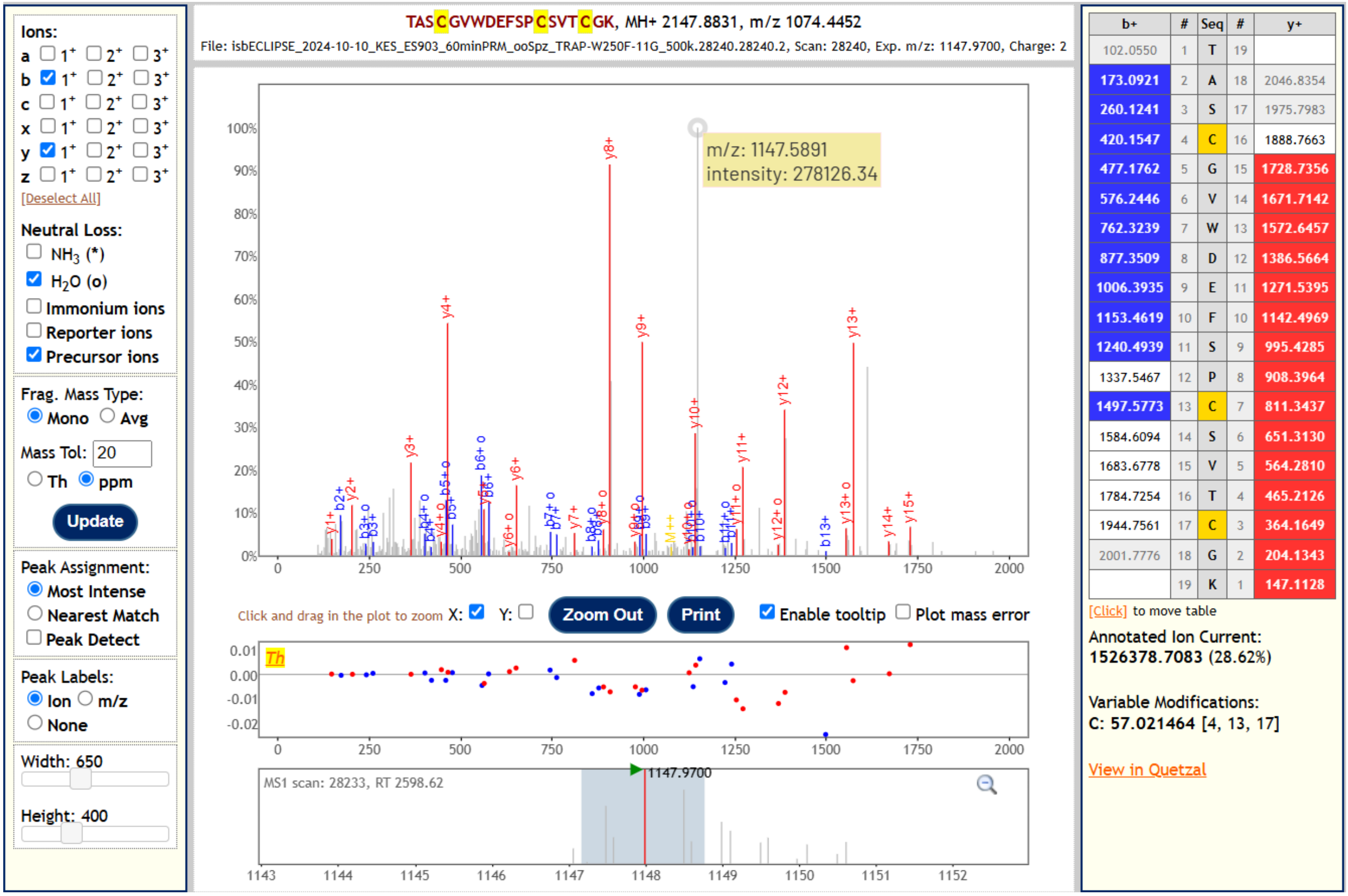


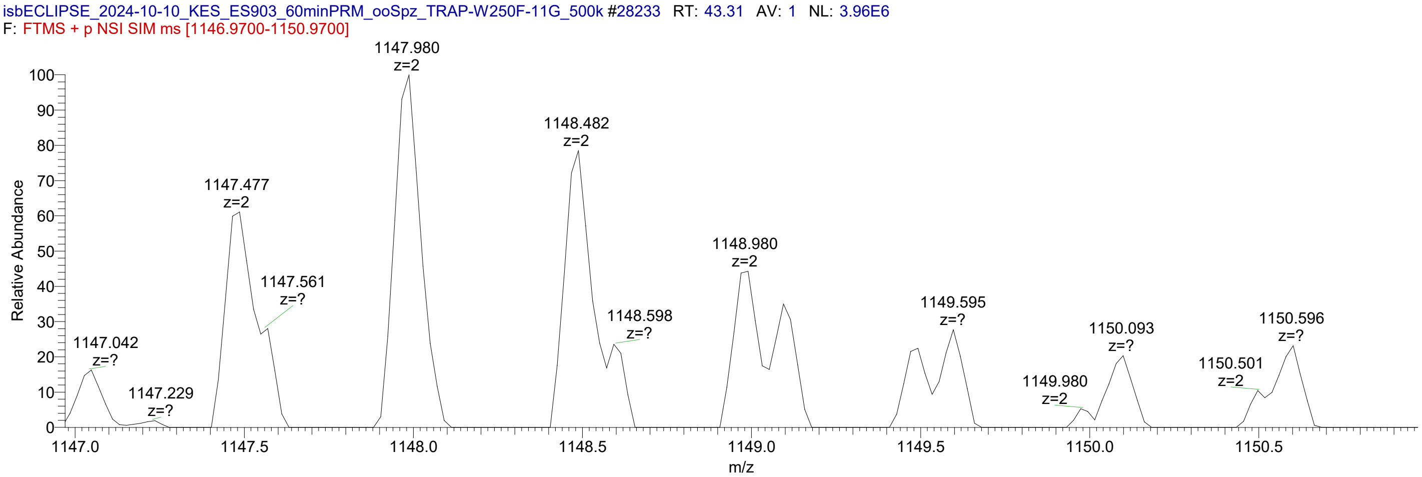


The unannotated peaks at 1147.5891 *m/z* and 1610.8632 *m/z* in the MS2 spectrum are singly charged species of unknown origin, likely arising from the interfering species visible in the MS1 spectrum and resulting in a chimeric spectrum.

***Pf*TRAP W250F clone 11G oocyst sporozoites** **TASCGVWDEFSPCSVTCGK**

This unmodified peptide species co-elutes with the O-Fuc species, indicating it arises from neutral loss due to in-source fragmentation.

| xcorr score | Expect Score | Fragment ions | Retention Time (min) | Retention Time (sec) | MS2 window center m/z | MS2 window width | Target m/z | Observed Precursor m/z | Peptide m/z | Neutral Loss |
|---|---|---|---|---|---|---|---|---|---|---|
| 2.632 | 5.48E-11 | 20/36 | 43.30 | 2597.90 | 1074.95 | 1.6 | 1074.45 | 1074.45 | 1074.45 | 0.00 |

**USI:** mzspec:PXD064887:isbECLIPSE_2024-10-10_KES_ES903_60minPRM_ooSpz_TRAP-W250F-11G_500k:scan:28224:TASC[Carbamidomethyl]GVWDEFSPC[Carbamidomethyl]SVTC[Carbamidomethyl]GK/2

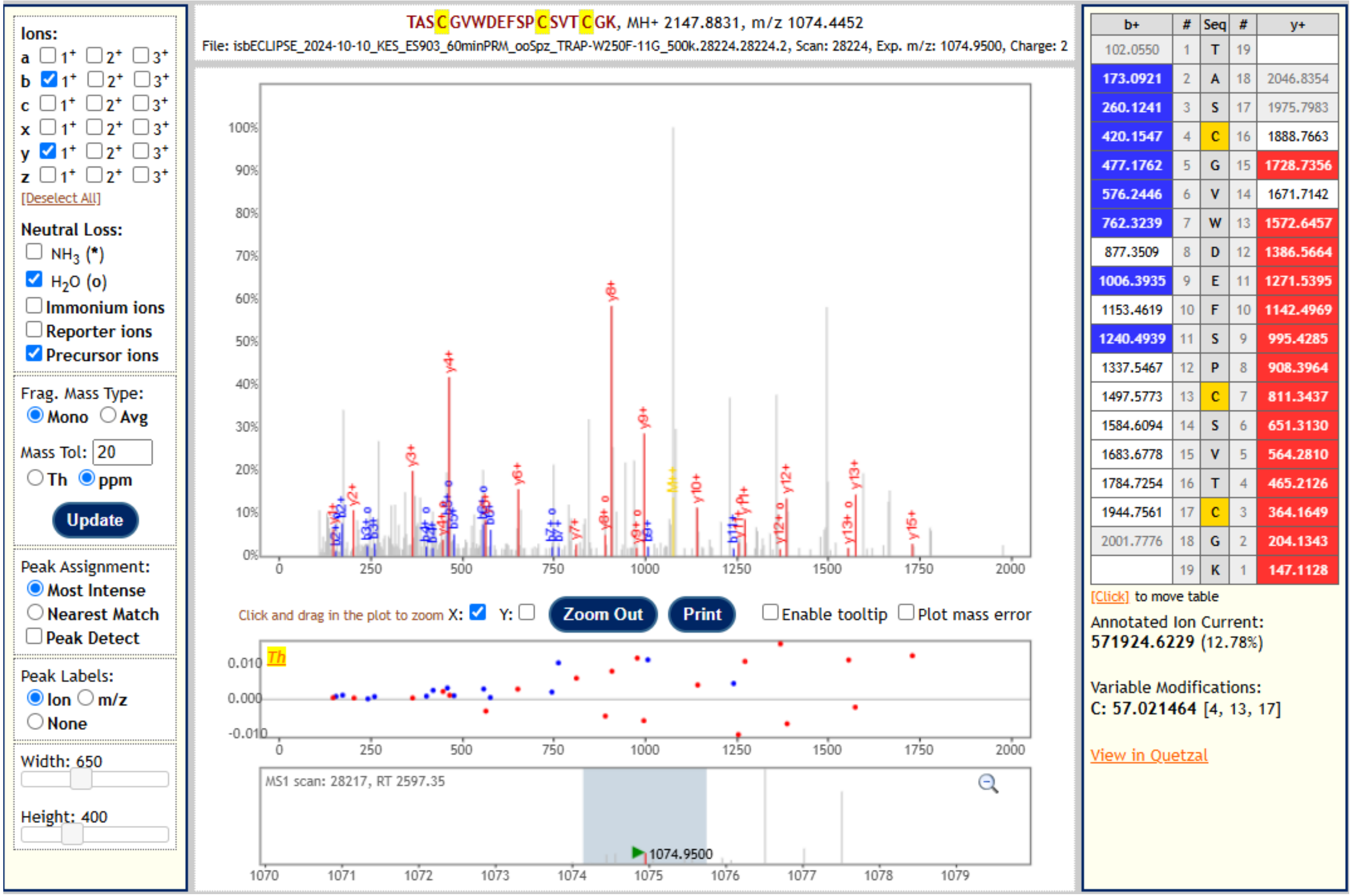


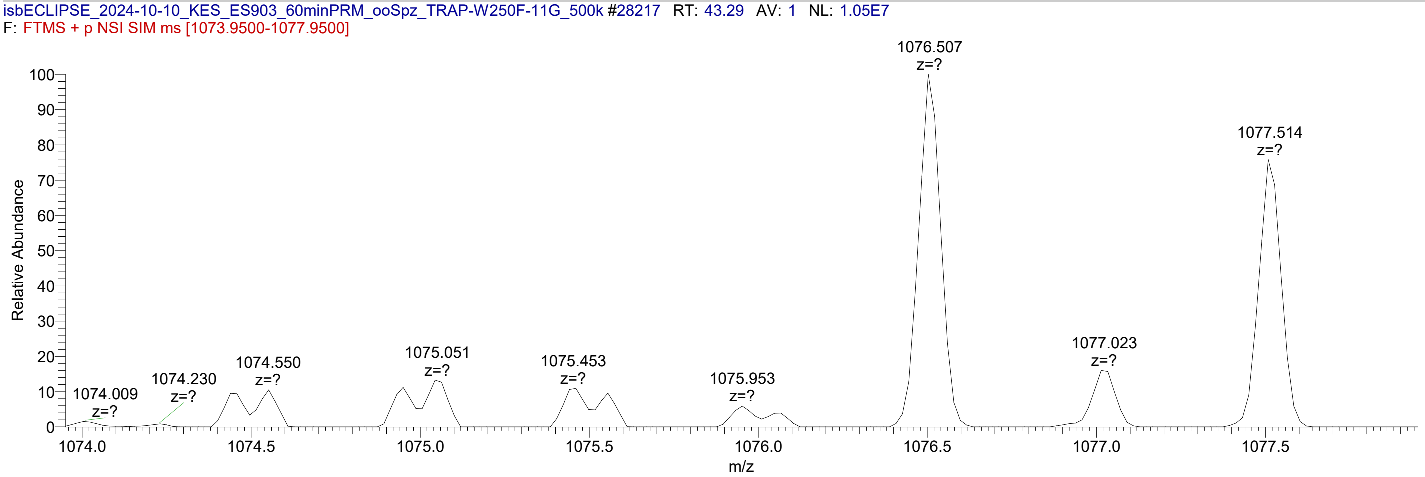


The MS2 spectrum is chimeric, likely due to the interfering species visible in the MS1 spectrum.

***Pf*TRAP W250F clone 11G oocyst sporozoites     TASCGVWDEFSPCSVT[FucGlc]CGK**

| xcorr score | Expect Score | Fragment ions | Retention Time (min) | Retention Time (sec) | MS2 window center m/z | MS2 window width | Target m/z | Observed Precursor m/z | Peptide m/z | Neutral Loss |
|---|---|---|---|---|---|---|---|---|---|---|
| 3.703 | 1.06E-12 | 20/36 | 41.69 | 2501.30 | 1229.00 | 1.6 | 1228.50 | 1228.51 | 1074.45 | 308.10 |

**USI:** mzspec:PXD064887:isbECLIPSE_2024-10-10_KES_ES903_60minPRM_ooSpz_TRAP-W250F-11G_500k:scan:27087:{dHex(1)Hex(1)}TASC[Carbamidomethyl]GVWDEFSPC[Carbamidomethyl]SVTC[Carbamidomethyl]GK/2

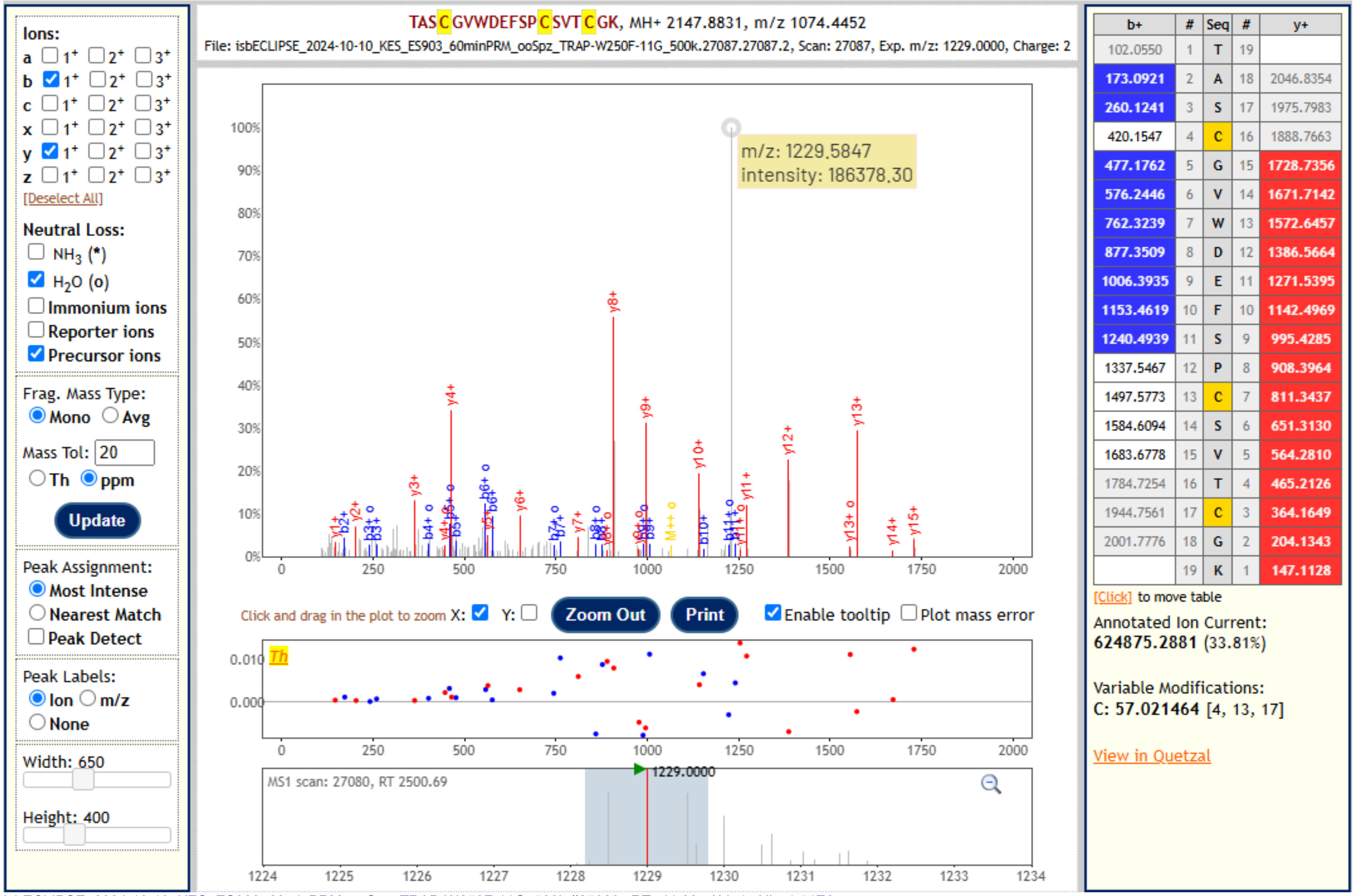


| b+ | # | Seq | # | y+ |
|---|---|---|---|---|
| 102.0550 | 1 | T | 19 | |
| 173.0921 | 2 | A | 18 | 2046.8354 |
| 260.1241 | 3 | S | 17 | 1975.7983 |
| 420.1547 | 4 | C | 16 | 1888.7663 |
| 477.1762 | 5 | G | 15 | 1728.7356 |
| 576.2446 | 6 | V | 14 | 1671.7142 |
| 762.3239 | 7 | W | 13 | 1572.6457 |
| 877.3509 | 8 | D | 12 | 1386.5664 |
| 1006.3935 | 9 | E | 11 | 1271.5395 |
| 1153.4619 | 10 | F | 10 | 1142.4969 |
| 1240.4939 | 11 | S | 9 | 995.4285 |
| 1337.5467 | 12 | P | 8 | 908.3964 |
| 1497.5773 | 13 | C | 7 | 811.3437 |
| 1584.6094 | 14 | S | 6 | 651.3130 |
| 1683.6778 | 15 | V | 5 | 564.2810 |
| 1784.7254 | 16 | T | 4 | 465.2126 |
| 1944.7561 | 17 | C | 3 | 364.1649 |
| 2001.7776 | 18 | G | 2 | 204.1343 |
| | 19 | K | 1 | 147.1128 |

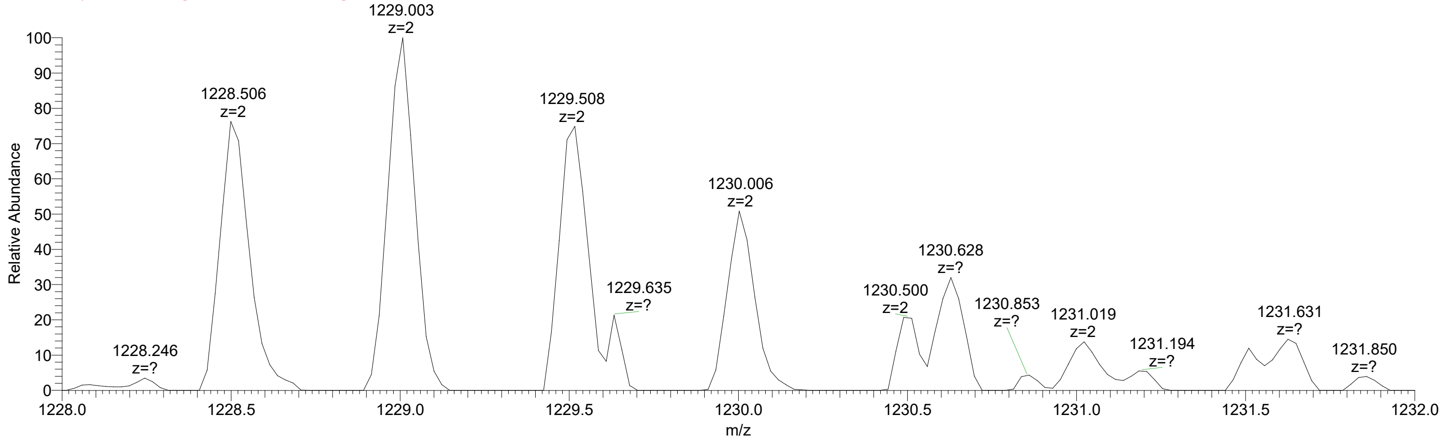


The unlabeled peak 1229.5847 is a singly charged species of unknown origin, likely arising from the interfering species visible in the MS1 spectrum, e.g., the 1229.635 *m/z* species.

***Pf*TRAP W250F clone 11G oocyst sporozoites  TASCGVWDEFSPCSVTCGK**

This unmodified peptide species co-elutes with the O-Fuc-Glc species, indicating it arises from neutral loss due to in-source fragmentation.

| xcorr score | Expect Score | Fragment ions | Retention Time (min) | Retention Time (sec) | MS2 window center m/z | MS2 window width | Target m/z | Observed Precursor m/z | Peptide m/z | Neutral Loss |
|---|---|---|---|---|---|---|---|---|---|---|
| 3.317 | 4.80E-09 | 16/36 | 41.69 | 2501.10 | 1074.95 | 1.6 | 1074.45 | 1074.45 | 1074.45 | 0.00 |

**USI:** mzspec:PXD064887:isbECLIPSE_2024-10-10_KES_ES903_60minPRM_ooSpz_TRAP-W250F-11G_500k:scan:27084:TASC[Carbamidomethyl]GVWDEFSPC[Carbamidomethyl]SVTC[Carbamidomethyl]GK/2

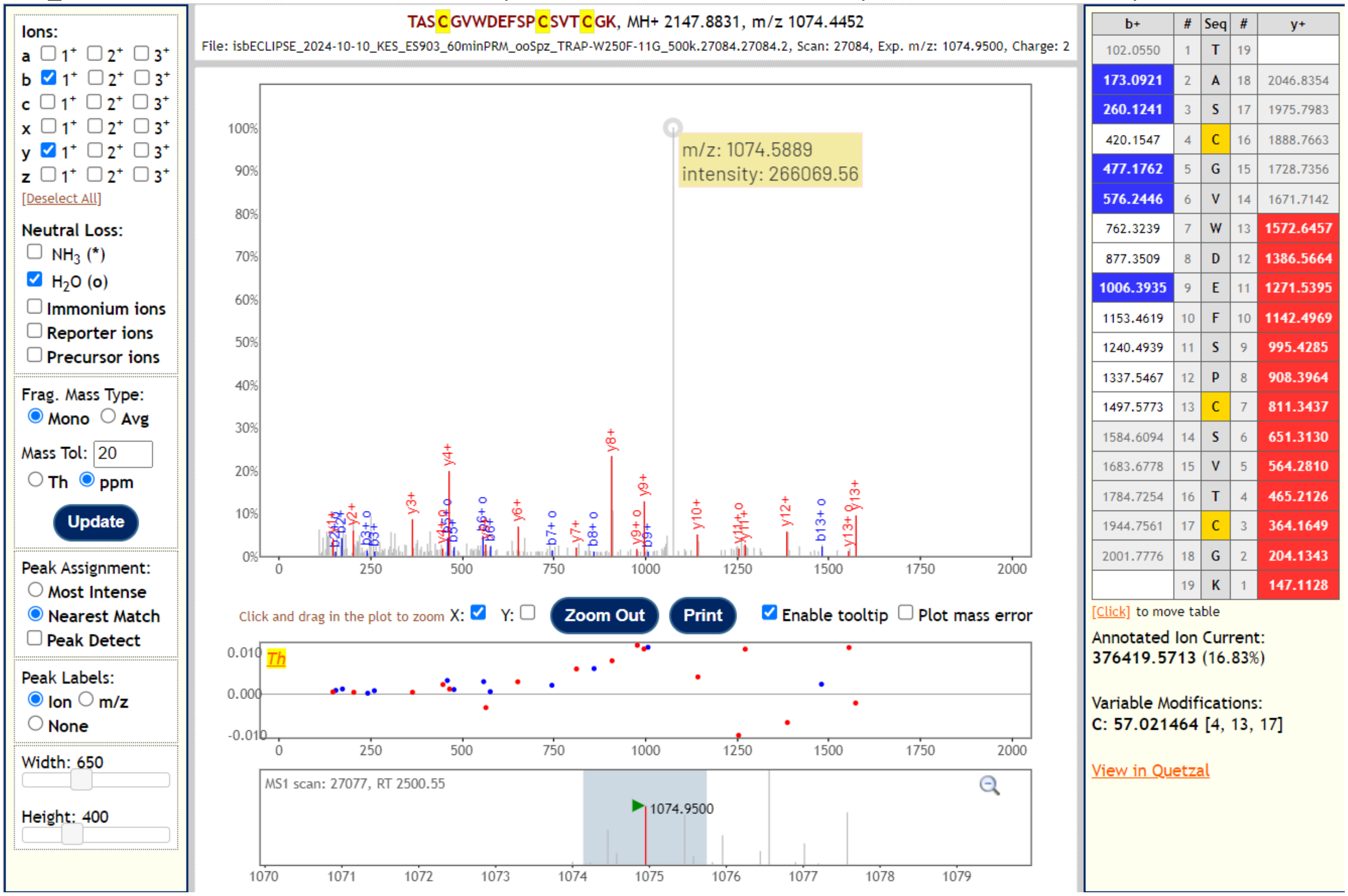


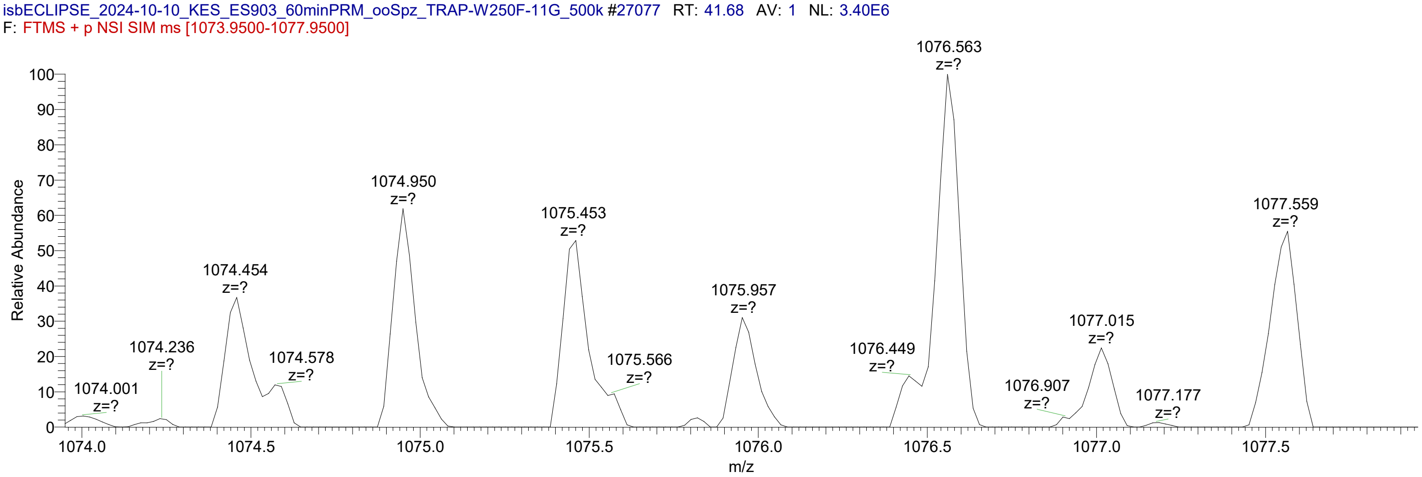


The unlabeled peak 1074.5889 is a singly charged species of unknown origin, likely arising from the interfering species visible in the MS1 spectrum, e.g., the 1074.578 *m/z* species.

#### 3. *Pf*TRAP T256A clone 4B oocyst sporozoites
#### Glycopeptide sequence: TASCGVWDEWPCSVACGK

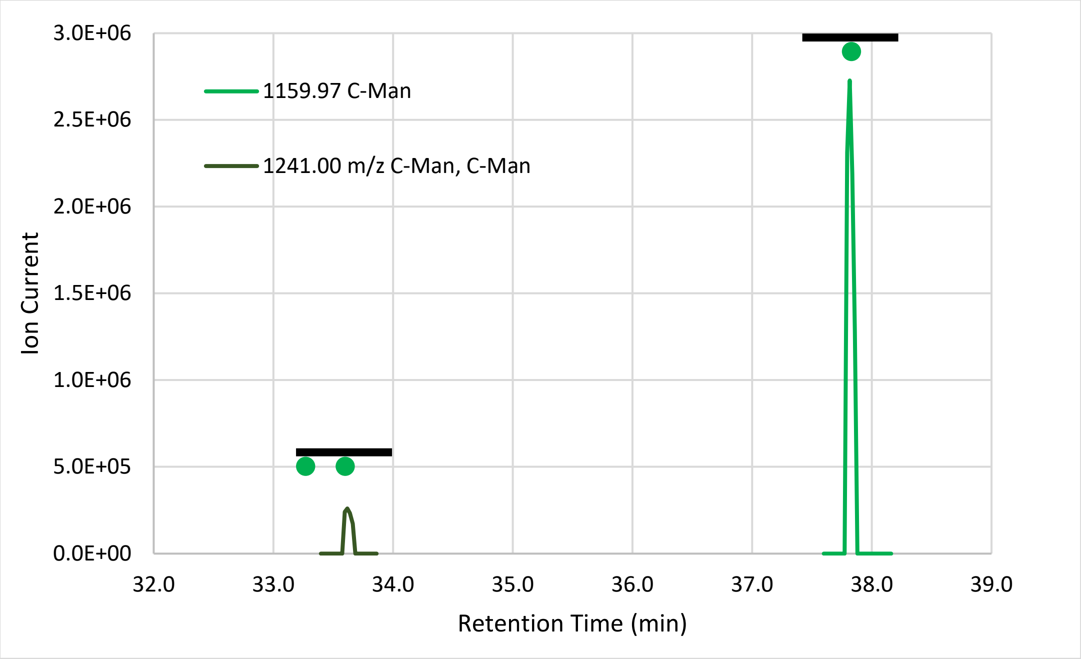


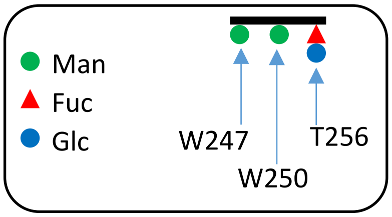


**Fig 3. Extracted ion chromatograms of the detected glycopeptides.** Signal was extracted from the monoisotopic peak for each species. Signal is only presented for peaks that produced MS2 spectra positively identifying the peptide.

**Table 3. Glycoforms positively identified by MS2 fragment spectra.**

| Glycoform | m/z (z=2) | RT (min) | Peak height | Percent |
|---|---|---|---|---|
| TASCGVW[ ]DEW[ ]SPCSVA[ ]CK | 1078.95 | | | 0.0% |
| TASCGVW[ ]DEW[ ]SPCSVA[Fuc ]CK | 1151.97 | | | 0.0% |
| TASCGVW[Man]DEW[ ]SPCSVA[ ]CK | 1159.97 | | | 0.0% |
| TASCGVW[ ]DEW[Man]SPCSVA[ ]CK | 1159.97 | 37.82 | 2.73E+06 | 91.3% |
| TASCGVW[ ]DEW[ ]SPCSVA[FucGlc]CK | 1233.00 | | | 0.0% |
| TASCGVW[Man]DEW[ ]SPCSVA[Fuc ]CK | 1233.00 | | | 0.0% |
| TASCGVW[ ]DEW[Man]SPCSVA[Fuc ]CK | 1233.00 | | | 0.0% |
| TASCGVW[Man]DEW[Man]SPCSVA[ ]CK | 1241.00 | 33.62 | 2.60E+05 | 8.7% |
| TASCGVW[Man]DEW[ ]SPCSVA[FucGlc]CK | 1314.03 | | | 0.0% |
| TASCGVW[ ]DEW[Man]SPCSVA[FucGlc]CK | 1314.03 | | | 0.0% |
| TASCGVW[Man]DEW[Man]SPCSVA[Fuc ]CK | 1314.03 | | | 0.0% |
| TASCGVW[Man]DEW[Man]SPCSVA[FucGlc]CK | 1395.05 | | | 0.0% |

***Pf*TRAP T256A clone 4B oocyst sporozoites** **TASCGVWDEW[Man]SPCSVACGK**

| xcorr score | Expect Score | Fragment ions | Retention time (min) | Retention time (sec) | MS2 window center m/z | MS2 window width | Target m/z | Observed Precursor m/z | Peptide m/z | Neutral Loss |
|---|---|---|---|---|---|---|---|---|---|---|
| 4.294 | 9.97E-15 | 21/36 | 37.83 | 2269.5 | 1160.47 | 1.6 | 1159.97 | 1159.97 | 1159.97 | 0.00 |

**USI:** mzspec:PXD064887:isbECLIPSE_2024-10-10_KES_ES903_60minPRM_ooSpz_TRAP-T256A-4B_600k:scan:24221:TASC[Carbamidomethyl]GVWDEW[Hex]SPC[Carbamidomethyl]SVAC[Carbamidomethyl]GK/2

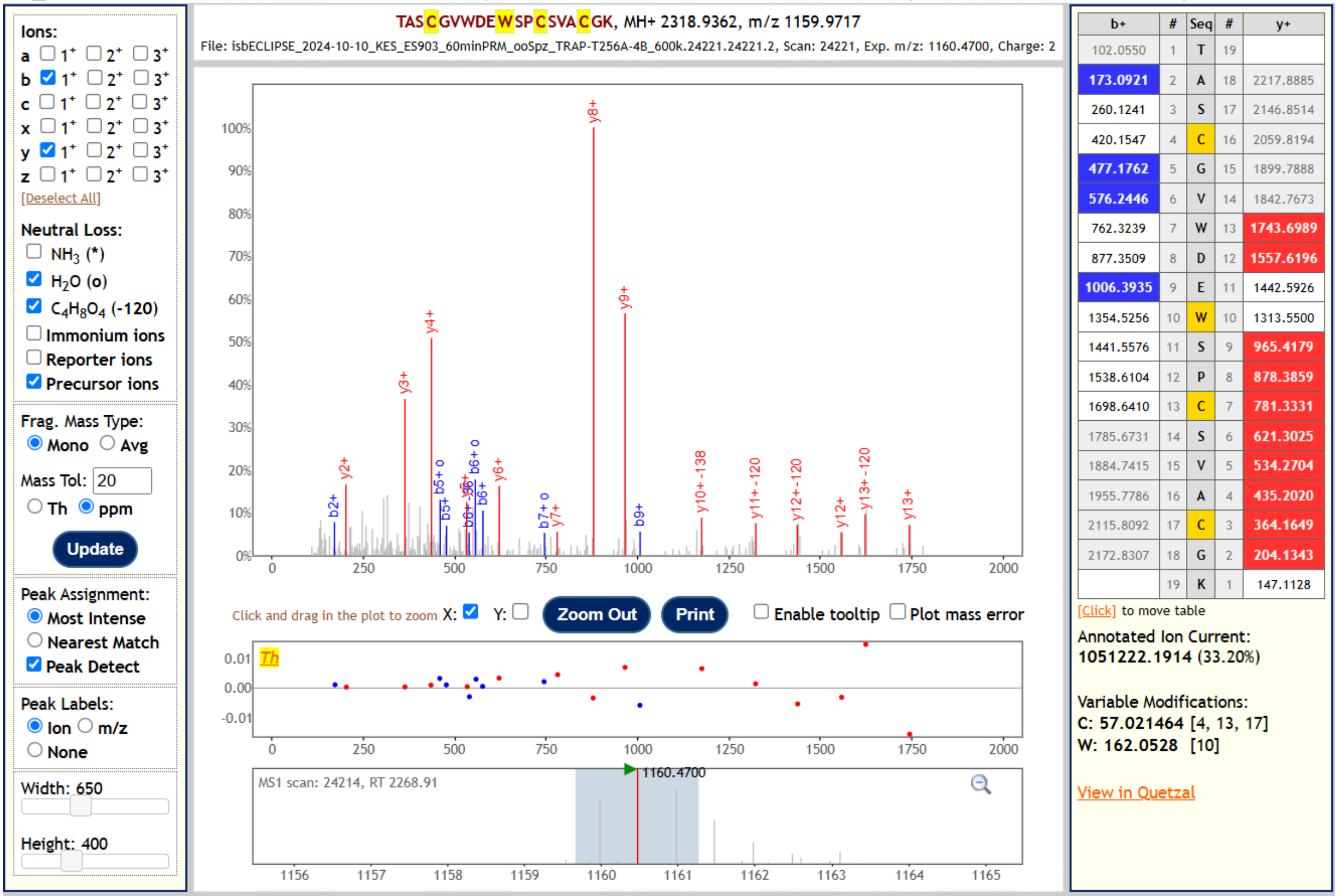


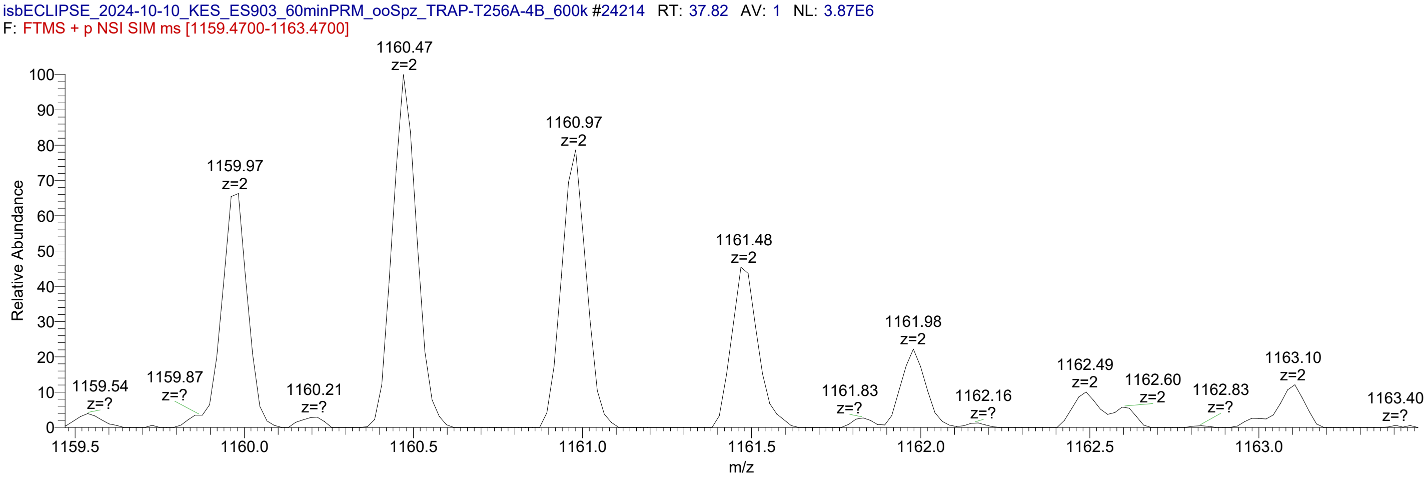

***Pf*****TRAP T256A clone 4B oocyst sporozoites TASCGVW[Man]DEW[Man]SPCSVACGK**

| xcorr score | Expect Score | Fragment ions | Retention time (min) | Retention time (sec) | MS2 window center m/z | MS2 window width | Target m/z | Observed Precursor m/z | Peptide m/z | Neutral Loss |
|---|---|---|---|---|---|---|---|---|---|---|
| 2.233 | 1.23E-08 | 10/36 | 33.61 | 2016.4 | 1241.50 | 1.6 | 1241.00 | 1241.00 | 1241.00 | 0.00 |

**USI:** mzspec:PXD064887:isbECLIPSE_2024-10-10_KES_ES903_60minPRM_ooSpz_TRAP-T256A-4B_600k:scan:21343:TASC[Carbamidomethyl]GVW[Hex]DEW[Hex]SPC[Carbamidomethyl]SVAC[Carbamidomethyl]GK/2

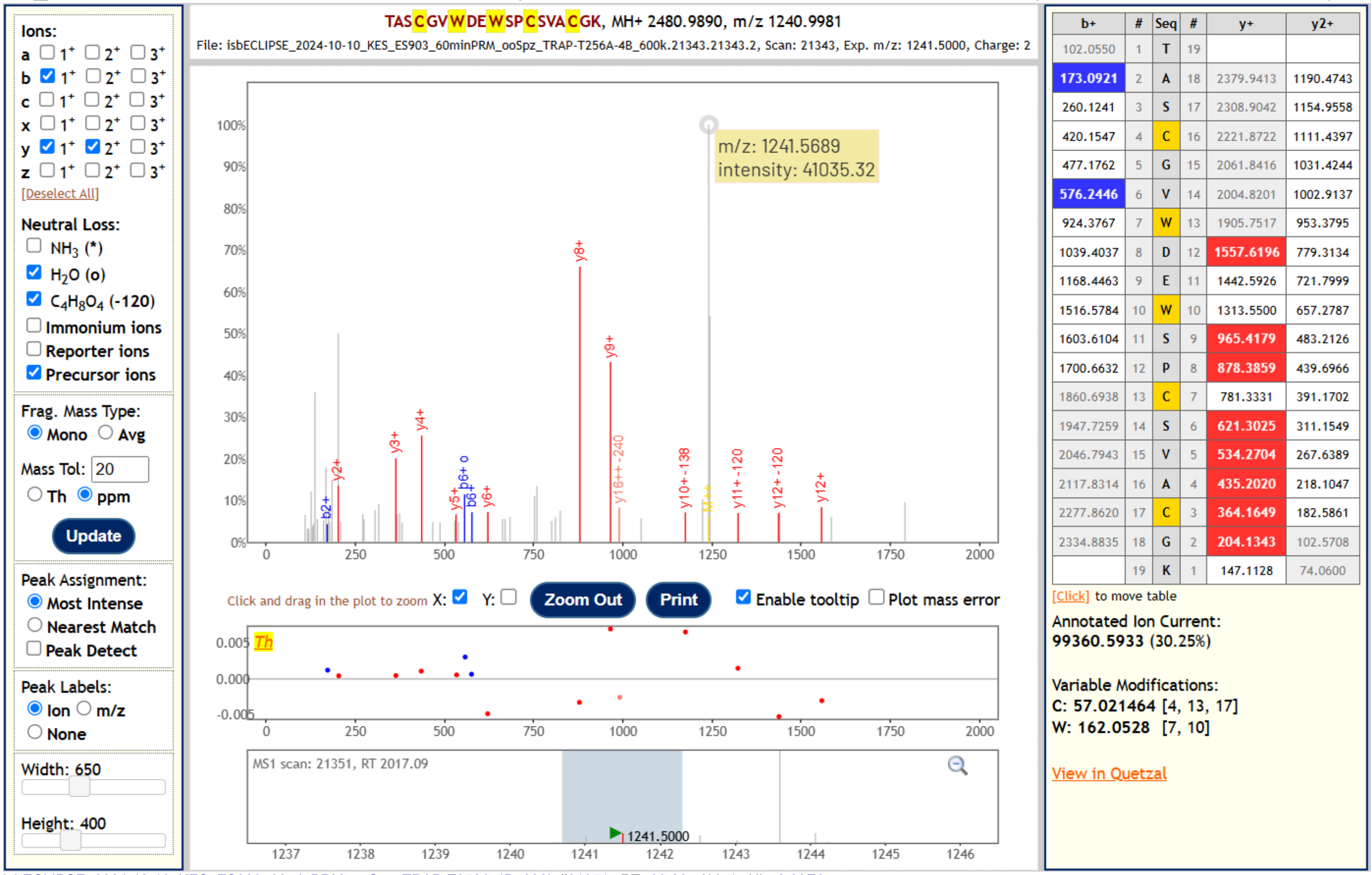


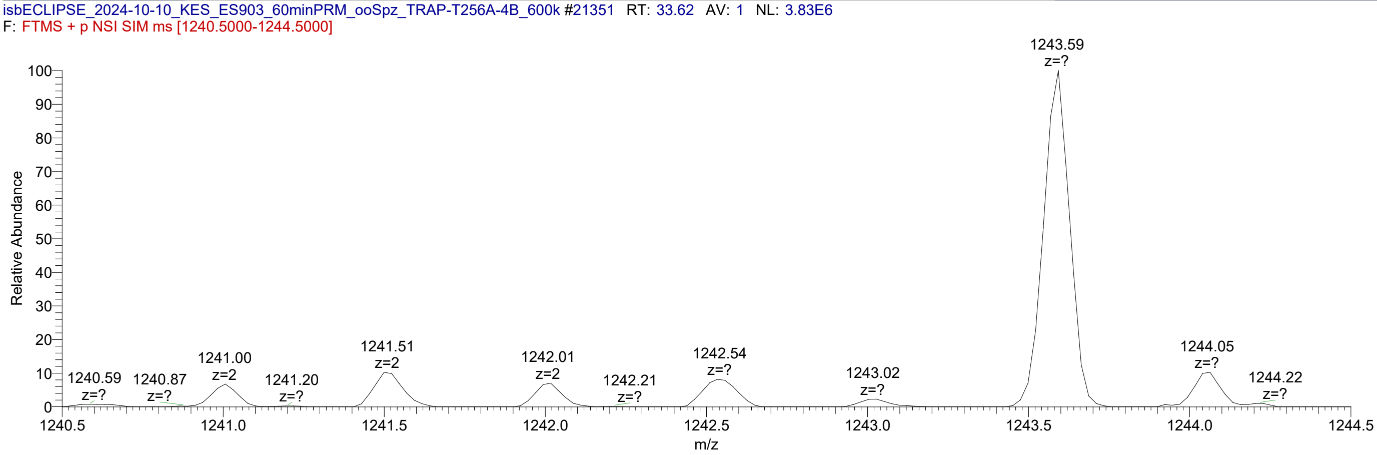


The unannotated MS2 peak 1241.5689 is a singly charged species of unknown origin, likely arising from an interfering species or background ion. No singly charged b- or y-ions positively identify the Trp-C-Man at Trp247 (the first Trp residue of the peptide). Comet lists the identity of this spectrum as TASCGVW[Ox]DEW[Man]SPCSVACGK with a neutral loss of 146 Da, i.e., O-Fuc. However, both explanations have equal Expect scores, and the isoform with oxidized Trp247 is simply listed first. The doubly mannosylated version of the peptide is the more likely explanation: not only does the fragment ion at 991.40 *m/z* match the doubly charged y16 ion with neutral loss of 120 from both Trp-C-Man moieties, but there is no O-Fuc attachment site, and there is no evidence that a moiety of 146 Da could be attached anywhere else.
