## Supplementary material for "Glycosylation of *Plasmodium falciparum* TRAP supports sporozoite motility and invasion": File S2

### FILE S2. Recombinant *Pf*TRAP TSR data.

#### Cell Line

CHO DPY19⁻ — A Chinese hamster ovary cell line lacking DPY19-L1, DPY19-L3, and DPY19-L4 (Shcherbakova *et al*. PMID: 28202721), adapted to grow in suspension in serum-free medium.

+*Pf*DPY19 — The cell line was transduced with *Pfdpy19.*

POFUT2⁻ — Endogenous *pofut2* was disrupted.

#### *Pf*TRAP TSR constructs

The *Pf*TRAP TSR constucts comprise residues 239-290 of *P. falciparum* TRAP (PlasmoDB.org gene ID PF3D7_1335900) as described in Wilder *et al*. (PMID: 3561879).

>PfTRAP_TSR_WT.HisAvi
EKTASCGVWDEWSPCSVTCGKGTRSRKREILHEGCTSELQEQCEEERCLPKRGSHHHHHHHHGLNDIFEAQKIEWHE
>PfTRAP_TSR_W250F.HisAvi
EKTASCGVWDE**F**SPCSVTCGKGTRSRKREILHEGCTSELQEQCEEERCLPKRGSHHHHHHHHGLNDIFEAQKIEWHE
>PfTRAP_TSR_T256A.HisAvi
EKTASCGVWDEWSPCSV**A**CGKGTRSRKREILHEGCTSELQEQCEEERCLPKRGSHHHHHHHHGLNDIFEAQKIEWHE

WT is the wild type sequence. The substitutions W250F and T256A refer to the localization of the glycosites $Trp^{250}$ and $Thr^{256}$ respectively, in the full-length *Pf*TRAP protein, and are the same substitutions made in the transgenic lines characterized in this manuscript. The His tag and Avi tag are represented in blue and orange text, respectively.

#### Guide sequences for the disruption of *C. griseus pofut2* gene

| Name | Sequence |
|---|---|
| Cg-pofut2-guide482 | GGGGGTTGACATCATACAGA |
| Cg-pofut2-guide483 | TGCTGGCAGCGGCATTCTGG |
| Cg-pofut2-guide484 | GGCGGCGGACATCCTGTCGG |

#### Diagnostic primers for screening the pofut2 gene disruptions.

| Name | Sequence |
|---|---|
| Cg-pofut2-oligo1 | GCGTTGCCCTGGATACTG |
| Cg-pofut2-oligo2 | GAGTTGTGAGAATTCATGGCGCGGACACACTTAC |
| Cg-pofut2-oligo3 | GCGCCATGAATTCTCACAACTCAAGGTCTGTGTC |
| Cg-pofut2-oligo4 | ACACCTGACGTCCACATCAG |

#### Source data for the Yield and $T_{melt}$ values given in Table 1 in the text.

Cell line and *Pf*TRAP TSR construct details are given above. Sample Number indicates independently produced batches of protein. Yield (mg), the total amount of protein recovered from the production run, was determined from the area under the curve of SEC traces (see below). Relative yield (µg/L culture) was determined from the Yield and the Culture Volume. The nominal yield based on SEC is reported for the O-fucosylation-null constructs (bottom two rows), but purity and recovery were too poor to accurately report yield or to obtain nanoDSF data. Sample purity (% Pure by MS) was determined by MS (see **File S3**). $T_{melt}$ was determined by nanoDSF (see below).

A total of three W250F samples were analyzed for this work. Yield data is reported here for W250F Samples 1 & 2, which were obtained from a cell line produced as described in Methods. Sample 3 was produced from a sub-clone that was obtained after selecting for higher yield in an attempt to improve recovery of the construct; consequently, yield from this line is not comparable to the other lines. To obtain this subclone, subclones from wells containing single colonies were seeded into 48-well plates and allowed to expand to confluency. Supernatants from confluent wells were clarified by centrifugation, diluted 1:1 in 10× KB buffer (1× PBS + 0.1% BSA, 0.05% NaN3, 0.02% Tween-20) and analyzed using biolayer interferometry (BLI) to identify wells with significantly higher production of the *Pf*TRAP_TSR constructs than that from the polyclonal parental lines. Measurements were performed using the Gator Plus instrument (GatorBio) with the HIS probes (GatorBio) for immobilization of the His-tagged *Pf*TRAP_TSR constructs with subsequent detection by 20 µg/mL mAb AKBR-5 (Wilder *et al.*, PMID: 3561879) diluted in 10× KB. Subclones that consistently produced relatively high levels of recombinant targets were further expanded, switched to suspension culture, and protein yields were assessed following target purification. MS confirmed that improved recovery from this subclone correlated with higher purity product. The $T_{melt}$ value was comparable to that obtained from Sample 1.

| Cell line | *Pf* TRAP TSR construct | Sample Number | Sample Date Produced | Culture Volume (L) | Yield (mg) | µg/L culture | Mean µg/L | S.D. | % Pure by MS | $T_{melt}$ by nanoDSF | | | |
|---|---|---|---|---|---|---|---|---|---|---|---|---|---|
| | | | | | | | | | | Rep 1 | Rep 2 | Mean | S.D |
| CHO DPY19$^{-}$ + *Pf* DPY19 | WT | Sample 1 | 2023-06-13 | 0.5 | 0.108 | 216 | 156 | 85 | 99.7% | 62.7 | 62.7 | 63.2 | 1.4 |
| CHO DPY19$^{-}$ + *Pf* DPY19 | WT | Sample 2 | 2023-11-15 | 2.0 | 0.192 | 96 | | | 99.0% | 62.1 | 65.2 | | |
| CHO DPY19$^{-}$ | WT | Sample 1 | 2023-08-15 | 1.0 | 0.054 | 54 | 46 | 12 | 99.8% | 55.8 | 55.4 | 55.3 | 1.1 |
| CHO DPY19$^{-}$ | WT | Sample 2 | 2023-08-30 | 1.0 | 0.037 | 37 | | | 99.2% | 53.8 | 56.2 | | |
| CHO DPY19$^{-}$ + *Pf* DPY19 | W250F | Sample 1 | 2023-06-13 | 0.5 | 0.016 | 32 | 20 | 17 | 80.7% | 42.0 | 41.7 | 41.1 | 1.3 |
| CHO DPY19$^{-}$ + *Pf* DPY19 | W250F | Sample 2 | 2024-01-04 | 3.0 | 0.023 | 7.7 | | | | | | | |
| CHO DPY19$^{-}$ + *Pf* DPY19 | W250F | Sample 3 | 2024-02-28 | 0.5 | 0.071 | 142 | | | 99.8% | 39.2 | 41.4 | | |
| CHO DPY19$^{-}$, POFUT2$^{-}$, +*Pf* DPY19 | WT | Sample 1 | 2024-07-02 | 1.0 | 0.024 | 24 | | | 21.4% | | | | |
| CHO DPY19$^{-}$ + *Pf* DPY19 | T256A | Sample 1 | 2024-10-15 | 3.0 | 0.016 | 5.3 | | | 53.0% | | | | |

#### Differential Scanning Fluorimetry (DSF)

Source data for the $T_{melt}$ values given in Table 1 in the text. Fluorescence (ratio of 350 nm and 330 nm) with respect to temperature is plotted for each protein construct. Two technical replicates (left two columns) of two independently produced samples are shown. A spline is fitted to each curve (blue line). The inflection point, *i.e.*, $T_{melt}$, is shown by a dashed line. This inflection point was determined from the maximum of the first derivative plot (right).

**CHO DPY19⁻ +*Pf*DPY19** **_Pf_TRAP TSR WT**

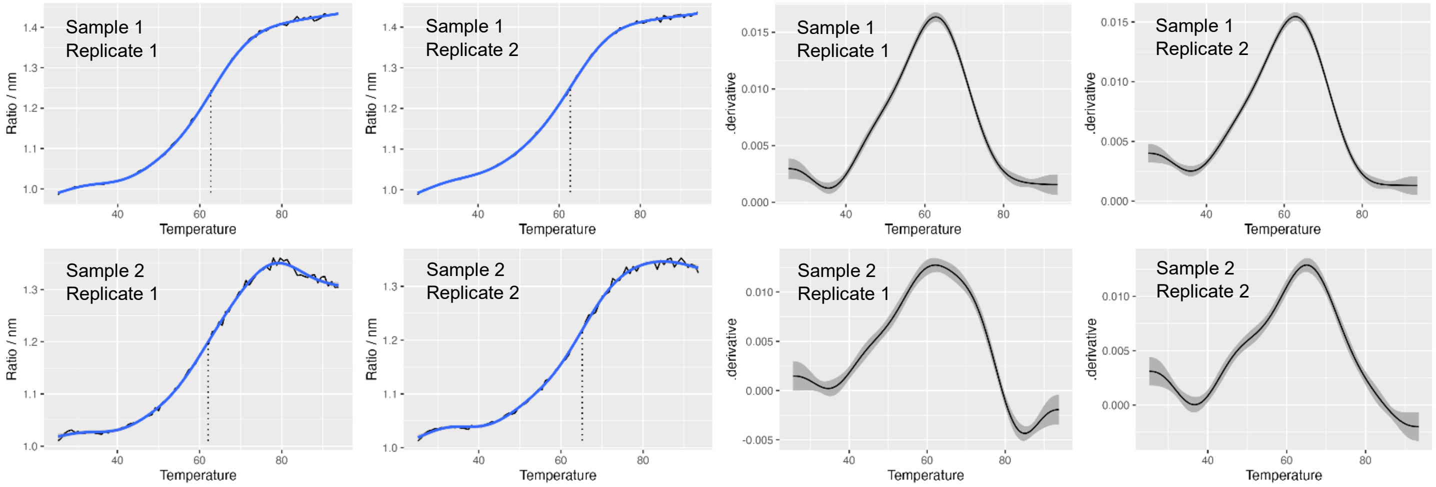


**CHO DPY19⁻** **_Pf_TRAP TSR WT**

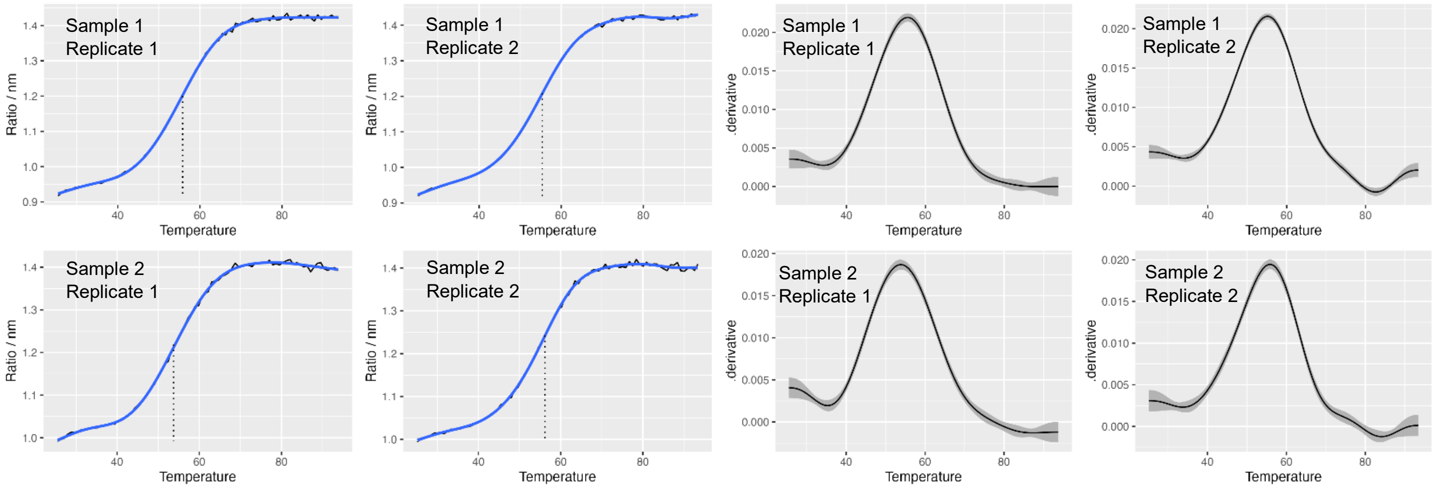


**CHO DPY19⁻ +*Pf*DPY19** **PfTRAP TSR W250F**

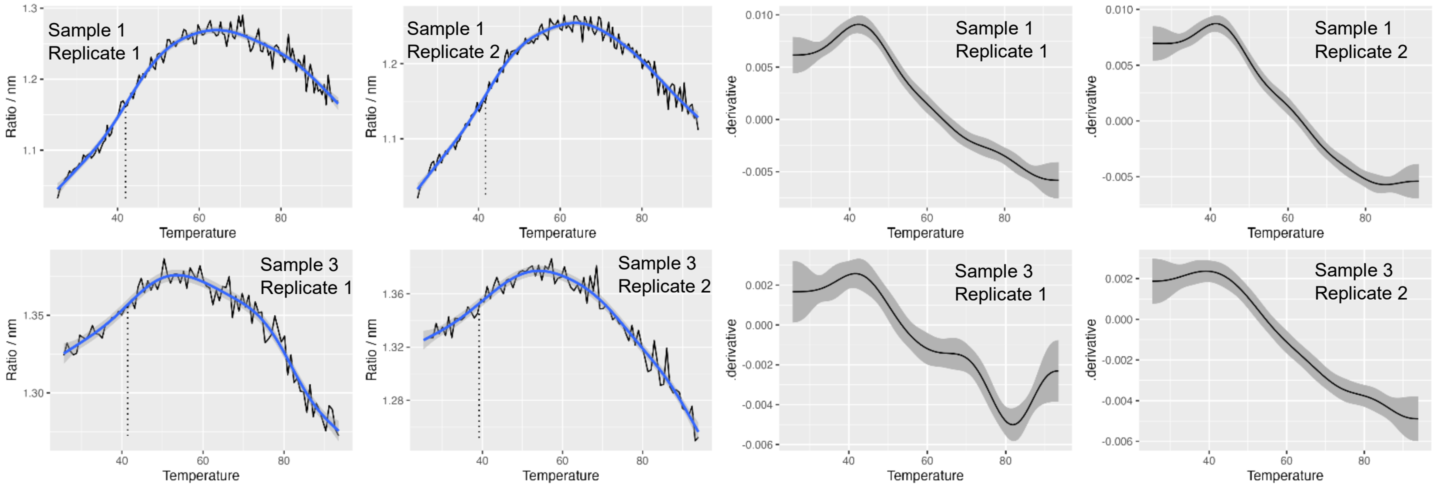

#### Protein yield determined by Size Exclusion Chromatography (SEC)

Source data for the Yield values given in Table 1 in the text.

| Cell line | *Pf* TRAP TSR construct | Sample Number | Sample Date Produced | Culture Volume (L) | Yield (mg) | µg/L culture | % Pure by MS | $T_{melt}$ by nanoDSF | |
|---|---|---|---|---|---|---|---|---|---|
| | | | | | | | | Rep 1 | Rep 2 |
| CHO DPY19$^{-}$ + *Pf* DPY19 | WT | Sample 1 | 2023-06-13 | 0.5 | 0.108 | 216 | 99.7% | 62.7 | 62.7 |

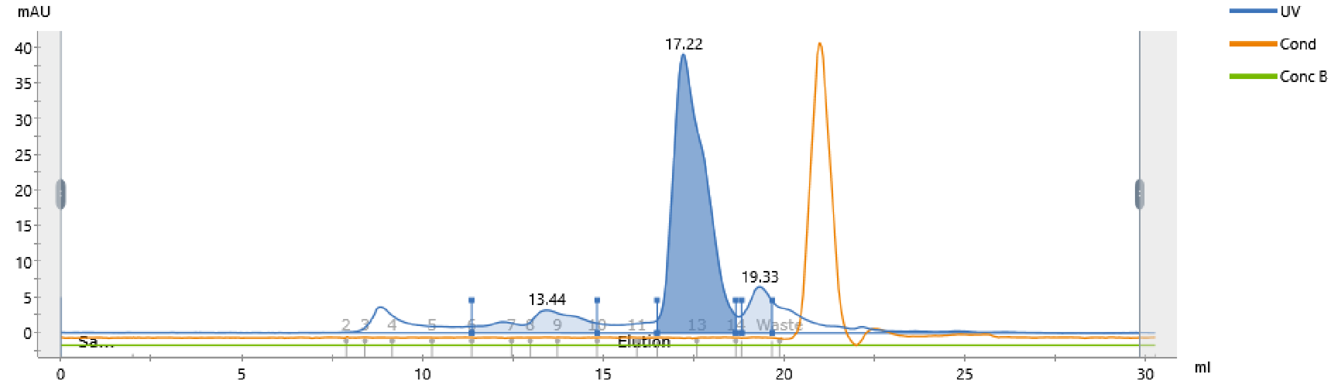


| Peak | Retention ml | Area ml*mAU | Area % | | Ext coeff. mg $^{-1}$ ml cm $^{-1}$ | Fraction(s) | Volume ml | Amount mg | Concentration mg/ml | Conductivity mS/cm |
|---|---|---|---|---|---|---|---|---|---|---|
| Peak A | 13.443 | 6.201 | | 12.23 | | 6 - 9 | 3.469 | | | 17.20 |
| Peak B | 17.216 | 40.33 | | 79.57 | 1.873 | 12 - 13 | 2.177 | 0.108 | 0.049 | 17.20 |
| Peak C | 19.325 | 4.153 | | 8.19 | | 15 | 0.844 | | | 17.20 |

| Cell line | *Pf* TRAP TSR construct | Sample Number | Sample Date Produced | Culture Volume (L) | Yield (mg) | µg/L culture | % Pure by MS | $T_{melt}$ by nanoDSF | |
|---|---|---|---|---|---|---|---|---|---|
| | | | | | | | | Rep 1 | Rep 2 |
| CHO DPY19$^{-}$ + *Pf* DPY19 | WT | Sample 2 | 2023-11-15 | 2.0 | 0.192 | 96 | 99.0% | 62.1 | 65.2 |

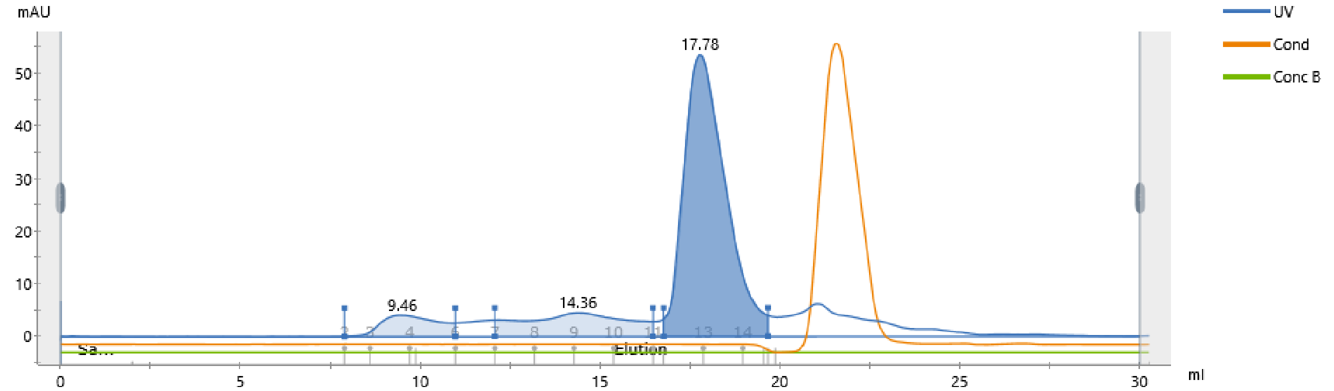


| Peak | Retention ml | Area ml*mAU | Area % | | Ext coeff. mg $^{-1}$ ml cm $^{-1}$ | Fraction(s) | Volume ml | Amount mg | Concentration mg/ml | Conductivity mS/cm |
|---|---|---|---|---|---|---|---|---|---|---|
| Peak A | 9.457 | 7.492 | | 7.92 | | 2 - 5 | 3.083 | | | 17.25 |
| Peak B | 14.358 | 15.02 | | 15.88 | | 7 - 10 | 4.400 | | | 17.25 |
| Peak C | 17.779 | 72.05 | | 76.19 | 1.873 | 12 - 15 | 2.901 | 0.192 | 0.066 | 17.22 |

| Cell line | *Pf* TRAP TSR construct | Sample Number | Sample Date Produced | Culture Volume (L) | Yield (mg) | µg/L culture | % Pure by MS | $T_{melt}$ by nanoDSF | |
|---|---|---|---|---|---|---|---|---|---|
| | | | | | | | | Rep 1 | Rep 2 |
| CHO DPY19$^{-}$ | WT | Sample 1 | 2023-08-15 | 1.0 | 0.054 | 54 | 99.8% | 55.8 | 55.4 |

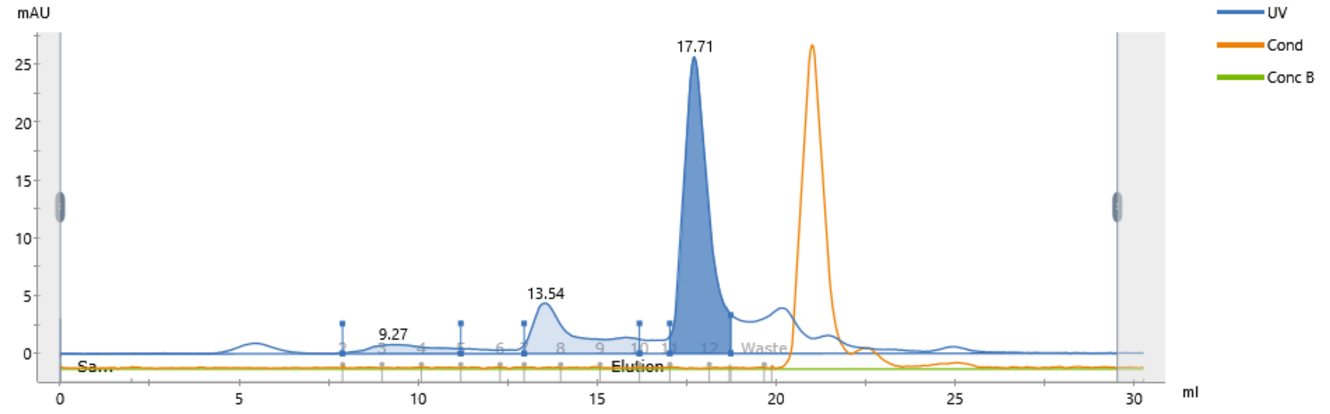


| Peak | Retention ml | Area ml*mAU | Area % | Ext coeff. mg $^{-1}$ ml cm $^{-1}$ | Fraction(s) | Volume ml | Amount mg | Concentration mg/ml | Conductivity mS/cm |
|---|---|---|---|---|---|---|---|---|---|
| Peak A | 9.272 | 1.626 | 5.82 | | 2 - 4 | 3.300 | | | 17.22 |
| Peak B | 13.542 | 6.155 | 22.04 | | 7 - 9 | 3.217 | | | 17.22 |
| Peak C | 17.712 | 20.15 | 72.14 | 1.873 | 11 - 13 | 1.700 | 0.054 | 0.032 | 17.21 |

| Cell line | *Pf* TRAP TSR construct | Sample Number | Sample Date Produced | Culture Volume (L) | Yield (mg) | µg/L culture | % Pure by MS | $T_{melt}$ by nanoDSF | |
|---|---|---|---|---|---|---|---|---|---|
| | | | | | | | | Rep 1 | Rep 2 |
| CHO DPY19$^{-}$ | WT | Sample 2 | 2023-08-30 | 1.0 | 0.037 | 37 | 99.2% | 53.8 | 56.2 |

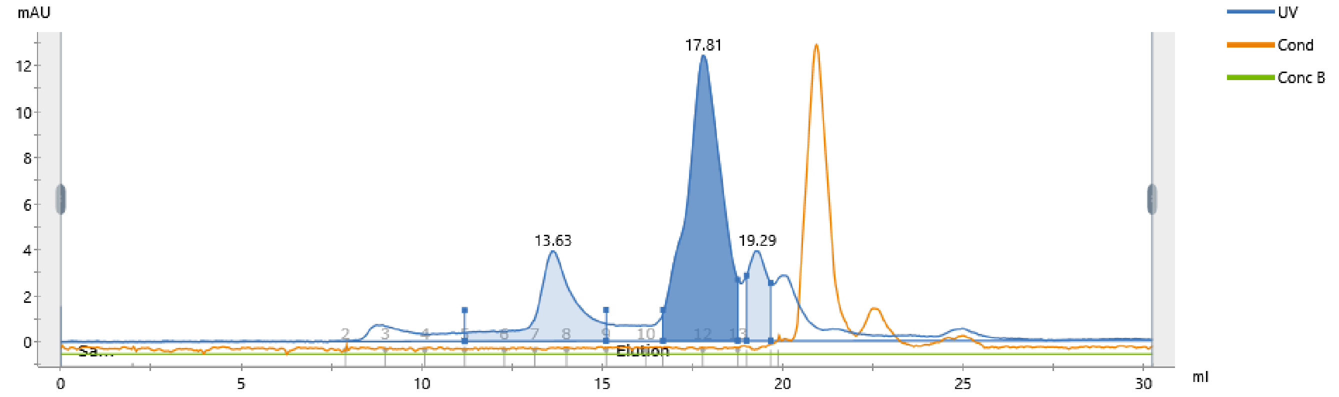


| Peak | Retention ml | Area ml*mAU | Area % | Ext coeff. mg $^{-1}$ ml cm $^{-1}$ | Fraction(s) | Volume ml | Amount mg | Concentration mg/ml | Conductivity mS/cm |
|---|---|---|---|---|---|---|---|---|---|
| Peak A | 13.634 | 4.852 | 23.32 | | 5 - 8 | 3.921 | | | 17.23 |
| Peak B | 17.810 | 13.72 | 65.94 | 1.873 | 11 - 12 | 2.071 | 0.037 | 0.018 | 17.23 |
| Peak C | 19.289 | 2.234 | 10.74 | | 14 | 0.667 | | | 17.23 |

| Cell line | *Pf* TRAP TSR construct | Sample Number | Sample Date Produced | Culture Volume (L) | Yield (mg) | µg/L culture | % Pure by MS | $T_{melt}$ by nanoDSF | |
|---|---|---|---|---|---|---|---|---|---|
| | | | | | | | | Rep 1 | Rep 2 |
| CHO DPY19$^{-}$ + *Pf* DPY19 | W250F | Sample 1 | 2023-06-13 | 0.5 | 0.016 | 32 | 80.7% | 42.0 | 41.7 |

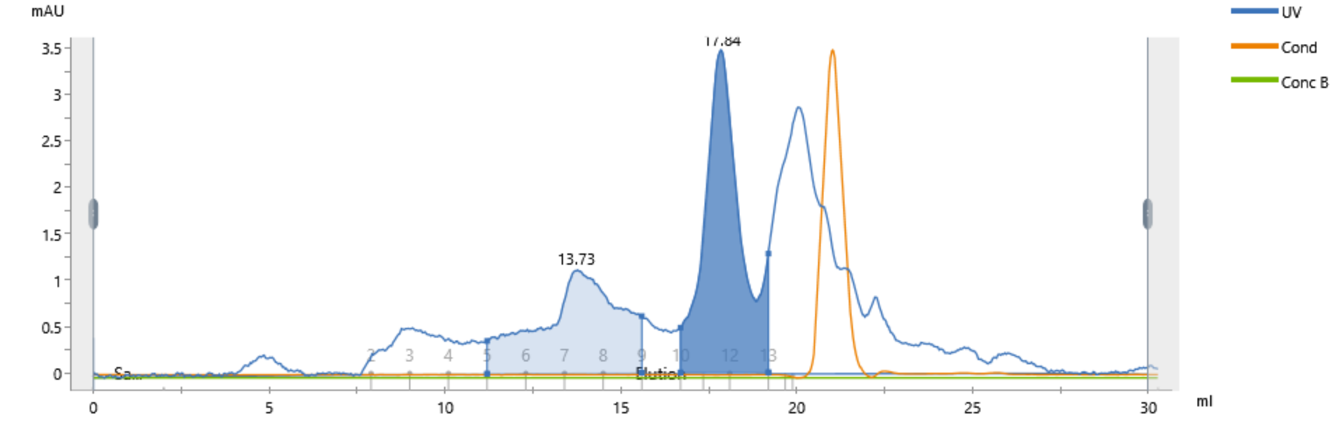


| Peak | Retention ml | Area ml*mAU | Area % | Ext coeff. mg $^{-1}$ ml cm $^{-1}$ | Fraction(s) | Volume ml | Amount mg | Concentration mg/ml | Conductivity mS/cm |
|---|---|---|---|---|---|---|---|---|---|
| Peak A | 13.731 | 2.893 | 41.63 | | 5 - 8 | 4.400 | | | 17.20 |
| Peak B | 17.838 | 4.057 | 58.37 | 1.268 | 10 - 12 | 2.496 | 0.016 | 0.006 | 17.20 |

| Cell line | *Pf* TRAP TSR construct | Sample Number | Sample Date Produced | Culture Volume (L) | Yield (mg) | µg/L culture | % Pure by MS | $T_{melt}$ by nanoDSF | |
|---|---|---|---|---|---|---|---|---|---|
| | | | | | | | | Rep 1 | Rep 2 |
| CHO DPY19$^{-}$ + *Pf* DPY19 | W250F | Sample 2 | 2024-01-04 | 3.0 | 0.023 | 7.7 | | | |

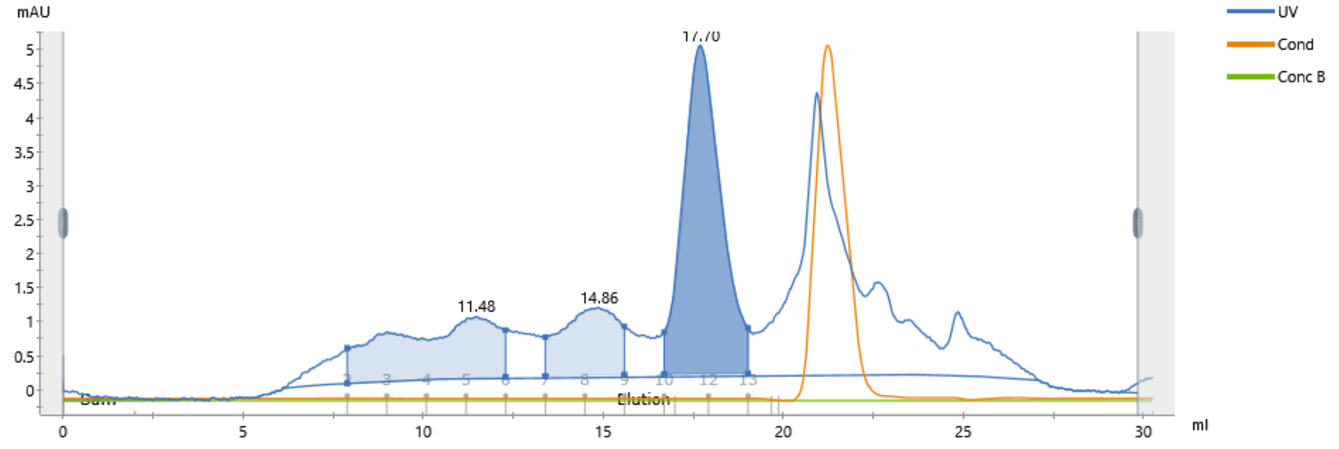


| Peak | Retention ml | Area ml*mAU | Area % | Ext coeff. mg $^{-1}$ ml cm $^{-1}$ | Fraction(s) | Volume ml | Amount mg | Concentration mg/ml | Conductivity mS/cm |
|---|---|---|---|---|---|---|---|---|---|
| Peak A | 11.475 | 3.012 | 28.18 | | 2 - 5 | 4.400 | | | 17.22 |
| Peak B | 14.855 | 1.798 | 16.81 | | 7 - 8 | 2.199 | | | 17.22 |
| Peak C | 17.704 | 5.881 | 55.01 | 1.268 | 10 - 12 | 2.330 | 0.023 | 0.010 | 17.22 |

| Cell line | *Pf* TRAP TSR construct | Sample Number | Sample Date Produced | Culture Volume (L) | Yield (mg) | µg/L culture | % Pure by MS | $T_{melt}$ by nanoDSF Rep 1 | $T_{melt}$ by nanoDSF Rep 2 |
|---|---|---|---|---|---|---|---|---|---|
| CHO DPY19⁻ + *Pf* DPY19 | W250F | Sample 3 | 2024-02-28 | 0.5 | 0.071 | 142 | 99.8% | 39.2 | 41.4 |

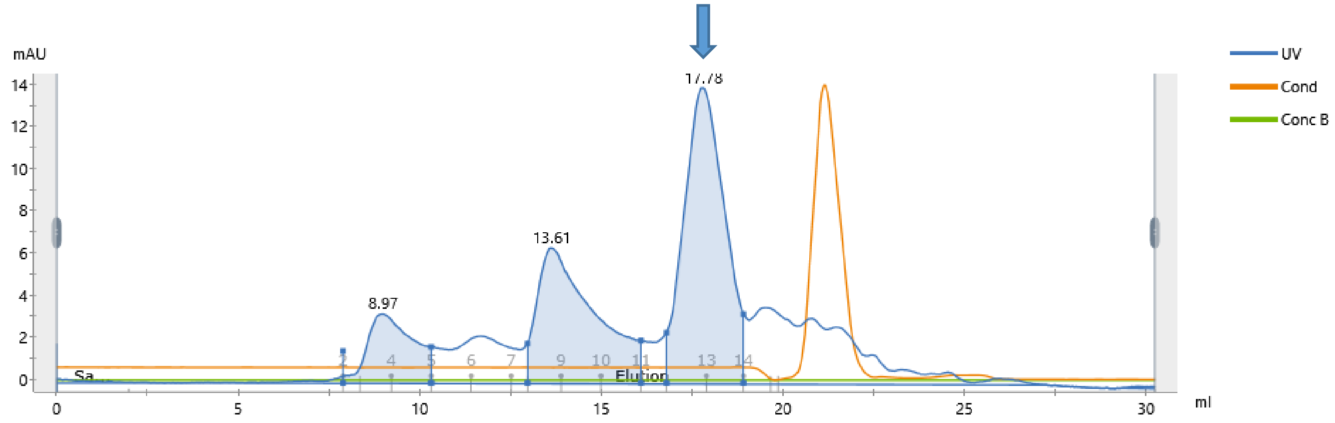


| Peak | Retention ml | Area ml*mAU | Area % | Ext coeff. mg⁻¹ ml cm⁻¹ | Fraction(s) | Volume ml | Amount mg | Concentration mg/ml | Conductivity mS/cm |
|---|---|---|---|---|---|---|---|---|---|
| Peak A | 8.973 | 4.753 | 13.82 | | 2 - 4 | 2.430 | | | 17.81 |
| Peak B | 13.612 | 11.70 | 34.02 | | 8 - 10 | 3.122 | | | 17.81 |
| Peak C | 17.781 | 17.95 | 52.17 | 1.268 | 12 - 13 | 2.118 | 0.071 | 0.033 | 17.81 |

| Cell line | *Pf* TRAP TSR construct | Sample Number | Sample Date Produced | Culture Volume (L) | Yield (mg) | µg/L culture | % Pure by MS | $T_{melt}$ by nanoDSF Rep 1 | $T_{melt}$ by nanoDSF Rep 2 |
|---|---|---|---|---|---|---|---|---|---|
| CHO DPY19⁻, POFUT2⁻, +*Pf* DPY19 | WT | Sample 1 | 2024-07-02 | 1.0 | 0.024 | 24 | 21.4% | | |

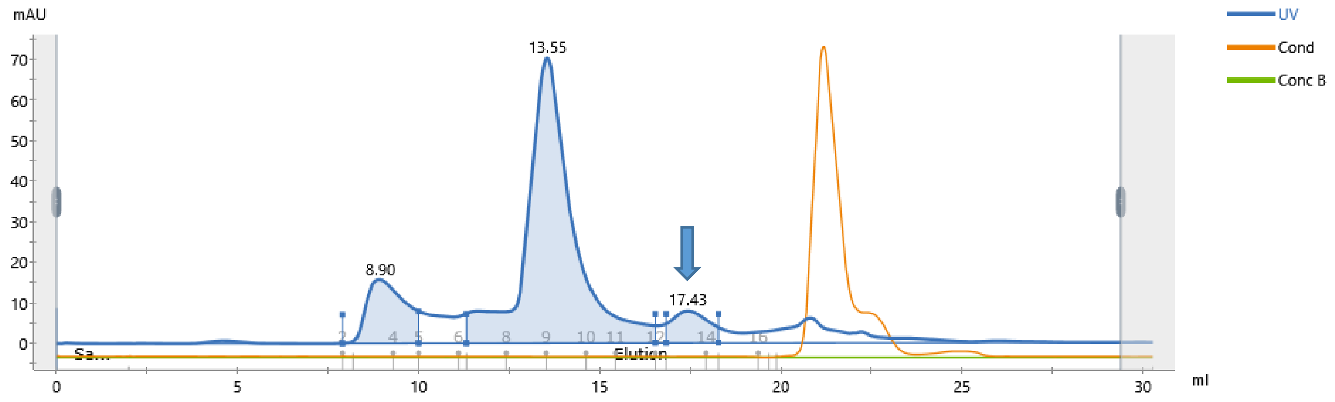


| Peak | Retention ml | Area ml*mAU | Area % | Ext coeff. mg⁻¹ ml cm⁻¹ | Fraction(s) | Volume ml | Amount mg | Concentration mg/ml | Conductivity mS/cm |
|---|---|---|---|---|---|---|---|---|---|
| Peak A | 8.905 | 19.03 | 17.55 | | 2 - 4 | 2.105 | | | 16.07 |
| Peak B | 13.548 | 80.54 | 74.29 | | 8 - 9 | 2.201 | | | 16.06 |
| Peak C | 17.433 | 8.845 | 8.16 | 1.873 | 13 - 14 | 1.440 | 0.024 | 0.016 | 16.06 |

| Cell line | *Pf* TRAP TSR construct | Sample Number | Sample Date Produced | Culture Volume (L) | Yield (mg) | µg/L culture | % Pure by MS | $T_{melt}$ by nanoDSF Rep 1 | $T_{melt}$ by nanoDSF Rep 2 |
|---|---|---|---|---|---|---|---|---|---|
| CHO DPY19⁻ + *Pf* DPY19 | T256A | Sample 1 | 2024-10-15 | 3.0 | 0.016 | 5.3 | 53.0% | | |

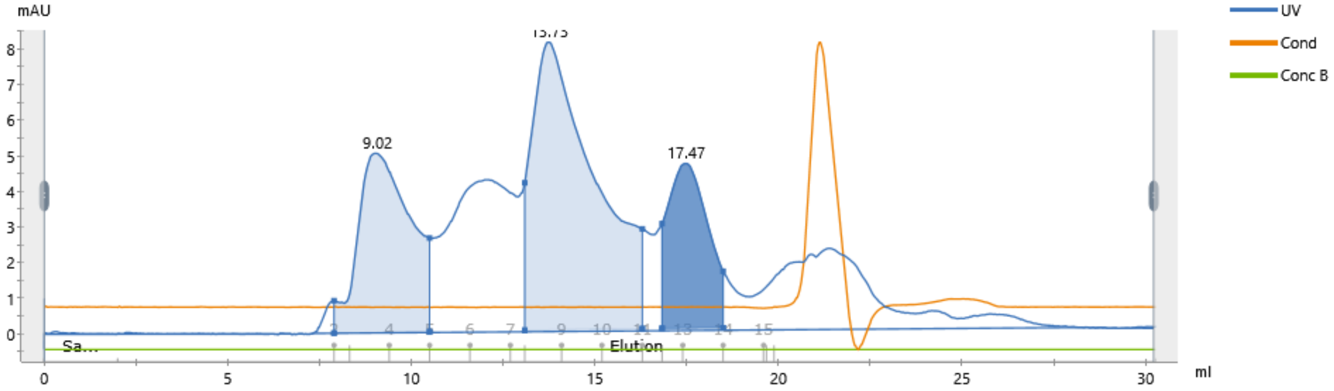


| Peak | Retention ml | Area ml*mAU | Area % | Ext coeff. mg⁻¹ ml cm⁻¹ | Fraction(s) | Volume ml | Amount mg | Concentration mg/ml | Conductivity mS/cm |
|---|---|---|---|---|---|---|---|---|---|
| Peak A | 9.024 | 8.389 | 27.69 | | 2 - 4 | 2.604 | | | 23.37 |
| Peak B | 13.734 | 16.07 | 53.05 | | 8 - 10 | 3.200 | | | 23.37 |
| Peak C | 17.473 | 5.837 | 19.27 | 1.879 | 12 - 13 | 1.663 | 0.016 | 0.009 | 23.38 |
