## Supplementary material for "Glycosylation of *Plasmodium falciparum* TRAP supports sporozoite motility and invasion": File S3

### FILE S3. Extracted ion chromatograms and mass spectra of the *Pf*TRAP glycopeptide detected in recombinant *Pf*TRAP TSR.

All source data for the figures shown here, including raw mass spectrometry data, extracted ion chromatogram values, and complete peak lists of identified peptides, have been deposited to the ProteomeXchange Consortium via the MassIVE partner repository with the dataset identifier PXD064887.
Files can be downloaded from https://massive.ucsd.edu/ProteoSAFe/static/massive.jsp

All MS2 spectra used to identify the *Pf*TRAP TSR glycopeptide, including those shown here, have been assigned Universal Spectrum Identifiers (USI) to enable investigation of the peak annotations. USI viewers are available at

- https://proteomecentral.proteomexchange.org/usi/
- https://proteomecentral.proteomexchange.org/quetzal/
- https://massive.ucsd.edu/ProteoSAFe/usi.jsp

Note that only Quetzal annotates peaks with neutral loss of 120.04 Da arising from cross-ring cleavage of Trp-C-Man.

Mass spectrometry evidence for glycosylation of recombinant *Pf*TRAP TSR is provided here for eight different samples. A total of three W250F samples were analyzed for this work. Yield data were taken for Samples 1 & 2. Sample 3 was produced from a sub-clone that was obtained after selecting for higher yield in an attempt to improve recovery of the construct. DSF and MS data are reported here for Samples 1 & 3.

| Cell line | *Pf* TRAP TSR construct | Sample Number | µg/L culture | Mean µg/L | S.D. | % Pure by MS | $T_{melt}$ by nanoDSF | | | |
|---|---|---|---|---|---|---|---|---|---|---|
| | | | | | | | Rep 1 | Rep 2 | Mean | S.D |
| CHO DPY19$^{-}$ + *Pf* DPY19 | WT | Sample 1 | 216 | 156 | 85 | 99.7% | 62.7 | 62.7 | 63.2 | 1.4 |
| CHO DPY19$^{-}$ + *Pf* DPY19 | WT | Sample 2 | 96 | | | 99.0% | 62.1 | 65.2 | | |
| CHO DPY19$^{-}$ | WT | Sample 1 | 54 | 46 | 12 | 99.8% | 55.8 | 55.4 | 55.3 | 1.1 |
| CHO DPY19$^{-}$ | WT | Sample 2 | 37 | | | 99.2% | 53.8 | 56.2 | | |
| CHO DPY19$^{-}$ + *Pf* DPY19 | W250F | Sample 1 | 32 | 20 | 17 | 80.7% | 42.0 | 41.7 | 41.1 | 1.3 |
| CHO DPY19$^{-}$ + *Pf* DPY19 | W250F | Sample 2 | 7.7 | | | | | | | |
| CHO DPY19$^{-}$ + *Pf* DPY19 | W250F | Sample 3 | 142 | | | 99.8% | 39.2 | 41.4 | | |
| CHO DPY19$^{-}$, POFUT2$^{-}$, +*Pf* DPY19 | WT | Sample 1 | 24 | | | 21.4% | | | | |
| CHO DPY19$^{-}$ + *Pf* DPY19 | T256A | Sample 1 | 5.3 | | | 53.0% | | | | |

**Cell Line**

CHO DPY19$^{-}$ A Chinese hamster ovary cell line lacking DPY19-L1, L3, and L4 (Shcherbakova *et al*. PMID: 28202721), adapted to grow in suspension in serum-free medium.

+*Pf*DPY19 The cell line was transduced with *Pfdpy19.*

POFUT2$^{-}$ Endogenous *POFUT2* was disrupted.

**The full sequences of the *Pf*TRAP TSR constructs expressed were as follows:**

>PfTRAP_TSR_WT.HisAvi
EKTASCGVWDEWSPCSVTCGKGTRSRKREILHEGCTSELQEQCEEERCLPKRGSHHHHHHHHGLNDIFEAQKIEWHE
>PfTRAP_TSR_W250F.HisAvi
EKTASCGVWDEFSPCSVTCGKGTRSRKREILHEGCTSELQEQCEEERCLPKRGSHHHHHHHHGLNDIFEAQKIEWHE
>PfTRAP_TSR_T256A.HisAvi
EKTASCGVWDEWSPCSVACGKGTRSRKREILHEGCTSELQEQCEEERCLPKRGSHHHHHHHHGLNDIFEAQKIEWHE

WT is the wild type sequence. The substitutions W250F and T256A refer to the localization of the glycosites Trp$^{250}$ and Thr$^{256}$ respectively, in the full-length *Pf*TRAP protein, and are the same substitutions made in the transgenic lines characterized in this manuscript. The His tag and Avi tag are represented in blue and orange text, respectively.

**% Pure by MS.** The proportion of signal originating from *Pf*TRAP relative to background CHO protein was calculated by summing the maximum precursor intensities (extracted from the mzML by InteractParser in the TPP) for all detected peptide ions with unique *m/z*. Detailed peak lists and underlying calculations are available with the raw data in MassIVE.

| Cell line | *Pf* TRAP TSR construct | Sample Number | Sample Date Produced | Stoichiometry of major glycoforms from MS data | | | | | | | | | | | |
|---|---|---|---|---|---|---|---|---|---|---|---|---|---|---|---|
| | | | | C-Man | Mean | S.D | C-Man×2 | Mean | S.D | O-Fuc | Mean | S.D | O-Fuc-Glc | Mean | S.D |
| CHO DPY19$^{-}$ + *Pf* DPY19 | WT | Sample 1 | 2023-06-13 | 20.92% | 22.05% | 1.61% | 0.00% | 0.00% | 0.00% | 0.29% | 0.26% | 0.05% | 99.71% | 99.74% | 0.05% |
| CHO DPY19$^{-}$ + *Pf* DPY19 | WT | Sample 2 | 2023-11-15 | 23.19% | | | 0.00% | | | 0.22% | | | 99.78% | | |
| CHO DPY19$^{-}$ | WT | Sample 1 | 2023-08-15 | 0.00% | 0.00% | 0.00% | 0.00% | 0.00% | 0.00% | 0.23% | 0.29% | 0.09% | 99.77% | 99.71% | 0.09% |
| CHO DPY19$^{-}$ | WT | Sample 2 | 2023-08-30 | 0.00% | | | 0.00% | | | 0.36% | | | 99.64% | | |
| CHO DPY19$^{-}$ + *Pf* DPY19 | W250F | Sample 1 | 2023-06-13 | 0.00% | | | 0.00% | | | 0.61% | | | 98.68% | | |
| CHO DPY19$^{-}$ + *Pf* DPY19 | W250F | Sample 2 | 2024-01-04 | | 0.00% | 0.00% | | 0.00% | 0.00% | | 0.77% | 0.21% | | 97.86% | 1.17% |
| CHO DPY19$^{-}$ + *Pf* DPY19 | W250F | Sample 3 | 2024-02-28 | 0.00% | | | 0.00% | | | 0.92% | | | 97.03% | | |
| CHO DPY19$^{-}$, POFUT2$^{-}$, +*Pf* DPY19 | WT | Sample 1 | 2024-07-02 | 24.96% | | | 0.34% | | | 0.02% | | | 1.36% | | |
| CHO DPY19$^{-}$ + *Pf* DPY19 | T256A | Sample 1 | 2024-10-15 | 13.86% | | | 0.35% | | | 0.00% | | | 0.00% | | |

**Stoichiometry**

O-Fuc O-fucose at Thr$^{256}$

O-Fuc-Glc O-fucose-β-1,3-glucose at Thr$^{256}$

C-Man C-mannose at Trp$^{250}$

C-Man×2 C-mannose at Trp$^{250}$ and Trp$^{247}$

The following types of evidence are provided for each sample:

1. **Extracted ion chromatogram.** Xcalibur Qual Browser was used to extract the MS1 signal with respect to retention time from a narrow *m/z* range (0.1–0.2 *m/z*) containing the monoisotopic peak of the predicted glycoforms in each experiment. Peaks were only plotted if an MS2 spectrum at or near the apex produced a high-quality peptide spectrum match confirming the identity of the glycoform.
2. **Table of detected glycoforms and relative abundance.** Each data-dependent experiment method included an inclusion list targeting seven masses corresponding to the 12 possible glycoforms of the *Pf*TRAP TSR glycopeptide that contain the three predicted glycosites. These glycoforms encompass all possible permutations of C-Man at Trp$^{247}$ and Trp$^{250}$ and O-Fuc or O-Fuc-Glc at Thr$^{256}$. The table lists the glycoforms detected in the sample along with the retention time and MS1 peak height observed in the extracted ion chromatogram. These abundances are used to calculate the relative abundance of each glycoform reported in Table 1 of the manuscript. Not included in the tables are variants of the peptide detected at low abundance (<1% of the dominant species), including the peptide with oxidized Trp and various non-canonical glycoforms that are likely artifacts of the recombinant expression system, e.g., O-linked Hex and O-linked dHex from protein O-fucosyltransferases other than POFUT2. The full list of peptide spectrum matches identifying all glycoforms of the peptide in each experiment is available along with the raw data in MassIVE.

The following additional data is shown for representative glycopeptides detected in WT, W250F, and T256A variants of the *Pf*TRAP TSR:

1. **Representative MS2 spectra** annotated by the Lorikeet viewer as implemented in the Trans-Proteomic Pipeline were selected for each of the glycoforms shown in the XIC. Neutral loss of 120.04 Da ($C_4H_8O_4$) from cross-ring cleavage of Trp-C-Man is annotated when present.
2. **Spectrum information**, including the xcorr and Expect scores from the Comet search, the selected precursor *m/z*, and the *m/z* of the detected peptide. Note that, since O-linked glycans are labile in the gas phase, evidence of O-Fuc and O-Fuc-Glc present as neutral loss of 146.06 and 308.11 Da, respectively.
3. **The Universal Spectrum Identifier (USI)** for the representative spectrum. This USI can be used to investigate the annotated MS2 spectrum in the USI viewers listed above. These viewers allow the user to investigate alternative peptide sequences, modifications, and modification localizations. Note that the peptide sequence may include {dHex} or {dHex(1)Hex(1)} to account for the unlocalized O-glycan.

**1. *Pf*TRAP TSR WT expressed in CHO DPY19⁻ transduced with *Pfdpy19***
**Glycopeptide sequence: TASCGVWDEWSPCSVTCGK**

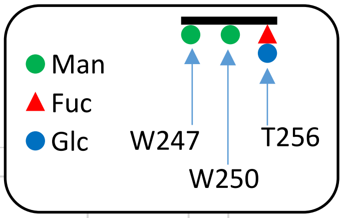


Sample 1 (2023-06-13)    Sample 2 (2023-11-15)

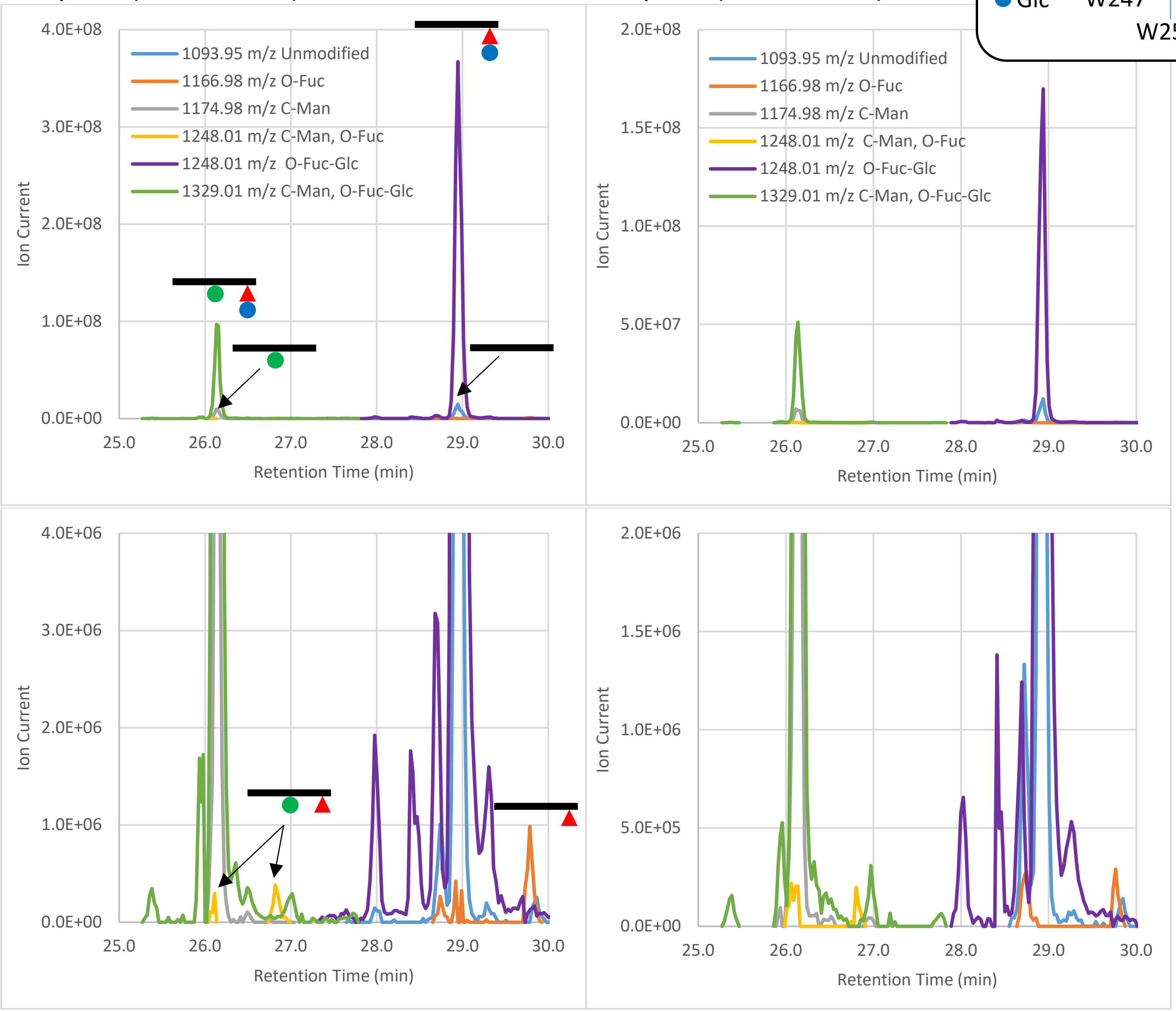


**Fig 1. Extracted ion chromatograms of the detected glycopeptides.** Left is Sample 1, right is Sample 2. Bottom plots are zoomed in on the y-axis to show low-abundance species. The unmodified peptide (1093.95 *m/z*) and the O-Fuc peptide (1166.95 *m/z*) co-eluting with the dominant O-Fuc-Glc peptide (1248.01 *m/z*) at 28.9 min arise from neutral loss of the O-glycan due to in-source fragmentation that occurs after the peptide has eluted from the column. The same phenomenon is observed for the secondary peptide at 26.1 min that has mannosylated $Trp^{250}$ in addition to O-Fuc-Glc (1329.01 *m/z*), except that the C-Man moiety remains intact, giving rise to the co-eluting 1174.98 m/z and 1248.01 m/z species. The two species with O-Fuc (1166.95 *m/z* and the C-mannosylated version, 1248.01 *m/z*) are also observed with distinct retention times (29.8 min and 26.8 min, respectively), indicating that they are present in the sample, albeit at very low levels. This indicates that the majority of O-Fuc on the TSR is extended to the O-Fuc-Glc disaccharide by endogenous B3GLCT. Both of the major species (1248.01 *m/z* and 1329.01 *m/z*) as well as co-eluting neutral loss species are also seen at very low levels as multiple peaks with similar but distinct retention times, e.g., the 1248.01 *m/z*, 1166.98 *m/z*, and 1093.95 *m/z* peaks at 28.7 min. The origin of these species is unknown, and may represent off-target activity by other endogenous glycosyltransferases. These unexpected species represent a very small fraction of the total protein signal and are assumed to have a negligible effect on the melting point assays.

**Table 1. Glycoforms positively identified by MS2 fragment spectra.**

| Glycoform | m/z (z=2) | Sample 1 | | | Sample 2 | | |
|---|---|---|---|---|---|---|---|
| | | RT (min) | Peak height | Percent | RT (min) | Peak height | Percent |
| TASCGVW[ ]DEW[ ]SPCSVT[ ]CK | 1093.95 | | | 0.00% | | | 0.00% |
| TASCGVW[ ]DEW[ ]SPCSVT[Fuc ]CK | 1166.98 | 29.78 | 9.87E+05 | 0.21% | 29.76 | 2.90E+05 | 0.13% |
| TASCGVW[Man]DEW[ ]SPCSVT[ ]CK | 1174.98 | | | 0.00% | | | 0.00% |
| TASCGVW[ ]DEW[Man]SPCSVT[ ]CK | 1174.98 | | | 0.00% | | | 0.00% |
| TASCGVW[ ]DEW[ ]SPCSVT[FucGlc]CK | 1248.01 | 28.95 | 3.67E+08 | 78.87% | 28.94 | 1.70E+08 | 76.68% |
| TASCGVW[Man]DEW[ ]SPCSVT[Fuc ]CK | 1248.01 | | | 0.00% | | | 0.00% |
| TASCGVW[ ]DEW[Man]SPCSVT[Fuc ]CK | 1248.01 | 26.82 | 3.82E+05 | 0.08% | 26.80 | 1.99E+05 | 0.09% |
| TASCGVW[Man]DEW[Man]SPCSVT[ ]CK | 1256.00 | | | 0.00% | | | 0.00% |
| TASCGVW[Man]DEW[ ]SPCSVT[FucGlc]CK | 1329.03 | | | 0.00% | | | 0.00% |
| TASCGVW[ ]DEW[Man]SPCSVT[FucGlc]CK | 1329.03 | 26.13 | 9.70E+07 | 20.83% | 26.14 | 5.12E+07 | 23.10% |
| TASCGVW[Man]DEW[Man]SPCSVT[Fuc ]CK | 1329.03 | | | 0.00% | | | 0.00% |
| TASCGVW[Man]DEW[Man]SPCSVT[FucGlc]CK | 1410.06 | | | 0.00% | | | 0.00% |

***Pf*TRAP TSR WT** **TASCGVWDEWSPCSVT[FucGlc]CGK**

| xcorr Score | Expect Score | Fragment Ions | Retention Time (sec) | Retention Time (min) | Precursor m/z | Peptide m/z | Neutral Loss |
|---|---|---|---|---|---|---|---|
| 5.423 | 2E-13 | 32/36 | 1736.9 | 28.95 | 1248.01 | 1093.95 | 308.11 |

**USI:** mzspec:PXD064887:Eclipse_2025-02-26_KES_ES903_DDAinclusion_PfTRAP_TSR-Dpy19plus_2023-06-13_4pmol:scan:8393:{dHex(1)Hex(1)}TASC[Carbamidomethyl]GVWDEWSPC[Carbamidomethyl]SVTC[Carbamidomethyl]GK/2

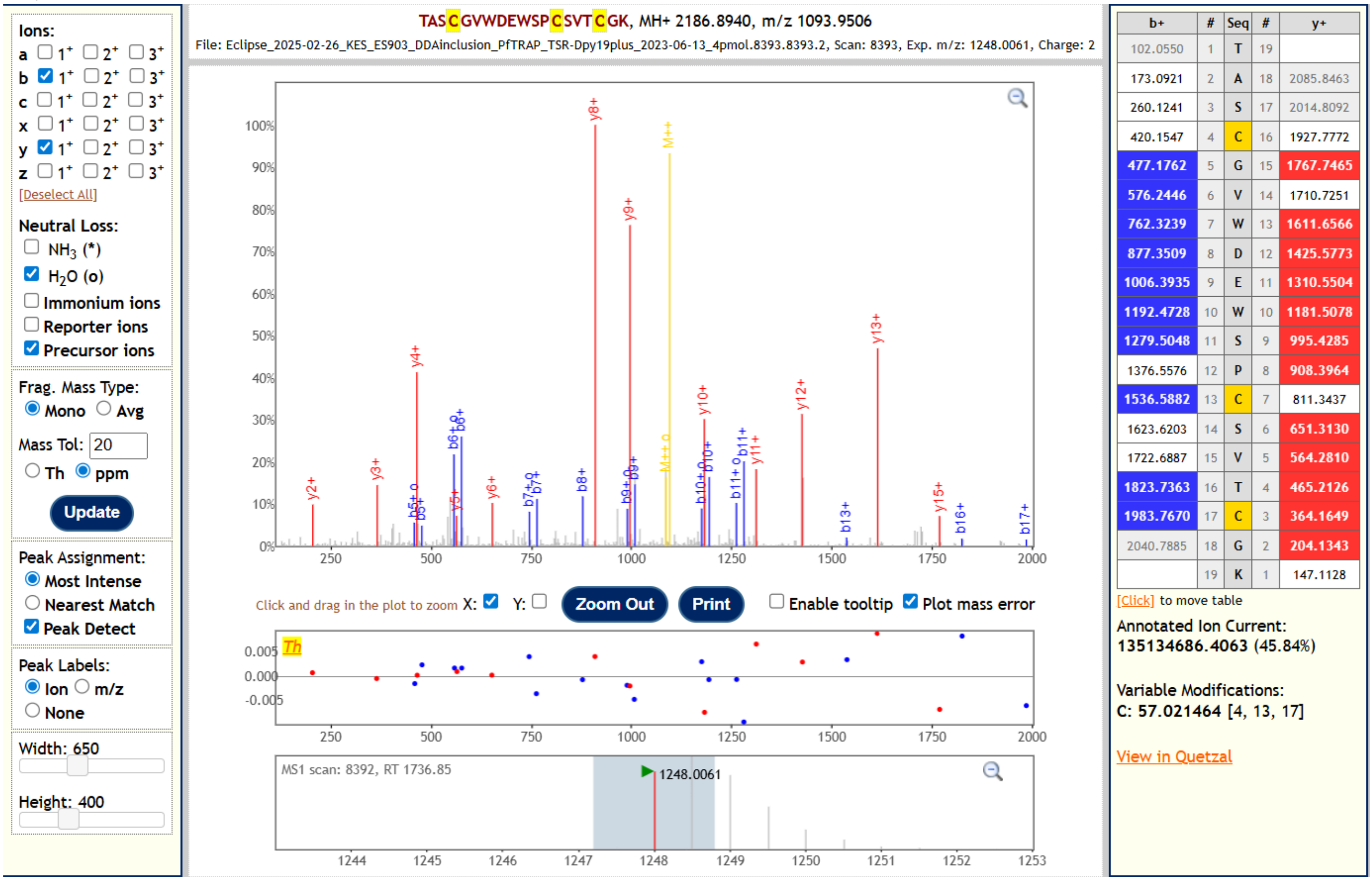


The most abundant *Pf*TRAP glycoform detected from the sample, this peptide was identified by allowing for neutral loss of 308.11 Da, consistent with the O-Fuc-Glc disaccharide. All identifying fragment ions match the unmodified peptide, and unfragmented, unmodified peptide is visible as the doubly charged species at 1093.95 *m/z* (yellow M++ peak). The localization of the glycan cannot be determined from this spectrum, but it is assumed to be Thr[256], which is a located in the canonical TSR O-fucosylation motif CXXTCXXG. The T256A mutant confirms this localization (example given below).

***Pf*TRAP TSR WT** **TASCGVWDEWSPCSVTCGK**

| xcorr Score | Expect Score | Fragment Ions | Retention Time (sec) | Retention Time (min) | Precursor m/z | Peptide m/z | Neutral Loss |
|---|---|---|---|---|---|---|---|
| 4.85 | 6.74E-12 | 32/36 | 1737.1 | 28.95 | 1093.95 | 1093.95 | 0.00 |

**USI:** mzspec:PXD064887:Eclipse_2025-02-26_KES_ES903_DDAinclusion_PfTRAP_TSR-Dpy19plus_2023-06-13_4pmol:scan:8394:TASC[Carbamidomethyl]GVWDEWSPC[Carbamidomethyl]SVTC[Carbamidomethyl]GK/2

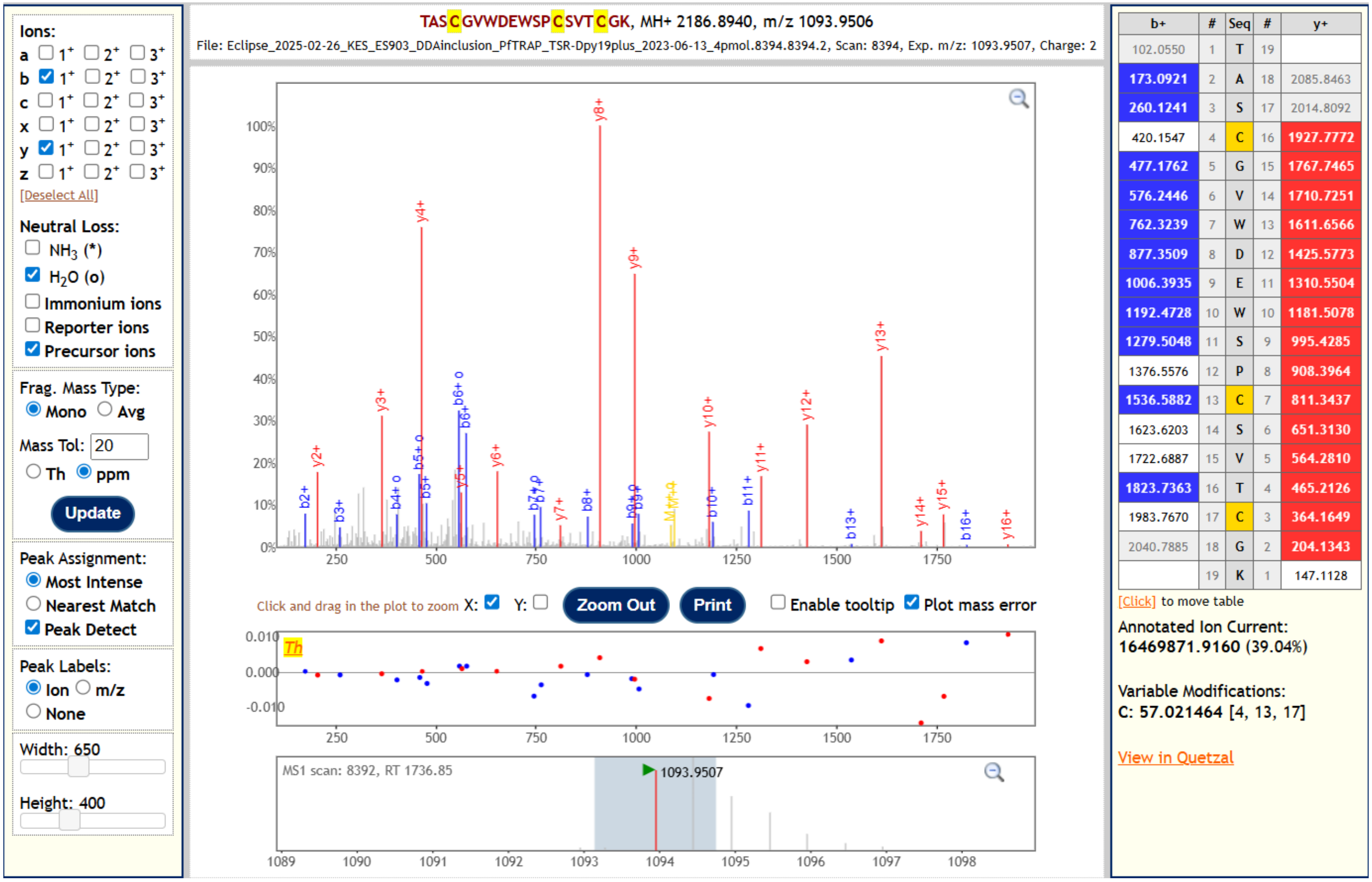


This unmodified peptide was identified from a 1093.95 *m/z* species co-eluting with the dominant 1248.01 *m/z* O-Fuc-Glc species, indicating that it arose from in-source fragmentation after eluting from the separation column.

***Pf*TRAP TSR WT** **TASCGVWDEWSPCSVT[Fuc]CGK**

| xcorr Score | Expect Score | Fragment Ions | Retention Time (sec) | Retention Time (min) | Precursor m/z | Peptide m/z | Neutral Loss |
|---|---|---|---|---|---|---|---|
| 4.842 | 9.12E-10 | 26/36 | 1787.1 | 29.79 | 1166.98 | 1093.95 | 146.06 |

**USI:** mzspec:PXD064887:Eclipse_2025-02-26_KES_ES903_DDAinclusion_PfTRAP_TSR-Dpy19plus_2023-06-13_4pmol:scan:8641:{dHex}TASC[Carbamidomethyl]GVWDEWSPC[Carbamidomethyl]SVTC[Carbamidomethyl]GK/2

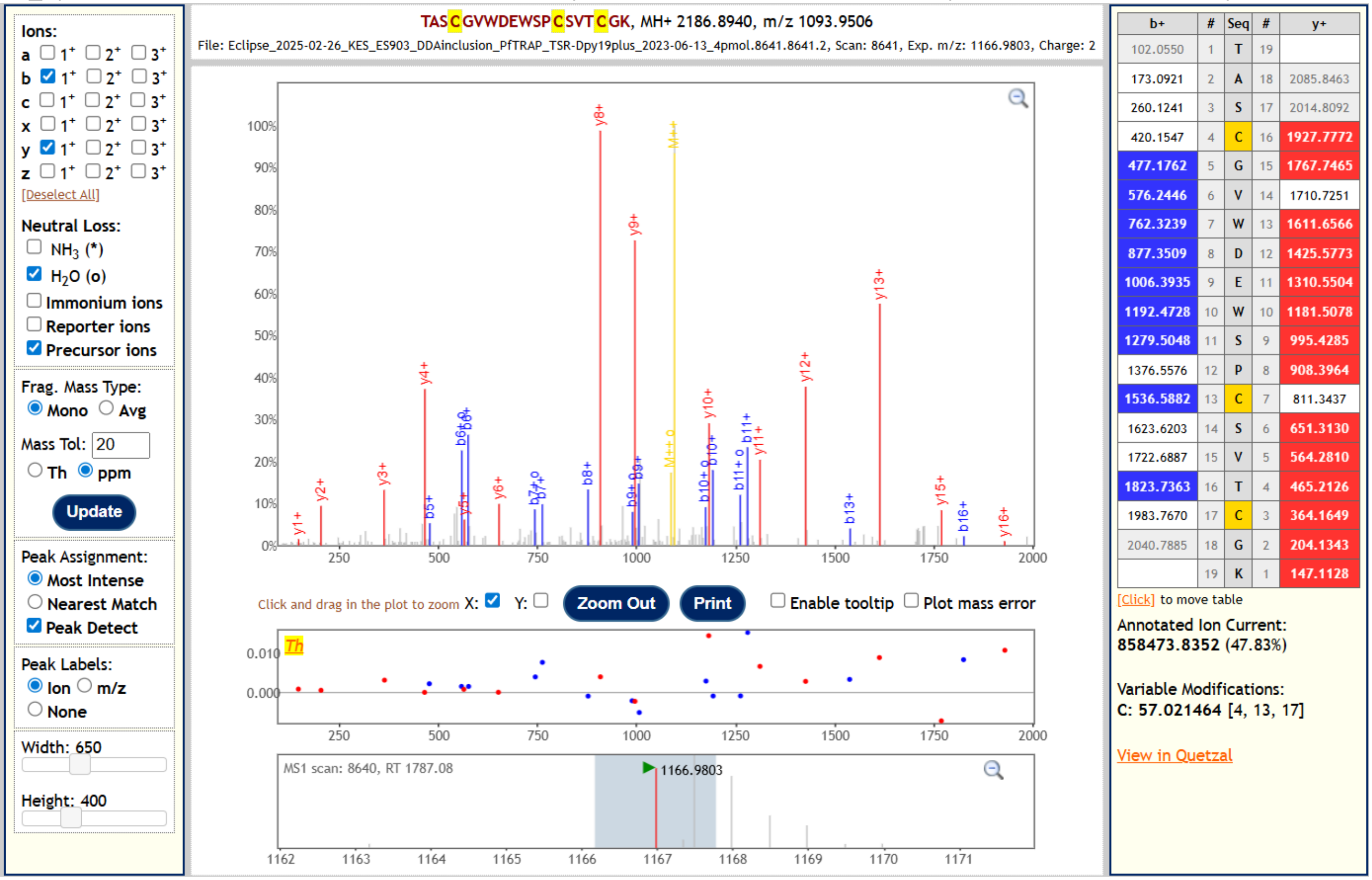


This peptide was identified by allowing for neutral loss of 146.06 Da, consistent with the O-Fuc monosaccharide. All identifying fragment ions match the unmodified peptide, and the unmodified, unfragmented peptide is visible as the charged species at 1093.95 *m/z* (yellow M++ peak). This peptide was detected at a retention time distinct from the O-Fuc-Glc peptide, indicating that it was present in the sample and did not arise from in-source fragmentation of the dominant disaccharide peptide.

***Pf*TRAP TSR WT** **TASCGVWDEW[Man]SPCSVT[FucGlc]CGK**

| xcorr Score | Expect Score | Fragment Ions | Retention Time (sec) | Retention Time (min) | Precursor m/z | Peptide m/z | Neutral Loss |
|---|---|---|---|---|---|---|---|
| 5.912 | 6.8E-11 | 29/36 | 1568.0 | 26.13 | 1329.03 | 1174.98 | 308.12 |

**USI:** mzspec:PXD064887:Eclipse_2025-02-26_KES_ES903_DDAinclusion_PfTRAP_TSR-Dpy19plus_2023-06-13_4pmol:scan:7561:{dHex(1)Hex(1)}TASC[Carbamidomethyl]GVWDEW[Hex]SPC[Carbamidomethyl]SVTC[Carbamidomethyl]GK/2

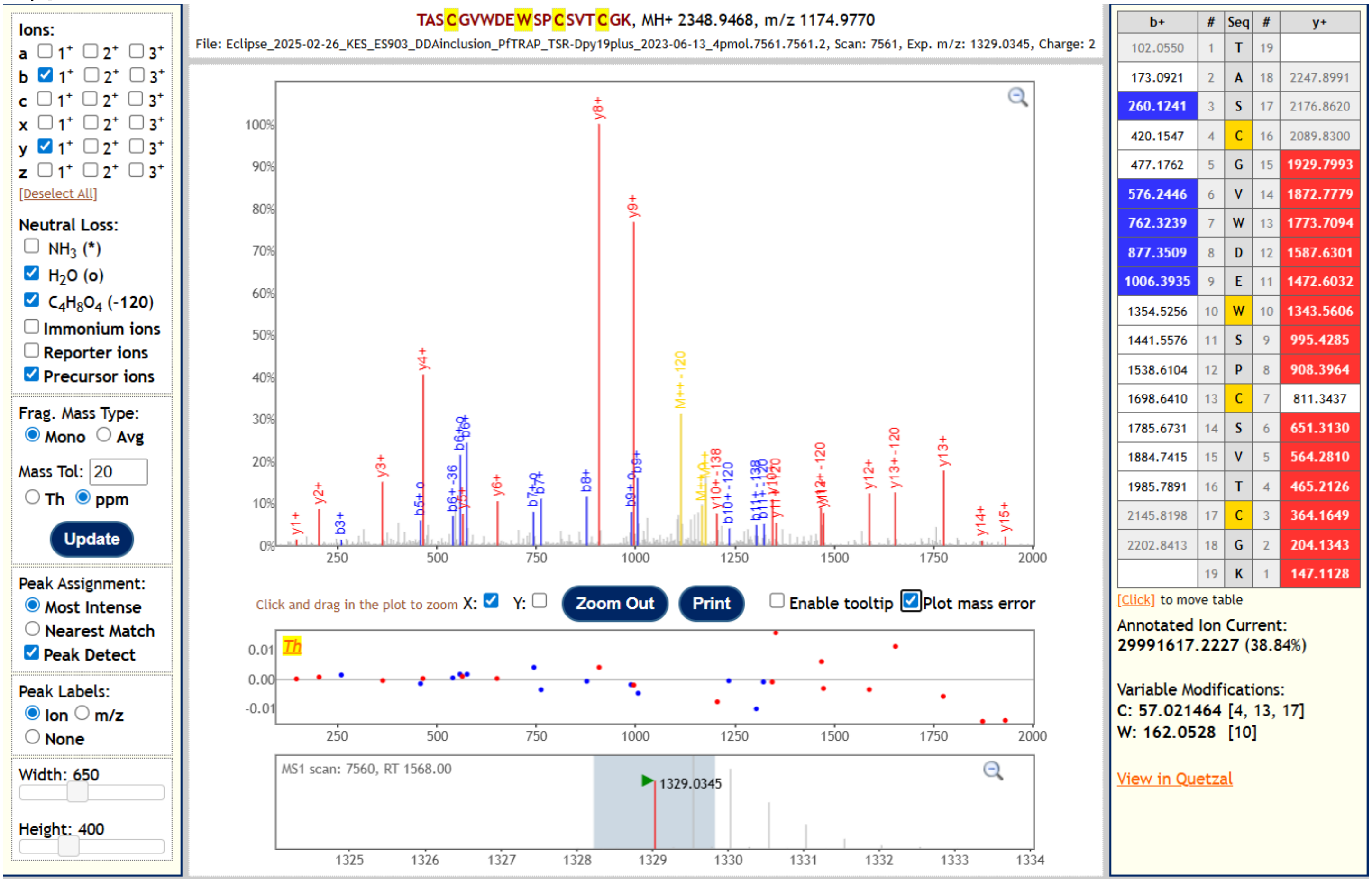


This is the second most abundant glycoform detected in the sample. The peptide was identified by allowing for neutral loss of 308.11 Da, consistent with the O-Fuc-Glc disaccharide. All identifying fragment ions match the peptide without O-Fuc-Glc, but with a Hex (i.e., C-Man), which can be localized to Trp$^{250}$. The mannosylated peptide lacking O-Fuc-Glc is visible as the doubly charged species at 1174.98 *m/z* (yellow M++ peak). Additionally, neutral loss of 120.04 Da, consistent with cross-ring cleavage of the C-Man is visible on precursor and fragment ions, further corroborating the presence and localization of the C-Man.

***Pf*TRAP TSR WT** **TASCGVWDEW[Man]SPCSVTCGK**

| xcorr Score | Expect Score | Fragment Ions | Retention Time (sec) | Retention Time (min) | Precursor m/z | Peptide m/z | Neutral Loss |
|---|---|---|---|---|---|---|---|
| 5.125 | 2.21E-11 | 29/36 | 1568.3 | 26.14 | 1174.98 | 1174.98 | 0.00 |

**USI:** mzspec:PXD064887:Eclipse_2025-02-26_KES_ES903_DDAinclusion_PfTRAP_TSR-Dpy19plus_2023-06-13_4pmol:scan:7562:TASC[Carbamidomethyl]GVWDEW[Hex]SPC[Carbamidomethyl]SVTC[Carbamidomethyl]GK/2

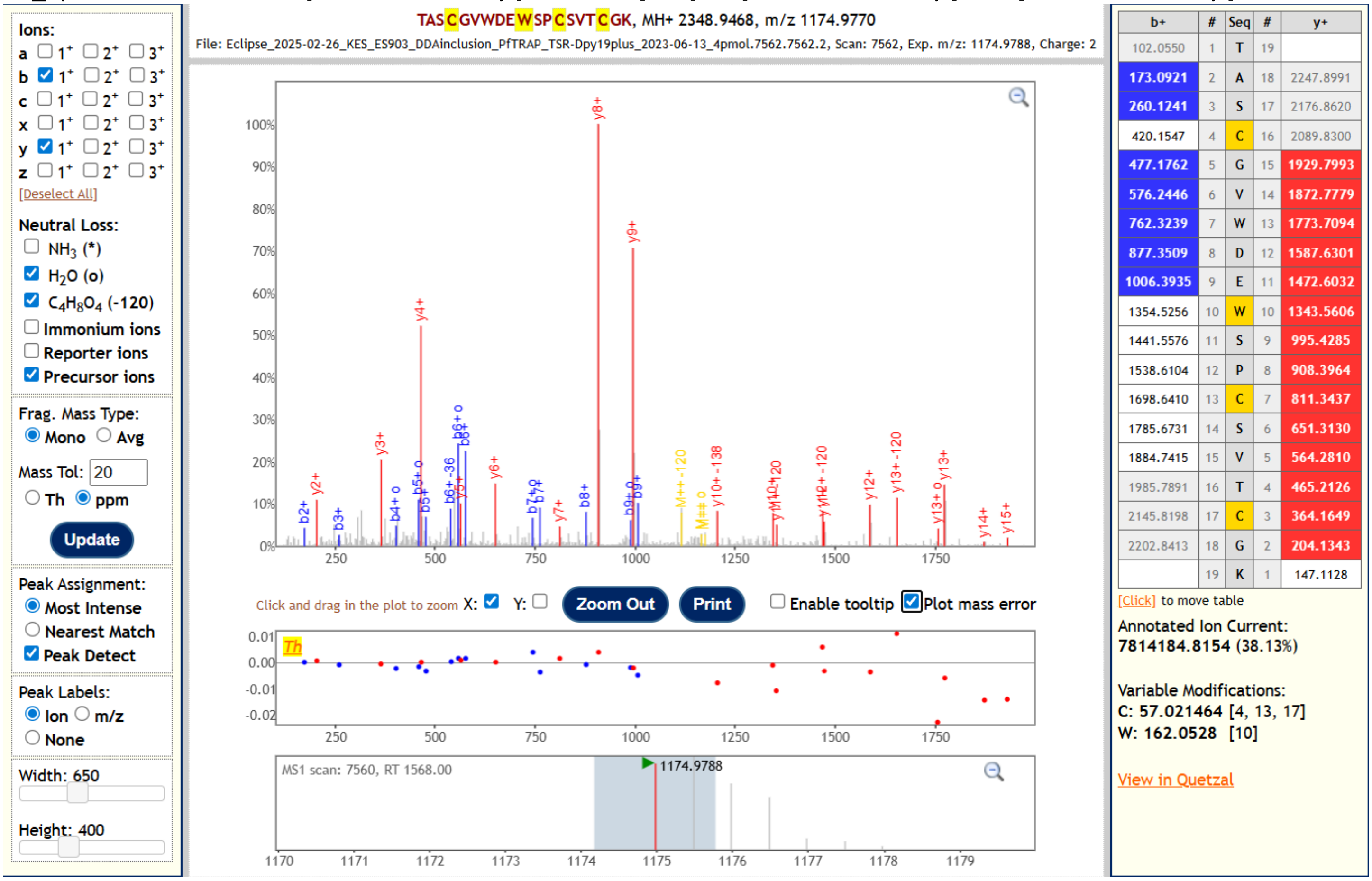


This mannosylated peptide was identified from a 1174.98 *m/z* species co-eluting with the dominant 1329.03 *m/z* C-Man + O-Fuc-Glc species, indicating that it arose from neutral loss of the O-Fuc-Glc moiety due to in-source fragmentation after eluting from the separation column. Fragment ions identify a Hex (i.e., C-Man) and localize it to Trp[250]. Neutral loss of 120.04 Da from cross-ring cleavage of the C-Man is visible on precursor and fragment ions, further corroborating the presence and localization of the C-Man.

***Pf*TRAP TSR WT** **TASCGVWDEW[Man]SPCSVT[Fuc]CGK**

| xcorr Score | Expect Score | Fragment Ions | Retention Time (sec) | Retention Time (min) | Precursor m/z | Peptide m/z | Neutral Loss |
|---|---|---|---|---|---|---|---|
| 4.737 | 6.24E-06 | 20/36 | 1610.9 | 26.85 | 1248.01 | 1174.98 | 146.06 |

**USI:** mzspec:PXD064887:Eclipse_2025-02-26_KES_ES903_DDAinclusion_PfTRAP_TSR-Dpy19plus_2023-06-13_4pmol:scan:7777:{dHex}TASC[Carbamidomethyl]GVWDEW[Hex]SPC[Carbamidomethyl]SVTC[Carbamidomethyl]GK/2

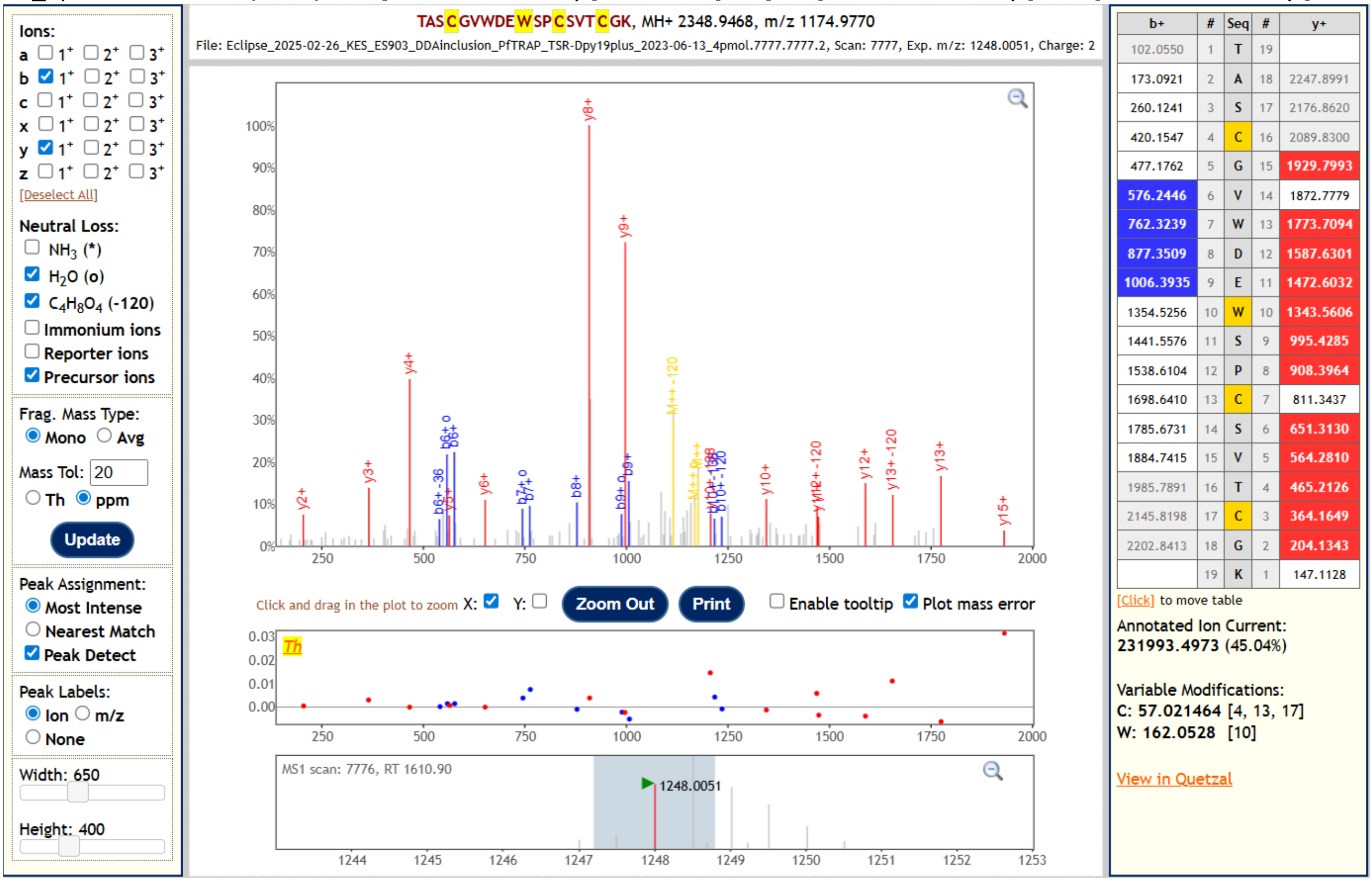


This mannosylated peptide was identified by allowing for neutral loss of 146.06 Da, consistent with the O-Fuc monosaccharide. All identifying fragment ions match the peptide without O-Fuc-Glc, but with a Hex (i.e., C-Man), which can be localized to Trp[250]. The mannosylated peptide lacking O-Fuc is visible as the doubly charged species at 1174.98 *m/z* (yellow M++ peak). Additionally, neutral loss of 120.04 Da, consistent with cross-ring cleavage of the C-Man is visible on precursor and fragment ions, further corroborating the presence and localization of the C-Man. This species was detected at a retention time distinct from the C-Man + O-Fuc-Glc species, indicating that it was present in the sample and did not arise from in-source fragmentation of the dominant disaccharide peptide.

#### 2. *Pf*TRAP TSR WT expressed in CHO DPY19⁻
#### Glycopeptide sequence: TASCGVWDEWSPCSVTCGK

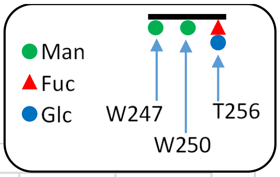


Sample 1 (2023-08-15)    Sample 2 (2023-08-30)

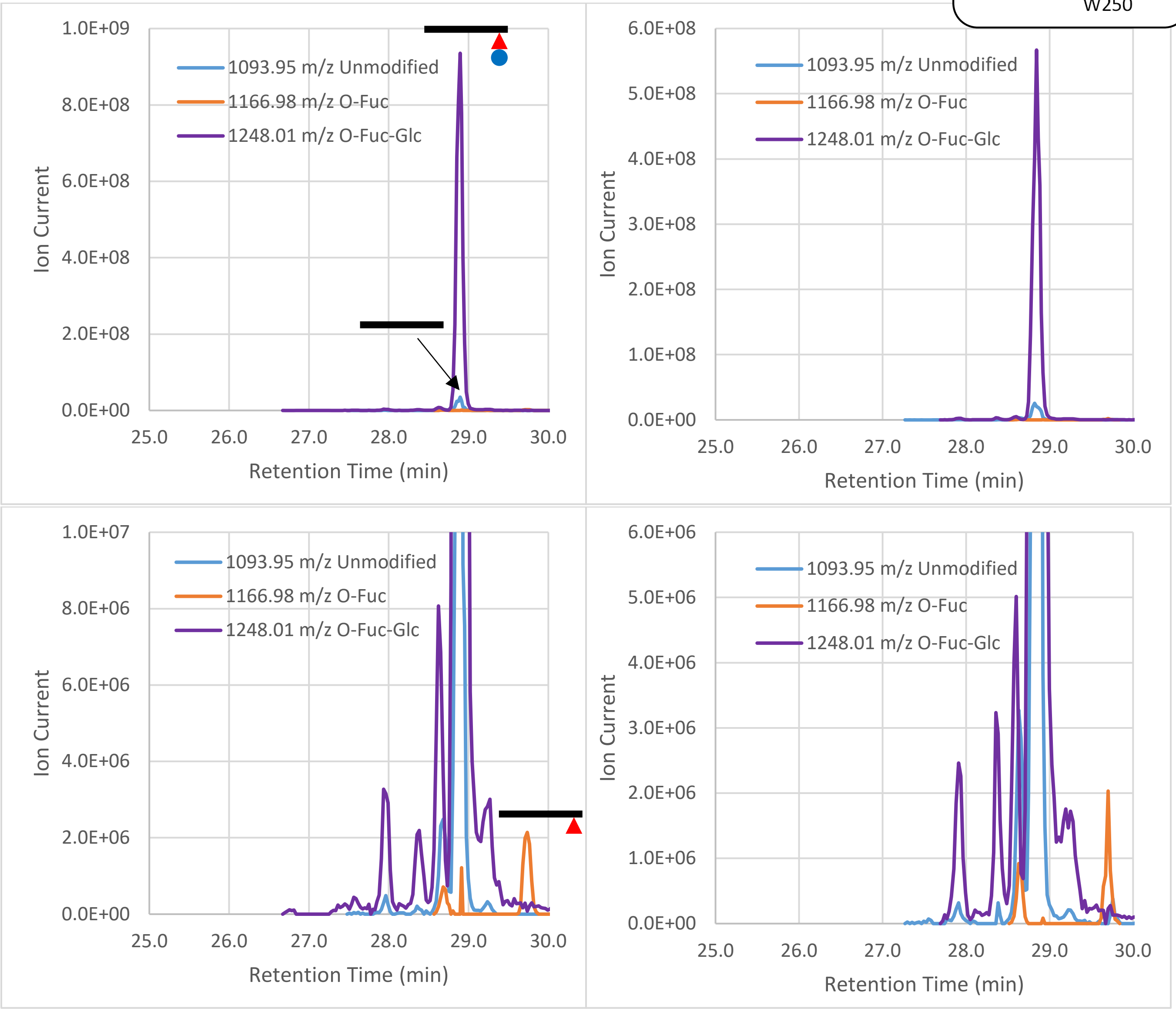


**Fig 2. Extracted ion chromatograms of the detected glycopeptides.** Left is Sample 1, right is Sample 2. Bottom plots are zoomed in on the y-axis to show low-abundance species. The unmodified peptide (1093.95 *m/z*) and the O-Fuc peptide (1166.95 *m/z*) co-eluting with the dominant O-Fuc-Glc peptide (1248.01 *m/z*) at 28.9 min arise from neutral loss of the O-glycan due to in-source fragmentation that occurs after the peptide has eluted from the column. No mannosylated peptide was detected in the samples, consistent with absence of endogenous DPY19, and confirming that *Pf*DPY19 alone is responsible for the C-Man observed on the protein expressed in cell lines transduced with that enzyme. The O-Fuc species (1166.95 *m/z*) is also observed with distinct retention time (29.7 min) indicating that it is present in the sample, albeit at very low levels. The major species (1248.01 *m/z*) as well as co-eluting neutral loss species are also seen at very low levels as multiple peaks with similar but distinct retention times, e.g., the 1248.01 *m/z*, 1166.98 *m/z*, and 1093.95 *m/z* peaks at 28.7 min. The origin of these species is unknown, and may represent off-target activity by other endogenous glycosyltransferases. These unexpected species represent a very small fraction of the total protein signal and are assumed to have a negligible effect on the melting point assays.

**Table 2. Glycoforms positively identified by MS2 fragment spectra.**

| Glycoform | m/z (z=2) | Sample 1 | | | Sample 2 | | |
|---|---|---|---|---|---|---|---|
| | | RT (min) | Peak height | Percent | RT (min) | Peak height | Percent |
| TASCGVW[ ]DEW[ ]SPCSVT[ ]CK | 1093.95 | | | 0.00% | | | 0.00% |
| TASCGVW[ ]DEW[ ]SPCSVT[Fuc ]CK | 1166.98 | 29.74 | 2.14E+06 | 0.23% | 29.70 | 2.03E+06 | 0.36% |
| TASCGVW[Man]DEW[ ]SPCSVT[ ]CK | 1174.98 | | | 0.00% | | | 0.00% |
| TASCGVW[ ]DEW[Man]SPCSVT[ ]CK | 1174.98 | | | 0.00% | | | 0.00% |
| TASCGVW[ ]DEW[ ]SPCSVT[FucGlc]CK | 1248.01 | 28.89 | 9.35E+08 | 99.77% | 28.84 | 5.67E+08 | 99.64% |
| TASCGVW[Man]DEW[ ]SPCSVT[Fuc ]CK | 1248.01 | | | 0.00% | | | 0.00% |
| TASCGVW[ ]DEW[Man]SPCSVT[Fuc ]CK | 1248.01 | | | 0.00% | | | 0.00% |
| TASCGVW[Man]DEW[Man]SPCSVT[ ]CK | 1256.00 | | | 0.00% | | | 0.00% |
| TASCGVW[Man]DEW[ ]SPCSVT[FucGlc]CK | 1329.03 | | | 0.00% | | | 0.00% |
| TASCGVW[ ]DEW[Man]SPCSVT[FucGlc]CK | 1329.03 | | | 0.00% | | | 0.00% |
| TASCGVW[Man]DEW[Man]SPCSVT[Fuc ]CK | 1329.03 | | | 0.00% | | | 0.00% |
| TASCGVW[Man]DEW[Man]SPCSVT[FucGlc]CK | 1410.06 | | | 0.00% | | | 0.00% |

#### 3. *Pf*TRAP TSR W250F expressed in CHO DPY19$^{-}$ transduced with *Pfdpy19*

#### Glycopeptide sequence: TASCGVWDEFSPCSVTCGK

Sample 1 (2023-06-13) Sample 3 (2024-02-28)

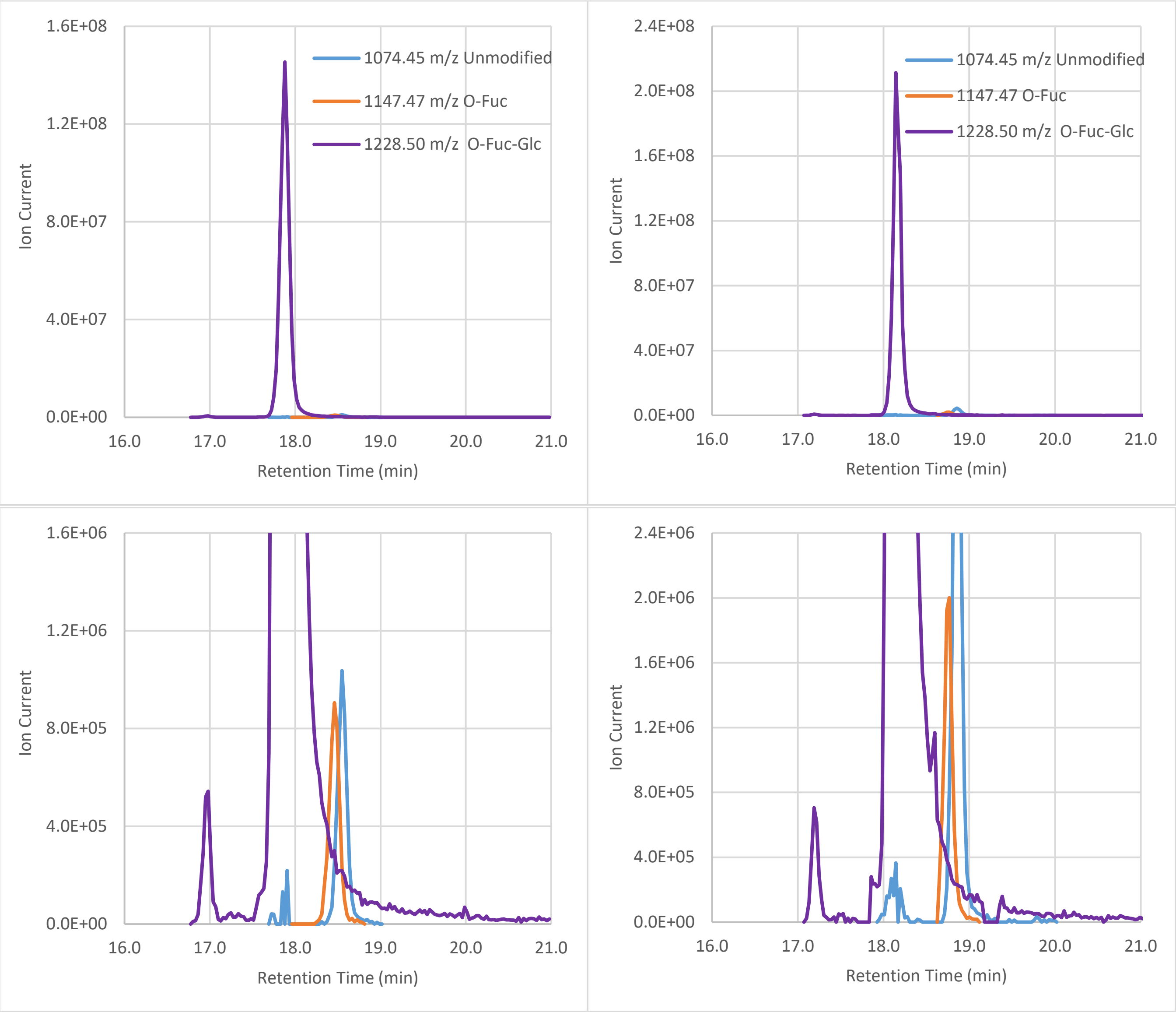


**Fig 3. Extracted ion chromatograms of the detected glycopeptides.** Left is Sample 1, right is Sample 3. Bottom plots are zoomed in on the y-axis to show low-abundance species. No mannosylated peptide was detected in the samples, confirming that the W/F substitution at Trp$^{250}$ prevented addition of C-Man at that position by *Pf*DPY19. A trace amount of the unmodified peptide (1074.45 *m/z*) co-elutes with the dominant O-Fuc-Glc peptide (1228.50 *m/z*), indicating neutral loss of the O-glycan due to in-source fragmentation that occurs after the peptide has eluted from the column. Both the unmodified peptide (1074.45 *m/z*) and the O-Fuc species (1166.95 *m/z*) are observed with distinct retention times, indicating that they are present in the sample, albeit at very low levels. As seen in the WT samples above, the major species (1228.50 *m/z*) is also seen at very low levels as multiple peaks with similar but distinct retention times. The origin of these species is unknown, and may represent off-target activity by other endogenous glycosyltransferases. These unexpected species represent a very small fraction of the total protein signal and are assumed to have a negligible effect on the melting point assays.

**Table 3. Glycoforms positively identified by MS2 fragment spectra.**

| Glycoform | m/z (z=2) | Sample 1 | | | Sample 3 | | |
|---|---|---|---|---|---|---|---|
| | | RT (min) | Peak height | Percent | RT (min) | Peak height | Percent |
| TASCGVW[ ]DEF[ ]SPCSVT[ ]CK | 1074.45 | 18.55 | 1.04E+06 | 0.70% | 18.86 | 4.46E+06 | 2.05% |
| TASCGVW[ ]DEF[ ]SPCSVT[Fuc ]CK | 1147.47 | 18.46 | 9.06E+05 | 0.61% | 18.77 | 2.00E+06 | 0.92% |
| TASCGVW[Man]DEF[ ]SPCSVT[ ]CK | 1155.47 | | | 0.00% | | | 0.00% |
| TASCGVW[ ]DEF[Man]SPCSVT[ ]CK | 1155.47 | | | 0.00% | | | 0.00% |
| TASCGVW[ ]DEF[ ]SPCSVT[FucGlc]CK | 1228.50 | 17.88 | 1.45E+08 | 98.68% | 18.14 | 2.11E+08 | 97.03% |
| TASCGVW[Man]DEF[ ]SPCSVT[Fuc ]CK | 1228.50 | | | 0.00% | | | 0.00% |
| TASCGVW[ ]DEF[Man]SPCSVT[Fuc ]CK | 1228.50 | | | 0.00% | | | 0.00% |
| TASCGVW[Man]DEF[Man]SPCSVT[ ]CK | 1236.50 | | | 0.00% | | | 0.00% |
| TASCGVW[Man]DEF[ ]SPCSVT[FucGlc]CK | 1309.53 | | | 0.00% | | | 0.00% |
| TASCGVW[ ]DEF[Man]SPCSVT[FucGlc]CK | 1309.53 | | | 0.00% | | | 0.00% |
| TASCGVW[Man]DEF[Man]SPCSVT[Fuc ]CK | 1309.53 | | | 0.00% | | | 0.00% |
| TASCGVW[Man]DEF[Man]SPCSVT[FucGlc]CK | 1390.55 | | | 0.00% | | | 0.00% |

***Pf*TRAP TSR W250F** **TASCGVWDEFSPCSVT[FucGlc]CGK**

| xcorr Score | Expect Score | Fragment Ions | Retention Time (sec) | Retention Time (min) | Precursor m/z | Peptide m/z | Neutral Loss |
|---|---|---|---|---|---|---|---|
| 5.466 | 1.05E-11 | 30/36 | 1088.6 | 18.14 | 1228.50 | 1074.45 | 308.11 |

**USI:** mzspec:PXD064887:isbECLIPSE_2024-04-01_ES904_PfTRAP-TSR_W250F_409-SC2_5pmol:scan:5252:{dHex(1)Hex(1)}TASC[Carbamidomethyl]GVWDEFSPC[Carbamidomethyl]SVTC[Carbamidomethyl]GK/2

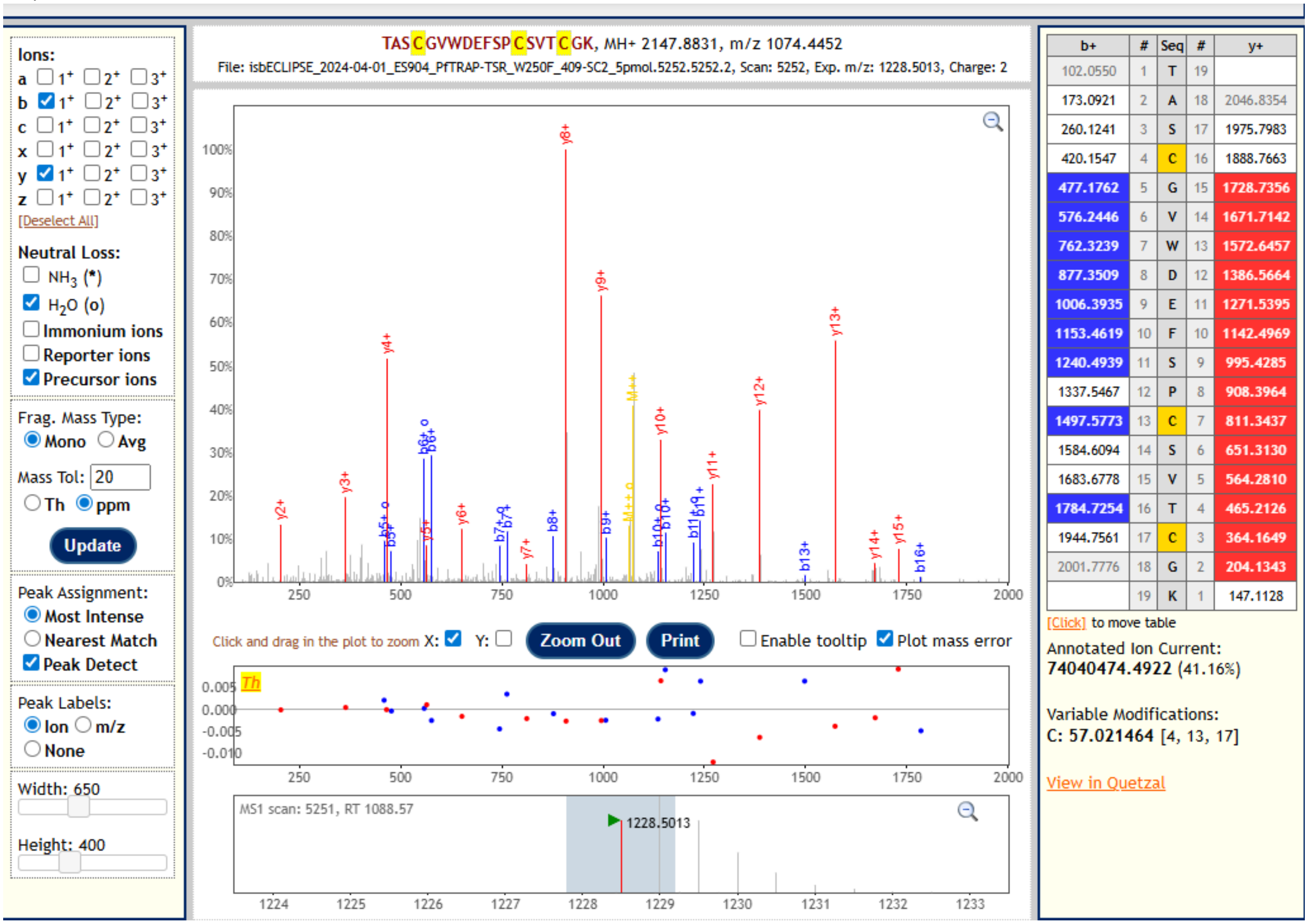


The most abundant *Pf*TRAP TSR glycoform detected from the sample, this peptide was identified by allowing for neutral loss of 308.11 Da, consistent with the O-Fuc-Glc disaccharide. All identifying fragment ions match the unmodified peptide, and the and the unmodified, unfragmented peptide is visible as the doubly charged species at at 1074.45 *m/z* (yellow M++ peak).

***Pf*TRAP TSR W250F** **TASCGVWDEFSPCSVT[Fuc]CGK**

| xcorr Score | Expect Score | Fragment Ions | Retention Time (sec) | Retention Time (min) | Precursor m/z | Peptide m/z | Neutral Loss |
|---|---|---|---|---|---|---|---|
| 5.676 | 1.23E-13 | 31/36 | 1126.1 | 18.77 | 1147.48 | 1074.45 | 146.06 |

**USI:** mzspec:PXD064887:isbECLIPSE_2024-04-01_ES904_PfTRAP-TSR_W250F_409-SC2_5pmol:scan:5428:{dHex}TASC[Carbamidomethyl]GVWDEFSPC[Carbamidomethyl]SVTC[Carbamidomethyl]GK/2

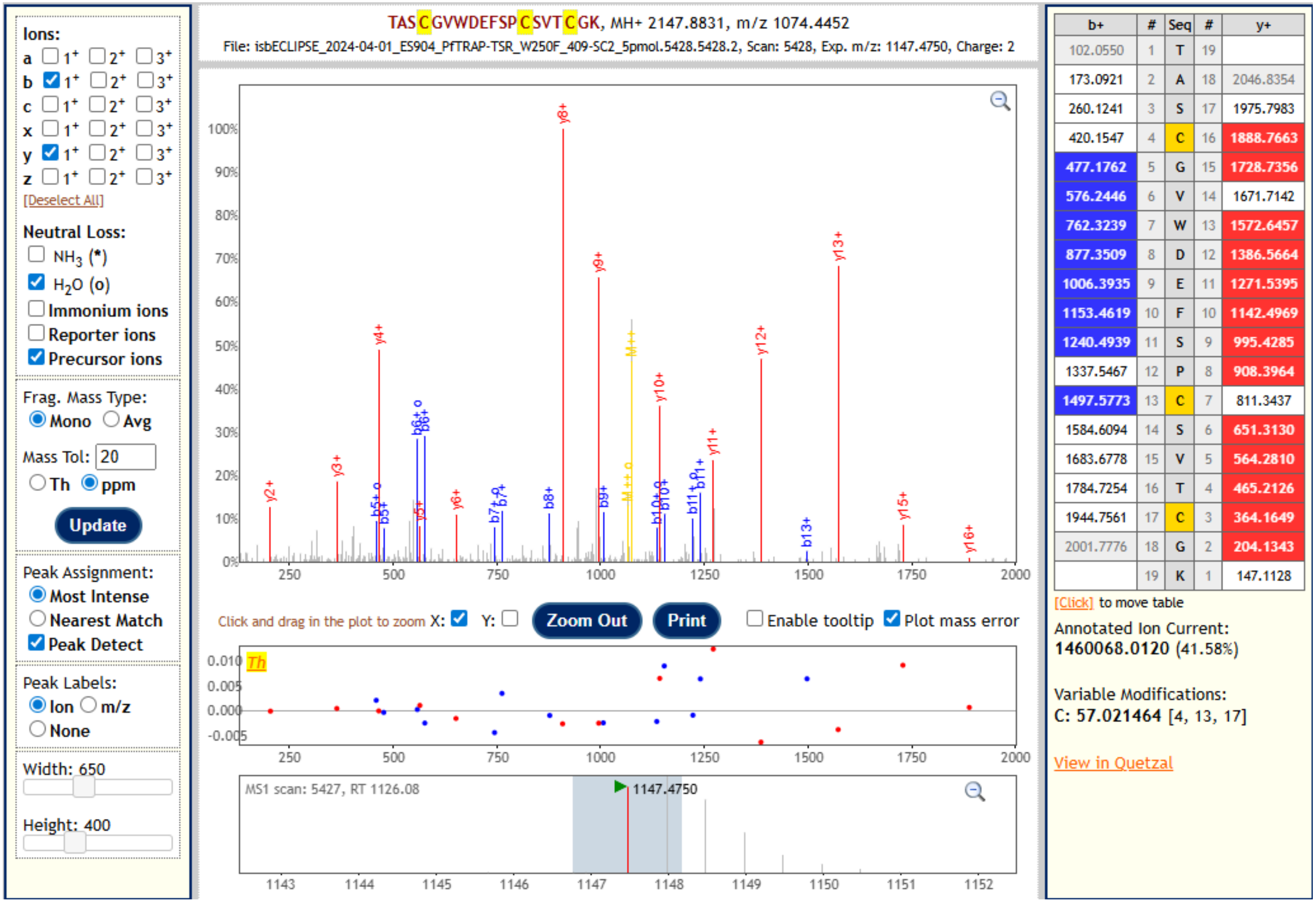


This peptide was identified by allowing for neutral loss of 146.06 Da, consistent with the O-Fuc monosaccharide. All identifying fragment ions match the unmodified peptide, and the unmodified, unfragmented peptide is visible as the doubly charged species at 1147.48 *m/z* (yellow M++ peak). This peptide was detected at a retention time distinct from the O-Fuc-Glc species, indicating that it was present in the sample and did not arise from in-source fragmentation of the dominant disaccharide peptide.

***Pf*TRAP TSR W250F** **TASCGVWDEFSPCSVTCGK**

| xcorr Score | Expect Score | Fragment Ions | Retention Time (sec) | Retention Time (min) | Precursor m/z | Peptide m/z | Neutral Loss |
|---|---|---|---|---|---|---|---|
| 5.187 | 4.68E-13 | 29/36 | 1131.3 | 18.86 | 1074.45 | 1074.45 | 0.00 |

**USI:** mzspec:PXD064887:isbECLIPSE_2024-04-01_ES904_PfTRAP-TSR_W250F_409-SC2_5pmol:scan:5452:TASC[Carbamidomethyl]GVWDEFSPC[Carbamidomethyl]SVTC[Carbamidomethyl]GK/2

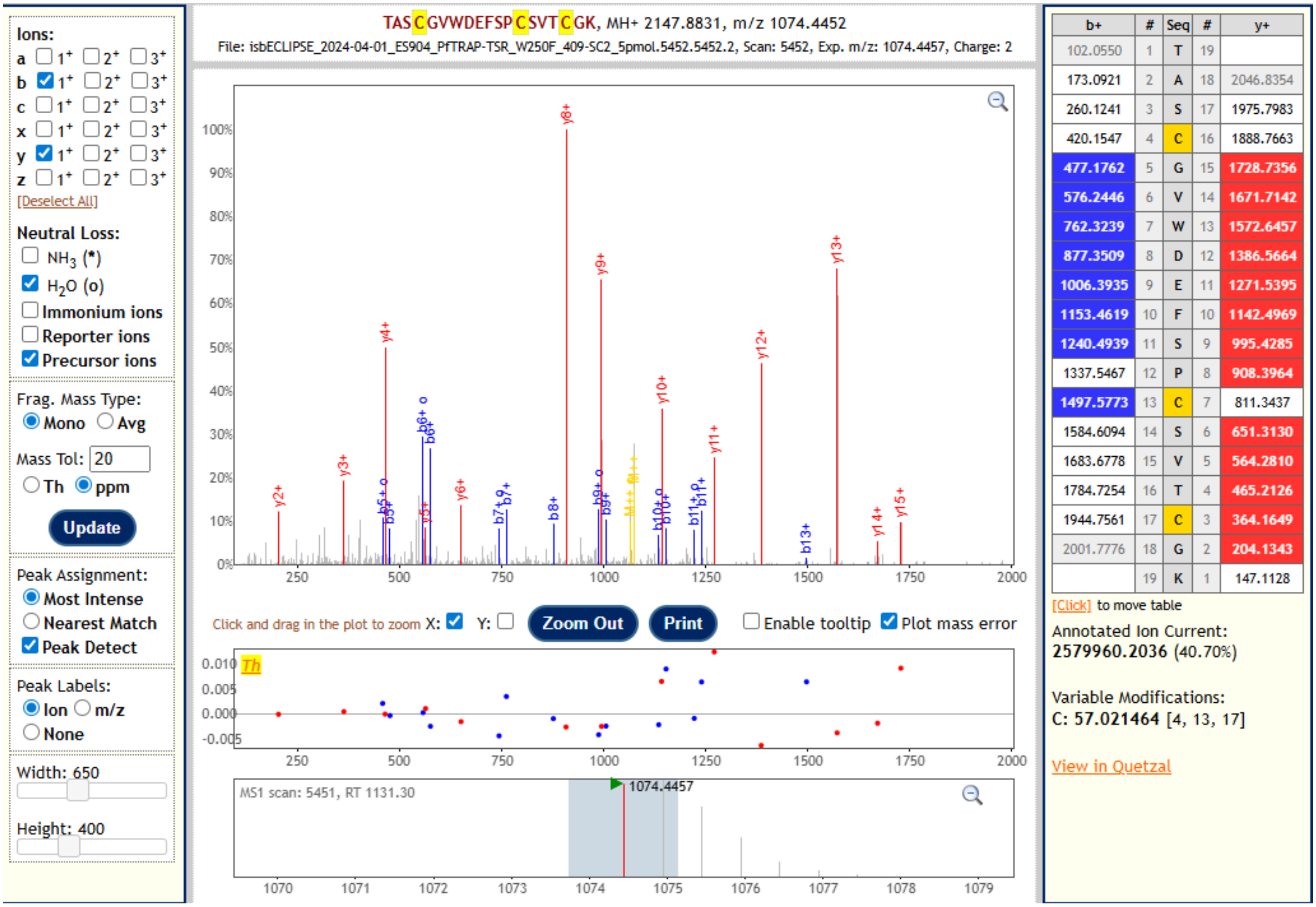


This unmodified peptide was detected at a retention time distinct from the O-Fuc-Glc species, indicating that it was present in the sample and did not arise from in-source fragmentation of the dominant disaccharide or the minor O-Fuc peptide.

**4. *Pf*TRAP TSR WT expressed in CHO DPY19⁻/POFUT2⁻ transduced with *Pfdpy19***

**Glycopeptide sequence: TASCGVWDEWSPCSVTCGK**

Sample 1 (2024-07-02)

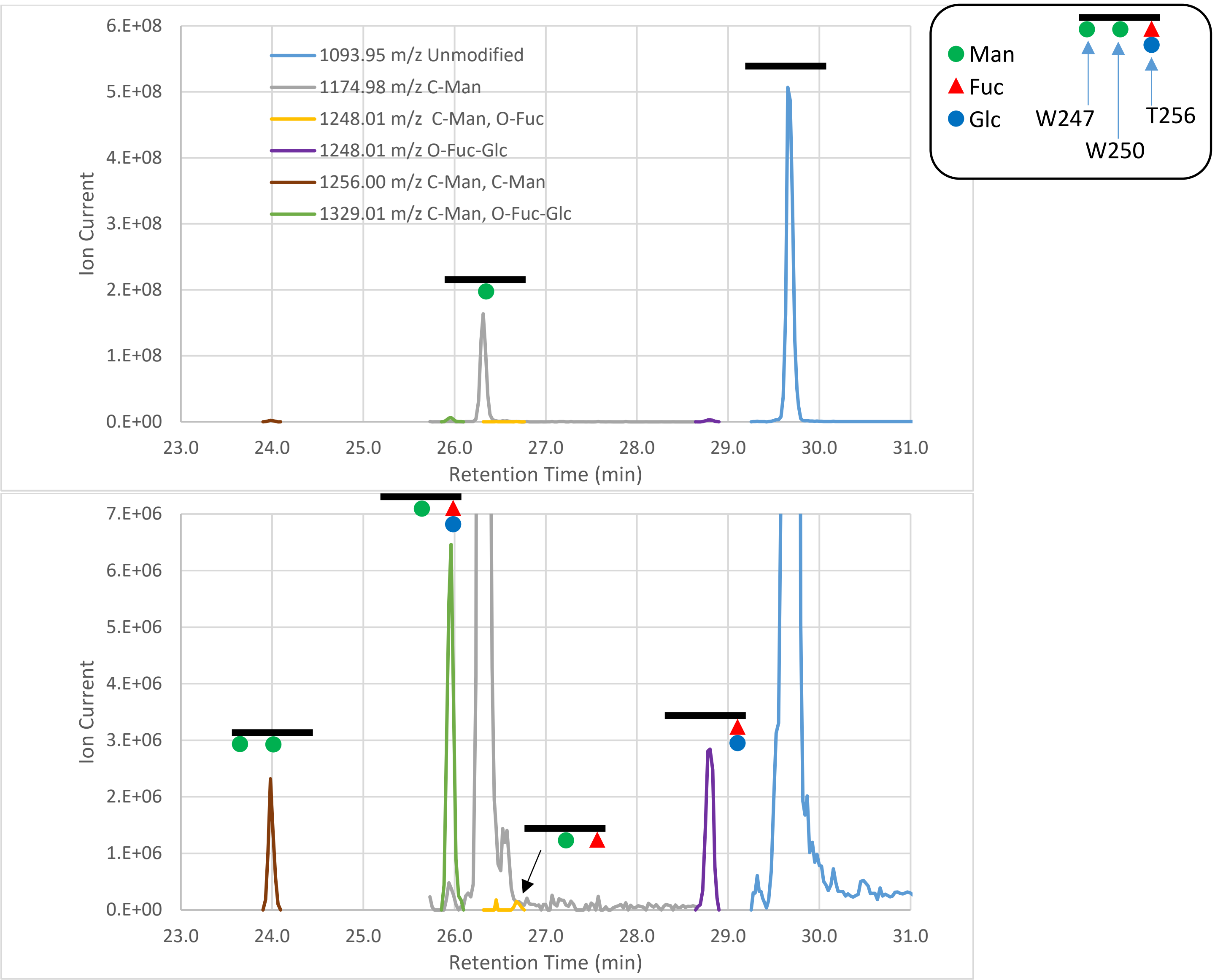


**Fig 4. Extracted ion chromatograms of the detected glycopeptides.** The bottom plot is zoomed in on the y-axis to show low-abundance species. The dominant species are the completely unmodified peptide and the peptide with C-Man at Trp$^{250}$ but no O-Fuc or O-Fuc-Glc, consistent with abrogation of POFUT2 activity. Trace amounts of peptide with O-Fuc and O-Fuc-Glc are detected at the expected retention times (based on the XIC of *Pf*TRAP TSR WT from parental line detailed above), suggesting that POFUT2 disruption was not complete, but nearly so. In addition to the expected glycoforms, the peptide was also observed with a mass matching addition of two Hex moieties (1256.00 *m/z*). MS2 spectra (example below) confirm that this species is modified with a second C-Man at Trp$^{247}$ in addition to the expected C-Man at Trp$^{250}$.

**Table 4. Glycoforms positively identified by MS2 fragment spectra.**

| Glycoform | m/z (z=2) | Sample 1 | | |
|---|---|---|---|---|
| | | RT (min) | Peak height | Percent |
| TASCGVW[ ]DEW[ ]SPCSVT[ ]CK | 1093.95 | 29.65 | 5.07E+08 | 74.28% |
| TASCGVW[ ]DEW[ ]SPCSVT[Fuc ]CK | 1166.98 | | | 0.00% |
| TASCGVW[Man]DEW[ ]SPCSVT[ ]CK | 1174.98 | | | 0.00% |
| TASCGVW[ ]DEW[Man]SPCSVT[ ]CK | 1174.98 | 26.32 | 1.64E+08 | 23.99% |
| TASCGVW[ ]DEW[ ]SPCSVT[FucGlc]CK | 1248.01 | 28.80 | 2.84E+06 | 0.42% |
| TASCGVW[Man]DEW[ ]SPCSVT[Fuc ]CK | 1248.01 | | | 0.00% |
| TASCGVW[ ]DEW[Man]SPCSVT[Fuc ]CK | 1248.01 | 26.68 | 1.53E+05 | 0.02% |
| TASCGVW[Man]DEW[Man]SPCSVT[ ]CK | 1256.00 | 23.98 | 2.32E+06 | 0.34% |
| TASCGVW[Man]DEW[ ]SPCSVT[FucGlc]CK | 1329.03 | | | 0.00% |
| TASCGVW[ ]DEW[Man]SPCSVT[FucGlc]CK | 1329.03 | 25.96 | 6.46E+06 | 0.95% |
| TASCGVW[Man]DEW[Man]SPCSVT[Fuc ]CK | 1329.03 | | | 0.00% |
| TASCGVW[Man]DEW[Man]SPCSVT[FucGlc]CK | 1410.06 | | | 0.00% |

***Pf*TRAP TSR WT** **TASCGVW[Man]DEW[Man]SPCSVTCGK**

| xcorr Score | Expect Score | Fragment Ions | Retention Time (sec) | Retention Time (min) | Precursor m/z | Peptide m/z | Neutral Loss |
|---|---|---|---|---|---|---|---|
| 4.644 | 4.28E-09 | 23/36 | 1438.9 | 23.98 | 1256.00 | 1256.00 | 0.00 |

**USI:** mzspec:PXD064887:Eclipse_2025-02-28_KES_ES903_DDAinclusion_PfTRAP_TSR-Dpy19plus-POFUT2minus_4uL:scan:7494:TASC[Carbamidomethyl]GVW[Hex]DEW[Hex]SPC[Carbamidomethyl]SVTC[Carbamidomethyl]GK/2

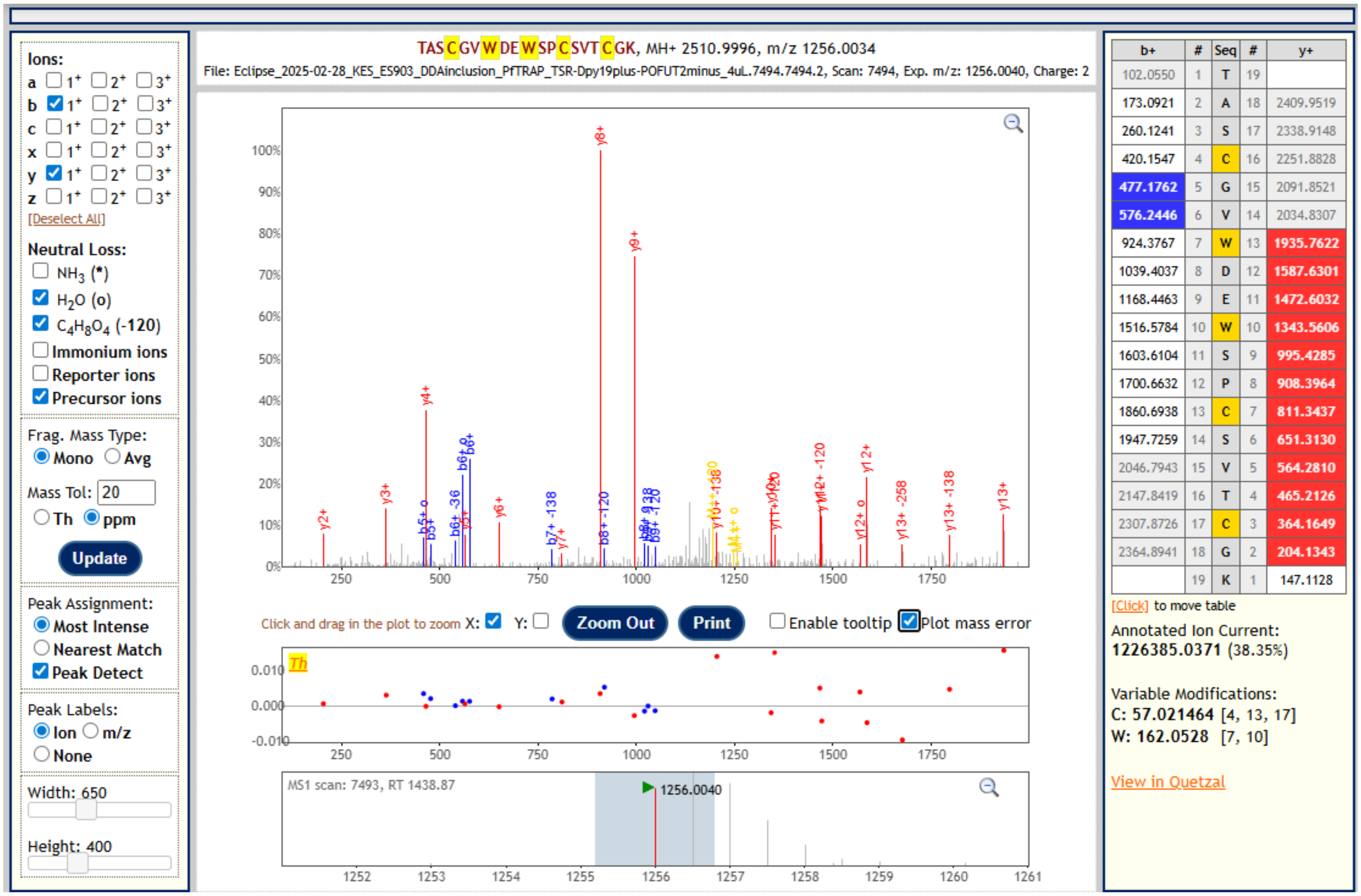


The precursor mass of this peptide corresponds to addition of two Hex moieties to the peptide. Fragment ions and characteristic neutral loss of 120.04 Da from cross-ring cleavage of C-Man confirm the presence of C-Man at Trp[247] and Trp[250].

#### 5. *Pf*TRAP TSR T256A expressed in CHO DPY19⁻ transduced with *Pfdpy19*

#### Glycopeptide sequence: TASCGVWDEWSPCSV<u>A</u>CGK

Sample 1 (2024-10-22)

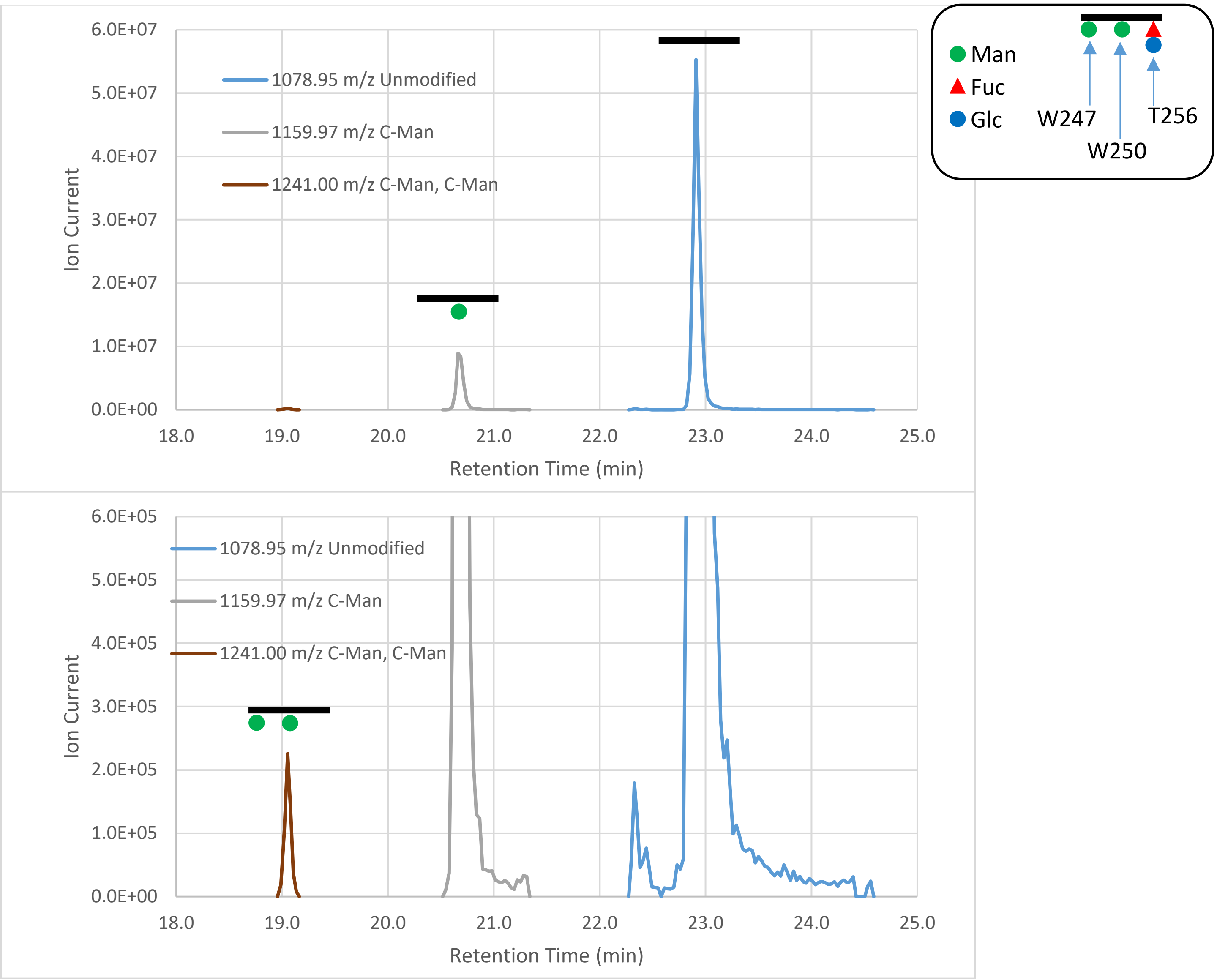


**Fig 5. Extracted ion chromatograms of the detected glycopeptides.** The bottom plot is zoomed in on the y-axis to show low-abundance species. The dominant species are the unmodified peptide and the peptide with C-Man at $Trp^{250}$ but no O-Fuc or O-Fuc-Glc. Peptide modified with O-Fuc-Glc, the dominant species in the WT and W250F constructs, is completely absent, confirming that T256 is the site of O-fucosylation recognized by POFUT2. In addition to the expected glycoforms, and similar to the *Pf*TRAP TSR WT variant expressed in a cell line with disrupted POFUT2, the peptide was also observed with a mass matching addition of two Hex moieties (1241.00 *m/z*). MS2 spectra (example below) confirm that this species is modified with a second C-Man at $Trp^{247}$ in addition to the expected C-Man at $Trp^{250}$.

| Glycoform | m/z (z=2) | Sample 1 | | |
|---|---|---|---|---|
| | | RT (min) | Peak height | Percent |
| TASCGVW[ ]DEW[ ]SPCSVA[ ]CK | 1078.95 | 22.91 | 5.53E+07 | 85.79% |
| TASCGVW[ ]DEW[ ]SPCSVA[Fuc ]CK | 1151.98 | | | 0.00% |
| TASCGVW[Man]DEW[ ]SPCSVA[ ]CK | 1159.97 | | | 0.00% |
| TASCGVW[ ]DEW[Man]SPCSVA[ ]CK | 1159.97 | 20.66 | 8.93E+06 | 13.86% |
| TASCGVW[ ]DEW[ ]SPCSVA[FucGlc]CK | 1233.00 | | | 0.00% |
| TASCGVW[Man]DEW[ ]SPCSVA[Fuc ]CK | 1233.00 | | | 0.00% |
| TASCGVW[ ]DEW[Man]SPCSVA[Fuc ]CK | 1233.00 | | | 0.00% |
| TASCGVW[Man]DEW[Man]SPCSVA[ ]CK | 1241.00 | 19.05 | 2.26E+05 | 0.35% |
| TASCGVW[Man]DEW[ ]SPCSVA[FucGlc]CK | 1314.03 | | | 0.00% |
| TASCGVW[ ]DEW[Man]SPCSVA[FucGlc]CK | 1314.03 | | | 0.00% |
| TASCGVW[Man]DEW[Man]SPCSVA[Fuc ]CK | 1314.03 | | | 0.00% |
| TASCGVW[Man]DEW[Man]SPCSVA[FucGlc]CK | 1395.05 | | | 0.00% |

***Pf*TRAP TSR T256A** **TASCGVWDEWSPCSVACGK**

| xcorr Score | Expect Score | Fragment Ions | Retention Time (sec) | Retention Time (min) | Precursor m/z | Peptide m/z | Neutral Loss |
|---|---|---|---|---|---|---|---|
| 4.642 | 7.47E-13 | 22/36 | 1376.3 | 22.94 | 1078.94 | 1078.95 | -0.01 |

**USI:** mzspec:PXD064887:isbECLIPSE_2024-10-23_KES_ES903_20minDDAinclusion_PfTRAP-TSR_T256A_3pmol:scan:6366:TASC[Carbamidomethyl]GVWDEWSPC[Carbamidomethyl]SVAC[Carbamidomethyl]GK/2

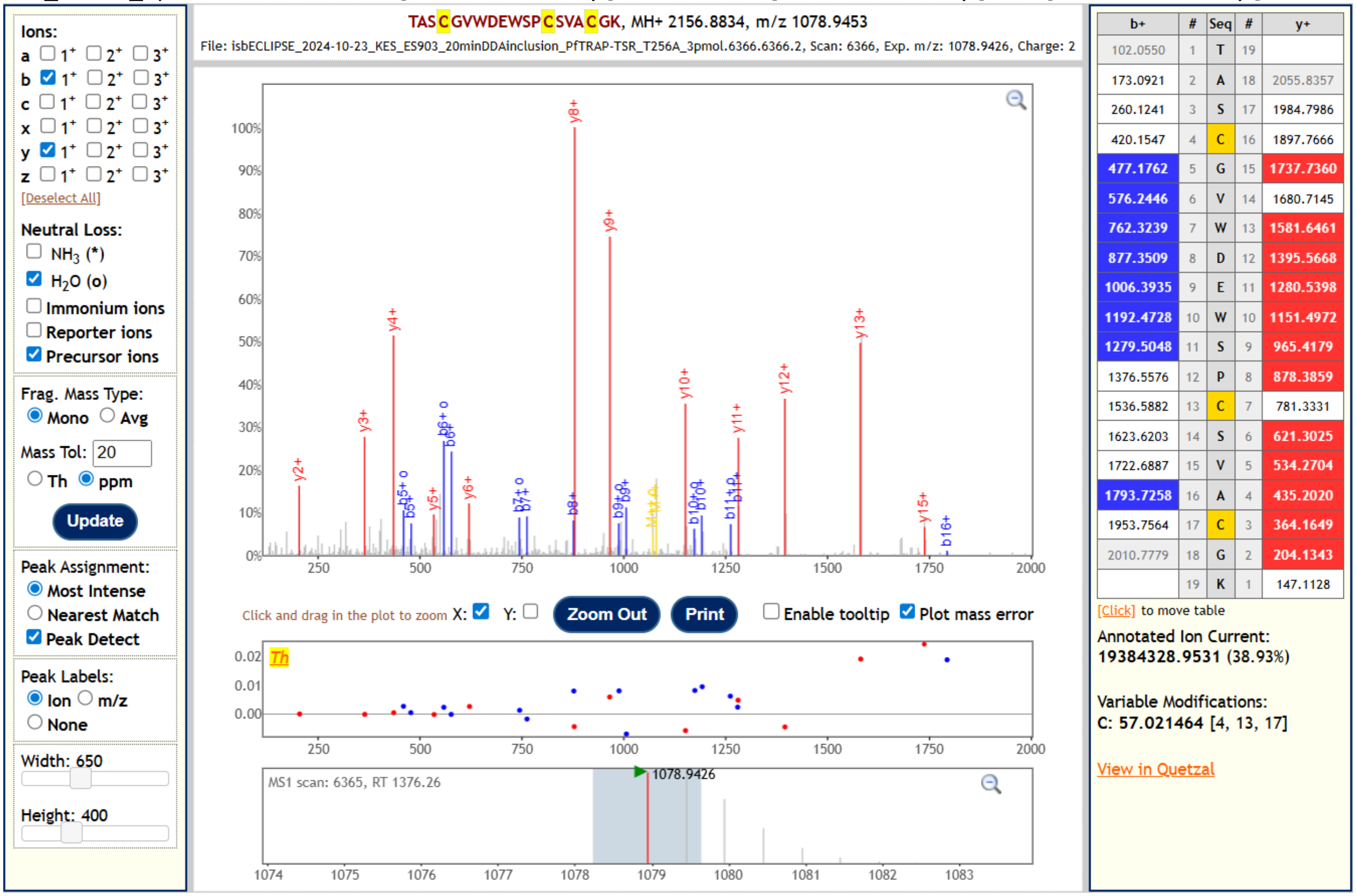


The most abundant glycopeptide in the sample is the unmodified peptide. In contrast to what is observed in the WT and W250F *Pf*TRAP TSR constructs, the peptide without O-Fuc-Glc is detected as a major species rather than as a minor product of neutral loss.

***Pf*TRAP TSR T256A** **TASCGVWDEW[Man]SPCSVACGK**

| xcorr Score | Expect Score | Fragment Ions | Retention Time (sec) | Retention Time (min) | Precursor m/z | Peptide m/z | Neutral Loss |
|---|---|---|---|---|---|---|---|
| 5.739 | 5.07E-17 | 24/36 | 1239.7 | 20.66 | 1159.97 | 1159.97 | 0.00 |

**USI:** mzspec:PXD064887:isbECLIPSE_2024-10-23_KES_ES903_20minDDAinclusion_PfTRAP-TSR_T256A_3pmol:scan:5742:TASC[Carbamidomethyl]GVWDEW[Hex]SPC[Carbamidomethyl]SVAC[Carbamidomethyl]GK/2

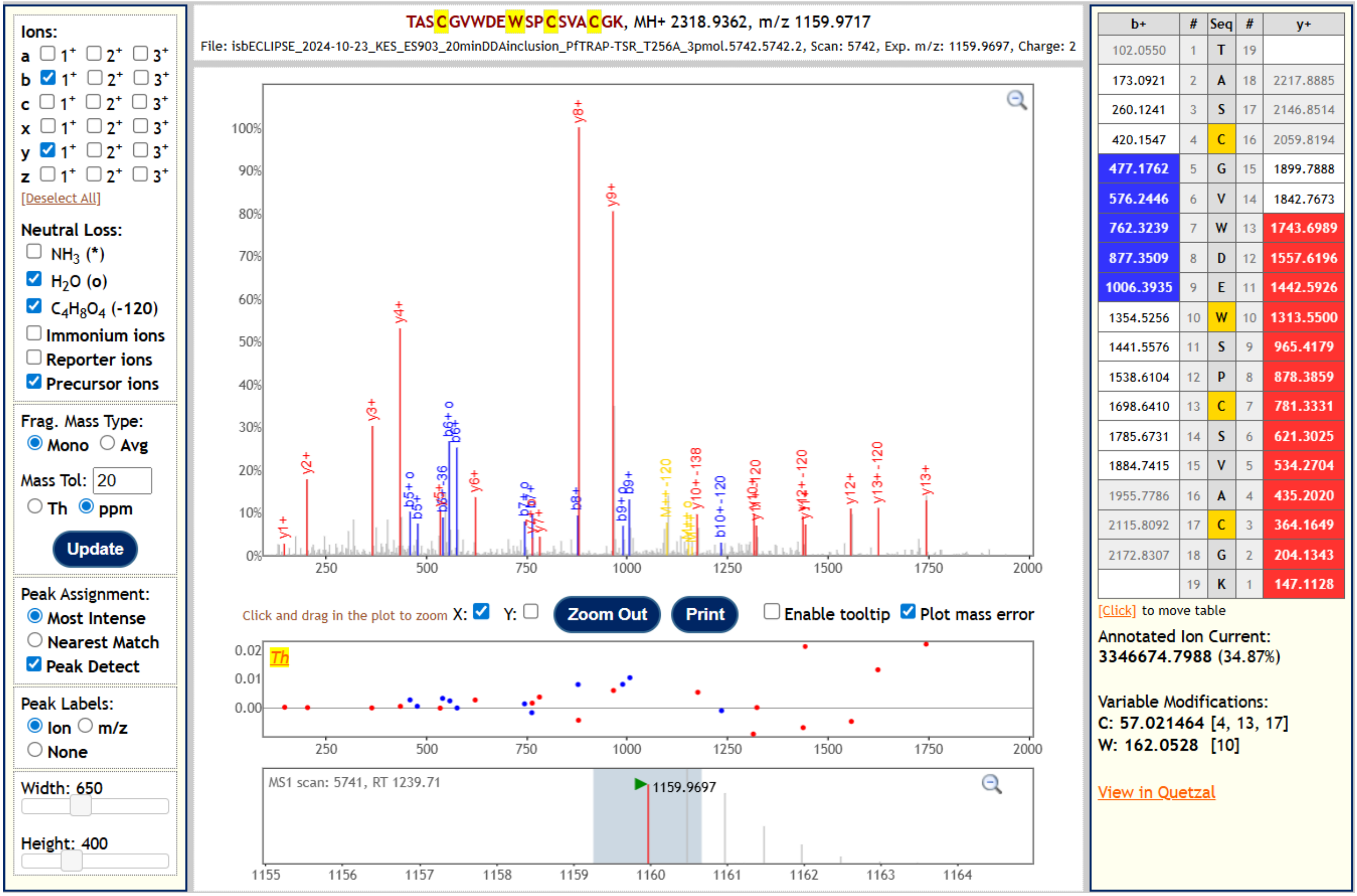


The second most abundant glycoform detected is the peptide with C-Man and without O-fucosylation. Fragment ions identify a Hex (i.e., C-Man) and localize it to Trp[250]. Neutral loss of 120.04 Da from cross-ring cleavage of the C-Man is visible on precursor and fragment ions, further corroborating the presence and localization of the C-Man.

***Pf*TRAP TSR T256A** **TASCGVW[Man]DEW[Man]SPCSVACGK**

| xcorr Score | Expect Score | Fragment Ions | Retention Time (sec) | Retention Time (min) | Precursor m/z | Peptide m/z | Neutral Loss |
|---|---|---|---|---|---|---|---|
| 3.245 | 5.99E-17 | 14/36 | 1143.1 | 19.05 | 1241.00 | 1241.00 | 0.00 |

**USI:** mzspec:PXD064887:isbECLIPSE_2024-10-23_KES_ES903_20minDDAinclusion_PfTRAP-TSR_T256A_3pmol:scan:5294:TASC[Carbamidomethyl]GVW[Hex]DEW[Hex]SPC[Carbamidomethyl]SVAC[Carbamidomethyl]GK/2

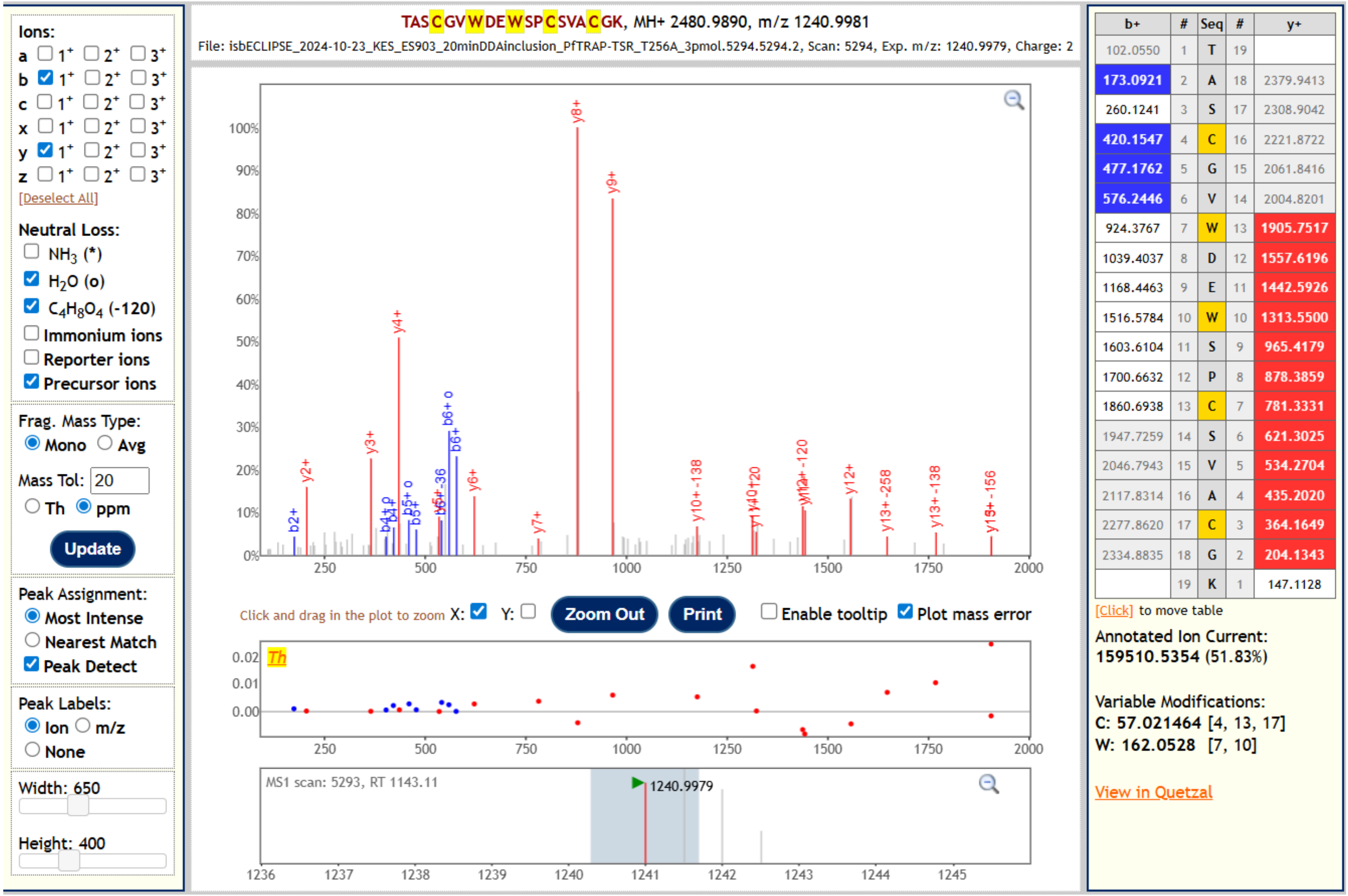


The precursor mass of this peptide corresponds to addition of two Hex moieties to the peptide. Fragment ions and characteristic neutral loss of 120.04 Da from cross-ring cleavage of C-Man confirm the presence of C-Man at Trp$^{247}$ and Trp$^{250}$.
